# The Dorsal Column Nuclei Scale Mechanical Sensitivity in Naive and Neuropathic Pain States

**DOI:** 10.1101/2024.02.20.581208

**Authors:** Aman Upadhyay, Mark A. Gradwell, Thomas J. Vajtay, James Conner, Arnab A. Sanyal, Chloe Azadegan, Komal R. Patel, Joshua K. Thackray, Manon Bohic, Fumiyasu Imai, Simon O. Ogundare, Yutaka Yoshida, Ishmail Abdus-Saboor, Eiman Azim, Victoria E. Abraira

## Abstract

Tactile perception relies on reliable transmission and modulation of low-threshold information as it travels from the periphery to the brain. During pathological conditions, tactile stimuli can aberrantly engage nociceptive pathways leading to the perception of touch as pain, known as mechanical allodynia. Two main drivers of peripheral tactile information, low-threshold mechanoreceptors (LTMRs) and postsynaptic dorsal column neurons (PSDCs), terminate in the brainstem dorsal column nuclei (DCN). Activity within the DRG, spinal cord, and DCN have all been implicated in mediating allodynia, yet the DCN remains understudied at the cellular, circuit, and functional levels compared to the other two. Here, we show that the gracile nucleus (Gr) of the DCN mediates tactile sensitivity for low-threshold stimuli and contributes to mechanical allodynia during neuropathic pain in mice. We found that the Gr contains local inhibitory interneurons in addition to thalamus-projecting neurons, which are differentially innervated by primary afferents and spinal inputs. Functional manipulations of these distinct Gr neuronal populations resulted in bidirectional changes to tactile sensitivity, but did not affect noxious mechanical or thermal sensitivity. During neuropathic pain, silencing Gr projection neurons or activating Gr inhibitory neurons was able to reduce tactile hypersensitivity, and enhancing inhibition was able to ameliorate paw withdrawal signatures of neuropathic pain, like shaking. Collectively, these results suggest that the Gr plays a specific role in mediating hypersensitivity to low-threshold, innocuous mechanical stimuli during neuropathic pain, and that Gr activity contributes to affective, pain-associated phenotypes of mechanical allodynia. Therefore, these brainstem circuits work in tandem with traditional spinal circuits underlying allodynia, resulting in enhanced signaling of tactile stimuli in the brain during neuropathic pain.

## INTRODUCTION

Mechanical allodynia is a common symptom of chronic pain, and is defined by tactile hypersensitivity and pathologically painful responses to innocuous stimuli^1,2^. Treatment of mechanical allodynia is hindered by a lack of complete understanding of underlying mechanisms and circuits. Seeking to address this, a significant effort has been made to investigate regions responsible for early processing of touch and pain in the context of neuropathic pain. This work has resulted in a better understanding of the cellular and circuit mechanisms, most notably in the dorsal root ganglia (DRG) and spinal cord, that contribute to mechanical allodynia. For example, sensitization of nociceptors to low-threshold stimuli^3–5^ or disruption of spinal circuits which normally gate touch^6,7^ allows for innocuous stimuli to engage nociceptive pathways and promote allodynia. Additionally, more upstream tactile and nociceptive pathways such as the parabrachial nucleus^8,9^ and somatosensory cortex^10^ have also been studied in the context of mechanical allodynia. Based on these findings, it is likely that pathological conditions dysregulate tactile information across multiple sites, resulting in aberrant responses to innocuous stimuli. Importantly, transections or nerve block at the level of the DRG and spinal cord are sufficient to block tactile allodynia^11–15^, suggesting that the transmission of tactile information to supraspinal centers is key for the perception of tactile stimuli as painful. Therefore, strategies to investigate circuits contributing to mechanical allodynia may involve looking at early mechanosensory regions which first respond to tactile stimuli, but are not as well characterized as the DRG and spinal cord. Interestingly, two early transmitters of tactile information, the DRG and spinal cord, project to the brainstem dorsal column nuclei complex (DCN).

These nuclei integrate touch in a somatotopic manner and project to the thalamus, where tactile signals are dispersed to relevant cortical regions^16,17^. However, relative to the spinal cord and DRG, the molecular profile, circuit organization, and potential contribution of the DCN to tactile hypersensitivity following nerve injury remain understudied. Interestingly, pharmacological manipulation of the DCN was able to alleviate tactile hypersensitivity during neuropathic pain in rodents^18^. Additionally, dorsal column stimulation has been successful in the management of pain symptoms in humans^19^. However, the underlying circuits mediating the DCN’s contribution to tactile hypersensitivity and allodynia are not well understood. Therefore, the DCN represents a clinically relevant, yet mechanistically understudied, region for the management of neuropathic pain.

Interestingly, activity in the DCN is altered during chronic pain. DCN neurons exhibit more spontaneous activity and afterdischarge during both inflammatory^20^ and neuropathic pain^21,22^ conditions, and display increased recruitment to peripheral nerve stimulation following nerve injury^23,24^. Furthermore, spinal changes associated with neuronal hyperactivity during injury-induced neuropathic pain, such as microglial recruitment^25^ and upregulation of neuropeptides^26,27^, are mirrored in the DCN^28,29^. These results suggest that parallel mechanisms may underlie spinal and DCN dysfunction, and that enhanced DCN activity in response to sensory stimuli may contribute to allodynia following nerve injury. Two main pathways conveying peripheral tactile stimuli to the DCN are collaterals of primary afferents innervating the spinal cord, as well as spinal projection neurons traveling through the dorsal columns (predominantly PSDCs). Both primary afferents and PSDCs have been implicated in mediating phenotypes of neuropathic pain, yet the DCN itself is understudied in this context. In the DRG, Aβ, AD and C-LTMRs are implicated in mediating allodynia during neuropathic pain^30–33^. PSDCs, which integrate LTMR inputs conveying touch, contribute to dorsal column stimulation-induced analgesia^19^.

Therefore, dysfunction of primary afferents and PSDCs, in addition to local immune and neurochemical changes which manifest in the DCN following nerve injury, are uniquely poised to alter DCN activity during pathological conditions. A better understanding of how DCN circuits encode tactile information, and how this encoding is altered following neuropathic injury to promote allodynia, will better inform mechanisms underlying both tactile sensitivity and mechanical allodynia.

Recent evidence suggests that computation within the DCN exceeds what is expected from a simple labeled-line pattern of transmission. In the macaque, cuneate neurons were found to exhibit responses to multiple classes of sensory stimuli and had intricate receptive fields and modality-tuning^34–36^. Recent work in mice has shown that the DCN receives convergent, somatotopically aligned inputs from LTMRs and PSDCs, mediating fine vibrotactile stimuli or high force mechanical stimuli, respectively^37^. This convergence closely resembles response profiles of cortical neurons and suggests that complex sensory processing can occur at earlier sites of tactile signaling than previously thought. In addition, descending cortical projections can influence the “gain” of sensory information by utilizing local inhibition to generate central-surround receptive fields^38–42^. Therefore, inhibition within the DCN represents a powerful mediator of spatial and modality-dependent signals in the DCN that may scale tactile information before it reaches the brain. Recently, DCN inhibition has been suggested to play a role in sensory feedback-mediated grasping during dexterous forelimb behavior^43^. However, the functional implications of local inhibition in the context of tactile sensitivity and mechanical allodynia are unknown. In the spinal cord, loss of inhibition is a key component of central sensitization during mechanical allodynia^44–47^, allowing for tactile information to engage nociceptive pathways^6,48–50^. A more complete understanding of how DCN inhibitory neurons shape tactile responses will not only better inform mechanisms of maintaining tactile sensitivity but will also redefine the anatomical substrates for tactile-evoked manifestations of neuropathic pain.

In this study, we investigated how DCN circuits can shape tactile sensitivity in naive mice as well as mechanical allodynia during neuropathic pain. We focused on the gracile nucleus (Gr) of the DCN, which receives tactile information from the lower body including the hindlimb. We found that VPL-projecting neurons (VPL-PNs) and local inhibitory neurons represent two key neuronal populations in the Gr, which are differentially innervated by LTMRs and PSDCs conveying tactile information. Silencing VPL-PNs reduced tactile sensitivity, while silencing inhibitory neurons induced tactile hypersensitivity and generated neuropathic pain-like paw withdrawal signatures. Tactile-evoked activity in the Gr was found to be increased following neuropathic injury, which was coupled with increased drive from primary afferents onto VPL-PNs. Lastly, activating Gr inhibition during neuropathic pain was able to reduce tactile hypersensitivity and ameliorate pain-related paw withdrawal signatures. These results demonstrate that the Gr plays a modality and intensity-specific role in the modulation of tactile signals which contributes to mechanical allodynia during neuropathic pain.

## RESULTS

### Ascending tactile pathways and descending cortical projections converge in the gracile nucleus

In the spinal cord, characterization of distinct projection neuron populations have revealed modality-specific outputs^37,51–55^, which are dynamically modulated by molecularly defined excitatory and inhibitory interneurons^44,48,56–60^. This interplay between spinal interneurons and projection neurons is what gives rise to ascending tactile signals. Additionally, modulation of either population can alter the transmission of tactile information^52,61,62^, even allowing for tactile signals to elicit pain^6,63^ or itch^64–66^. Despite being functionally implicated in tactile sensation, similar characterizations for DCN circuits in the context of somatosensation are nascent. The DCN are a key part of the medial lemniscal pathway, which integrates primary afferent and spinal inputs and projects to the ventral posterolateral thalamus (VPL) **(Figure 1A)**. Within the DCN, subnuclei are defined by somatotopy related to incoming sensory information^17^. The gracile nucleus (Gr) receives tactile information via primary afferents and spinal projections from below spinal segment T6, while the neighboring cuneate nucleus (Cu) receives similar projections from above spinal segment T6^17^. We focused our analyses on the Gr which receives information from the lower body, including the hindlimb where many rodent sensory assays and pain models have been validated^67^. It is well established that the DCN receives tactile information via a subset of LTMRs as well as spinal projections conveying touch, notably PSDCs^17,37,68^. To characterize the Gr, cutaneous hindlimb afferents were identified via injection of cholera toxin beta subunit (CTb) into the hindpaw, revealing dense projections within the ipsilateral Gr “core” **(Figure 1B)**. Comparatively, primary afferents labeled using a mouse genetics strategy (*Advillin^Cre^*;*Rosa26^LSL-SynGFP^*) were observed both in the Gr core and a surrounding “shell” region **(Figure 1C)**, suggesting that other sensory regions, such as the hindlimb, back hairy skin, or viscera may contribute to primary afferent innervation of the shell, while the hindpaw targets the core. To confirm the identity of primary afferents innervating the Gr, we used genetic intersections to visualize synaptic terminals of DRG afferents subtypes within the Gr. We observed dense projections from Aβ RA-LTMRs **(Figure S1A)** and Aβ Field-LTMRs **(Figure S1B)**, limited projections from Aβ SA-LTMRs **(Figure S1C)**, AD-LTMRs **(Figure S1D)**, and proprioceptors **(Figure S1H)**, and very few inputs from C-LTMRs **(Figure S1E)** and nociceptors **(Figure S1F,G)**. These results were largely consistent with previous literature identifying Aβ-LTMRs as the dominant source of primary afferent input into the Gr^68–70^. To investigate PSDC innervation of the Gr, we generated a genetic intersection where a large population of dorsal horn spinal neurons were labeled (*Cdx2^Cre^*;*Lbx1^FlpO^*;*Rosa26^LSL-FSF-TdTomato^*) **(Figure S1I)**. Importantly, labeled neurons were not present at or above the level of the DCN, indicating this intersection is well restricted to the spinal cord. Next, we generated *Cdx2^Cre^*;*Lbx1^FlpO^*;*Rosa26^LSL-FSF-Synaptophysin-GFP^* mice to visualize terminals of spinal projections in the Gr **(Figure 1D)**, which are known to predominantly originate from PSDCs. Similar to primary afferents, spinal inputs were found in both the Gr core and shell. Collectively, these results suggest that primary afferent and spinal projections target both Gr core and shell regions, and are thus poised to transmit ascending sensory information to similar regions of the Gr.

**Figure 1.**
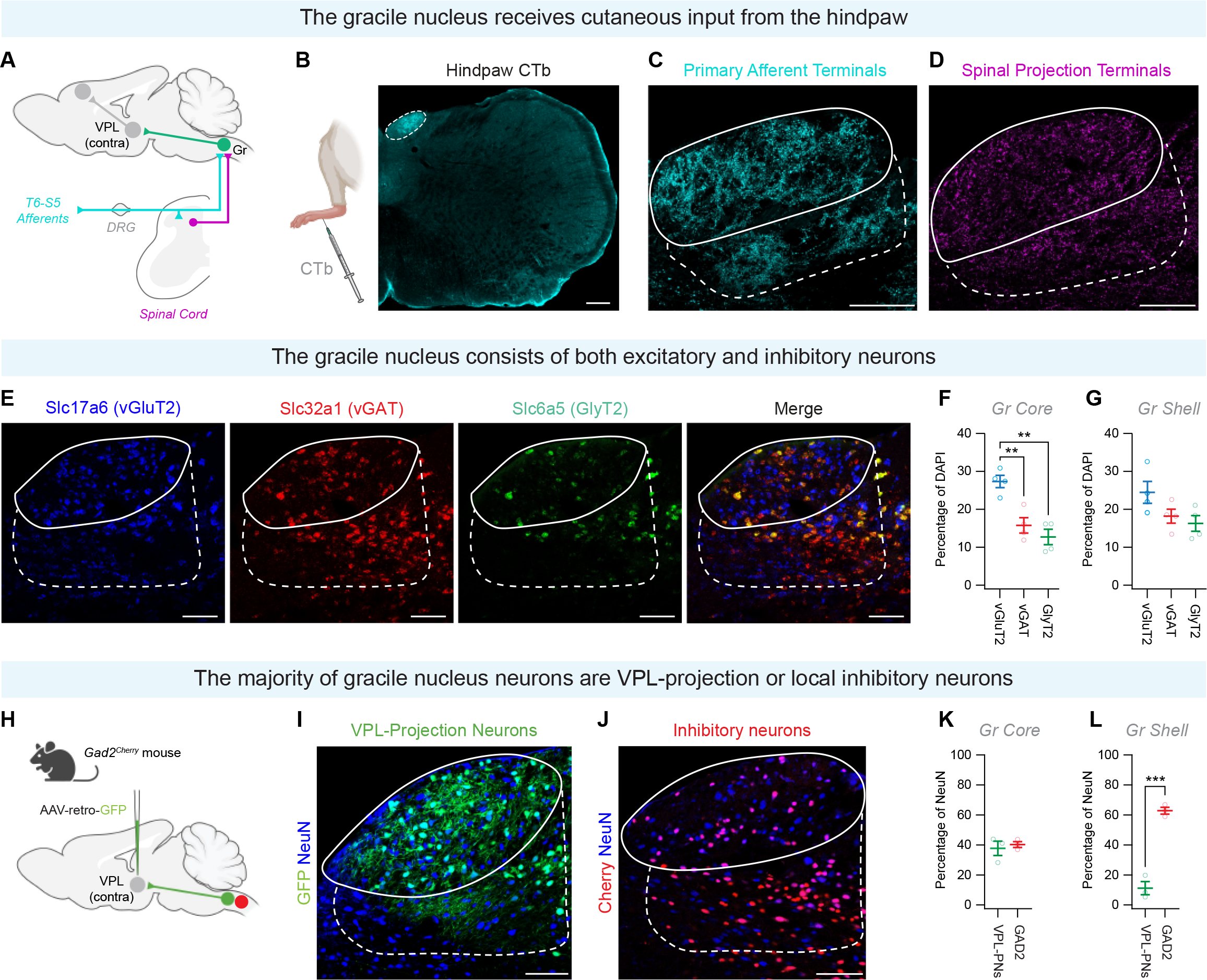
Ascending DRG and spinal pathways and descending cortical projections converge in the gracile nucleus A) The dorsal column nuclei complex (DCN) is located ventral to the cerebellum and consists of subnuclei organized by somatotopy. The gracile nucleus (Gr, green) is located medially and receives sensory inputs from the lower body from DRG afferents (cyan) and spinal projections (magenta). One of the main projections of the Gr is to the contralateral ventroposterolateral thalamus (VPL), which projects to other sensory regions such as the primary somatosensory cortex. B) Injection of cholera toxin beta subunit (CTb, cyan) into the hindpaw glabrous skin labeled hindlimb afferents targeting the Gr (white dashed circle). Scale bar = 200 um. C) *Advillin^Cre^*;*Rosa26^LSL-Synaptophysin-GFP^* mice were generated to visualize synaptic terminals of primary afferents (cyan) in the Gr core (solid white line) and shell (dashed white line). Scale bar = 100 um. D) *Cdx2^Cre^*;*Lbx1^FlpO^*;*Rosa26^LSL-FSF-Synaptophysin-GFP^*mice were generated to visualize terminals of spinal projections (magenta) in the Gr core (solid white line) and shell (dashed white line). Scale bar = 100 um. E) *In situ* hybridization to identify distribution of excitatory (vGluT2, blue) and inhibitory (vGAT, red, GlyT2, green) neurons in the Gr core (solid white line) and shell (dashed white line). Scale bar = 100 um. F,G) Quantification of excitatory and inhibitory neuronal markers in the Gr core (F) and shell (G). H) Retrograde injection of a GFP virus into the VPL was used to identify VPL-projecting neurons (VPL-PNs, green), while *Gad2^Cherry^*mice were used to fluorescently label inhibitory neurons (red). I,J) Representative images of Gr VPL-PNs (I)(green), inhibitory neurons (J)(red), and neurons immunolabeled using an antibody against NeuN (blue). Scale bar = 100 um. K,L) Quantification of VPL-PNs and inhibitory neurons in the Gr core (H) and shell (I) normalized to total NeuN. Data are reported as mean values ± SEM. *p ≤ 0.05, ∗∗p ≤ 0.01 , ∗∗∗p ≤ 0.001, ∗∗∗∗p ≤ 0.0001. For further details on genetic crosses and statistical tests, see STAR Methods.

In addition to ascending tactile pathways, which confer the ability to respond to peripheral stimuli, the DCN circuit activity is also heavily influenced by descending cortical projections. Cortical projections are hypothesized to mainly exert an inhibitory role, such as the suppression of sensory information during movement^42^, and have been shown to suppress Cu responses to tactile stimuli^43^. Mapping in the Cu has also revealed that the primary somatosensory cortex targets the Cu core while the primary motor cortex and rostral sensorimotor cortex targets the Cu shell^43^. However, less is known about their distribution and role in the Gr. To investigate cortical projections to the Gr, we first generated *Emx1^Cre^*;*Rosa26^LSL-Synaptophysin-TdTomato^*(Ai34) mice, where cortical afferent terminals were fluorescently labeled **(Figure S2A)**. Within the DCN, these terminals were found to target both Gr core and shell regions **(Figure S2B)**. Given the bias of sensory over motor cortical regions projecting to Gr regions innervated by cutaneous afferents^71^, we next investigated the origin of these cortical projections. We injected a Cre-dependent virus into the primary somatosensory cortex of *Rbp4^Cre^*mice, where Cre expression was restricted to layer 5 pyramidal cells **(Figure S2C)**. These injections were focused either to M1, S1-forelimb (S1-FL) or S1-hindlimb (S1-HL) **(Figure S2D-F)**. M1 injections resulted in very sparse labeling in the Gr **(Figure S2D)**. S1-FL injections did not target the Gr, and instead labeled the nearby Cu **(Figure S2E)**. Conversely, S1-HL injections resulted in labeling in the Gr core and shell, but not the Cu **(Figure S2F).** Therefore, the predominant somatosensory cortex projections to the Gr originate from S1-HL, aligning anatomically with somatotopically organized ascending input. Projections from the primary somatosensory cortex, which exhibits hyperexcitability and neuroplasticity during chronic pain^72–74^, represent a relevant source for descending modulation of Gr activity^39,40,42^. Therefore, altered S1 signaling to the Gr during pathological conditions^75–78^ may disrupt tactile signaling, allowing for aberrant tactile information to reach the thalamus and contribute to pain phenotypes^79,80^.

### The cytoarchitecture of the gracile nucleus is comprised of VPL-projecting neurons and local inhibitory interneurons

Within the DCN, complex pre- and postsynaptic mechanisms drive both feed-forward and feedback-driven inhibition^37,43,81–84^, suggesting inhibitory networks modulate the activity of Gr projection neurons. Therefore, we mapped the cellular landscape of the DCN relative to ascending sensory inputs. To characterize distributions of excitatory and inhibitory populations in the Gr, we used *in situ* hybridization to label vGluT2+ excitatory neurons and vGAT+ or GlyT2+ inhibitory neurons **(Figure 1E)**. The Gr core contained a relatively higher number of excitatory neurons compared to inhibitory neurons **(Figure 1F)**, while the shell contained a more even distribution of both populations **(Figure 1G)**. Of the many DCN outputs^17^, VPL-projecting neurons (VPL-PNs) are reported to be the most prominent cell type in the DCN^68^, suggesting they play a major role in DCN function. To characterize VPL-PNs in the context of local inhibition, we utilized viral retrograde tracing from the thalamus **(Figure 1H)** to fluorescently label VPL-PNs **(Figure 1I)**, and used a genetic mouse line (*Gad2^mCherry^*), to visualize inhibitory neurons **(Figure 1J)**. We observed that inhibitory neurons were intermingled with VPL-PNs in the core **(Figure 1K)**, but were densely populated in the shell region where VPL-PNs were sparse **(Figure 1L)**. Therefore, the Gr consists of a mixed core containing excitatory VPL-PNs and inhibitory interneurons, surrounded by an inhibitory shell.

Juxtacellular recordings in the Gr suggest that VPL-PNs exhibit spontaneous activity and cannot entrain to high frequency stimulation, while Gr inhibitory neurons do not exhibit spontaneous activity but can entrain to high frequencies^37^. These results suggest differences in Gr neuron properties which may underlie their responsiveness to tactile stimuli. To build upon these findings, we performed slice electrophysiology to investigate the intrinsic properties of both Gr VPL-PNs and inhibitory neurons. VPL-PNs were identified using retrograde viral tracing, while inhibitory neurons were labeled using a *Gad2^Cherry^*mouse **(Figure 2A)**. We found that VPL-PNs are more than twice as likely to exhibit spontaneous action potential (sAP) discharge than inhibitory neurons **(Figure 2B,C)**. Of the spontaneously active neurons, VPL-PNs exhibited a higher sAP frequency than inhibitory neurons **(Figure 2D)**, indicating that VPL-PNs are more active than inhibitory neurons and are a likely source of sustained excitation to their target region, the VPL. To further examine the intrinsic excitability of these two populations, we examined their responsiveness to depolarizing current injection **(Figure 2E)**. We found that VPL-PNs fire more action potentials than inhibitory neurons in response to small depolarizing current injection steps **(Figure 2F)**. This increased excitability in VPL-PNs was reflected by a lower rheobase **(Figure 2G)**, and a more hyperpolarized action potential threshold **(Figure 2H)**. Together, these data indicate that Gr VPL-PNs are a source of ongoing excitatory input to the VPL. In contrast, Gr inhibitory neurons are less likely to be spontaneously active and, therefore, their activity is likely to be more reliant upon excitatory input from sources such as primary afferents or PSDCs. Interestingly, we also show that Gr VPL-PNs exhibit more excitable properties than Gr inhibitory neurons. This suggests that in response to excitatory input Gr VPL-PNs are more likely to be recruited and excitement of Gr VPL-PNs would precede the activation of Gr inhibitory neurons. If Gr VPL-PNs and Gr inhibitory neurons are recruited by the same source, these properties would support the transduction of sensory information through the Gr and to the VPL, before its modulation by local inhibitory neurons.

**Figure 2.**
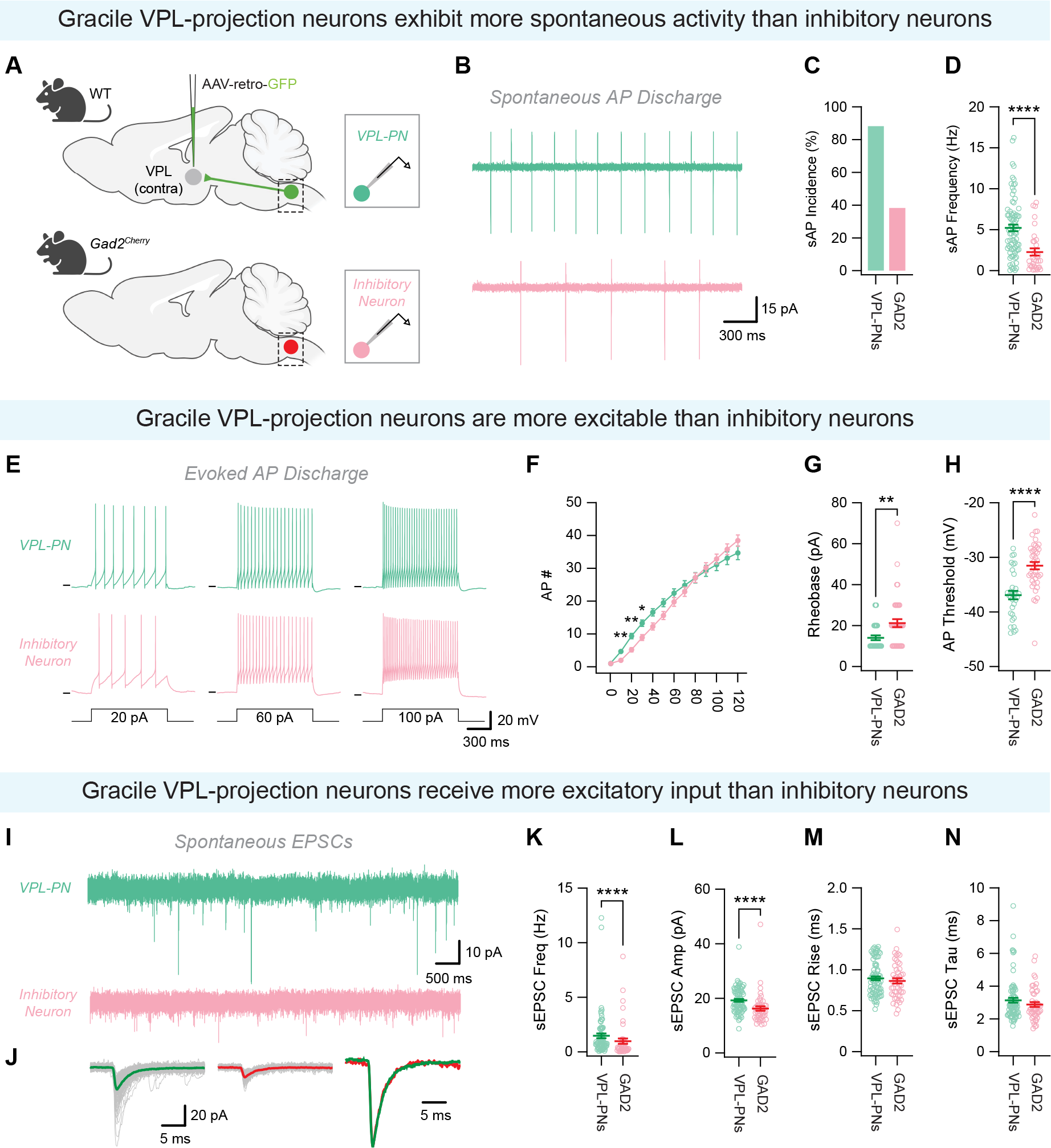
Gr VPL-PNs and inhibitory neurons exhibit different electrophysiological properties and excitatory drive. A) Strategy for identification of VPL-PNs and inhibitory neurons for slice electrophysiology. B) Cell-attached voltage clamp recordings showing spontaneous AP discharge in VPL-PN (green) and inhibitory neuron (red). C) Percentage of neurons exhibiting spontaneous AP discharge. D) Quantification of sAP frequency. E) Whole-cell current clamp recording (-60 mV) from VPL-PN (green) and inhibitory neuron (red) in response to depolarizing current injection. F) Quantification of AP number in response to depolarizing current injection. G) Quantification of AP rheobase. H) Quantification of AP threshold. I) Whole-cell voltage clamp recordings (-70 mV) of sEPSCs from VPL-PN (green) and inhibitory neuron (red). J) Captured sEPSCs from representative cells shown in (I). Right trace shows amplitude normalized overlay. K) Quantification of sEPSC frequency. L) Quantification of sEPSC amplitude. M) Quantification of sEPSC rise time. N) Quantification of sEPSC decay time. Data are reported as mean values ± SEM. *p ≤ 0.05, ∗∗p ≤ 0.01 , ∗∗∗p ≤ 0.001, ∗∗∗∗p ≤ 0.0001. For further details on genetic crosses and statistical tests, see STAR Methods.

### Gr VPL-PNs and inhibitory neurons are differentially innervated by primary afferent and spinal inputs

Recent work in the Gr suggests that somatotopically aligned primary afferents and PSDCs converge onto individual VPL-PNs^37^, suggesting these two pathways can merge to engage DCN outputs to the thalamus. However, the relative strengths of these two pathways onto VPL-PNs, as well as their influence on local inhibitory neurons, is not well characterized. To better understand how ascending tactile pathways engage distinct DCN circuits, we utilized slice electrophysiology to investigate primary afferent and spinal inputs onto Gr VPL-PNs or inhibitory neurons. To characterize primary afferent input onto Gr neurons, we generated *Advillin^Cre^*;*Rosa26^LSL-ChR2/eYFP^*(Ai32) mice, where channelrhodopsin was restricted to primary afferent terminals, and injected a retrograde-mCherry virus into the VPL to label VPL-PNs, or generated *Advillin^Cre^*;Ai32;*Gad2^Cherry^* mice, where inhibitory neurons were fluorescently labeled **(Figure 3A)**. Similarly for PSDCs, we generated *Cdx2^Cre^*;*Lbx1^FlpO^*;*Rosa26^CatCh/eYFP^*(Ai80) mice to expression channelrhodopsin in spinal projections, and injected a retrograde-mCherry virus into the VPL to label VPL-PNs, or generated *Cdx2^Cre^*;*Lbx1^FlpO^*;Ai80;*Gad2^Cherry^* mice, where inhibitory neurons were fluorescently labeled **(Figure 3A)**. Next, we recorded from Gr VPL-PNs or inhibitory neurons and activated either primary afferent or spinal projection terminals. Generally, optical stimulation of primary afferent terminals within the DCN evoked monosynaptic EPSCs with a greater amplitude and increased incidence when compared to stimulation of spinal projections within the DCN **(Figure 3B-D)**.

**Figure 3.**
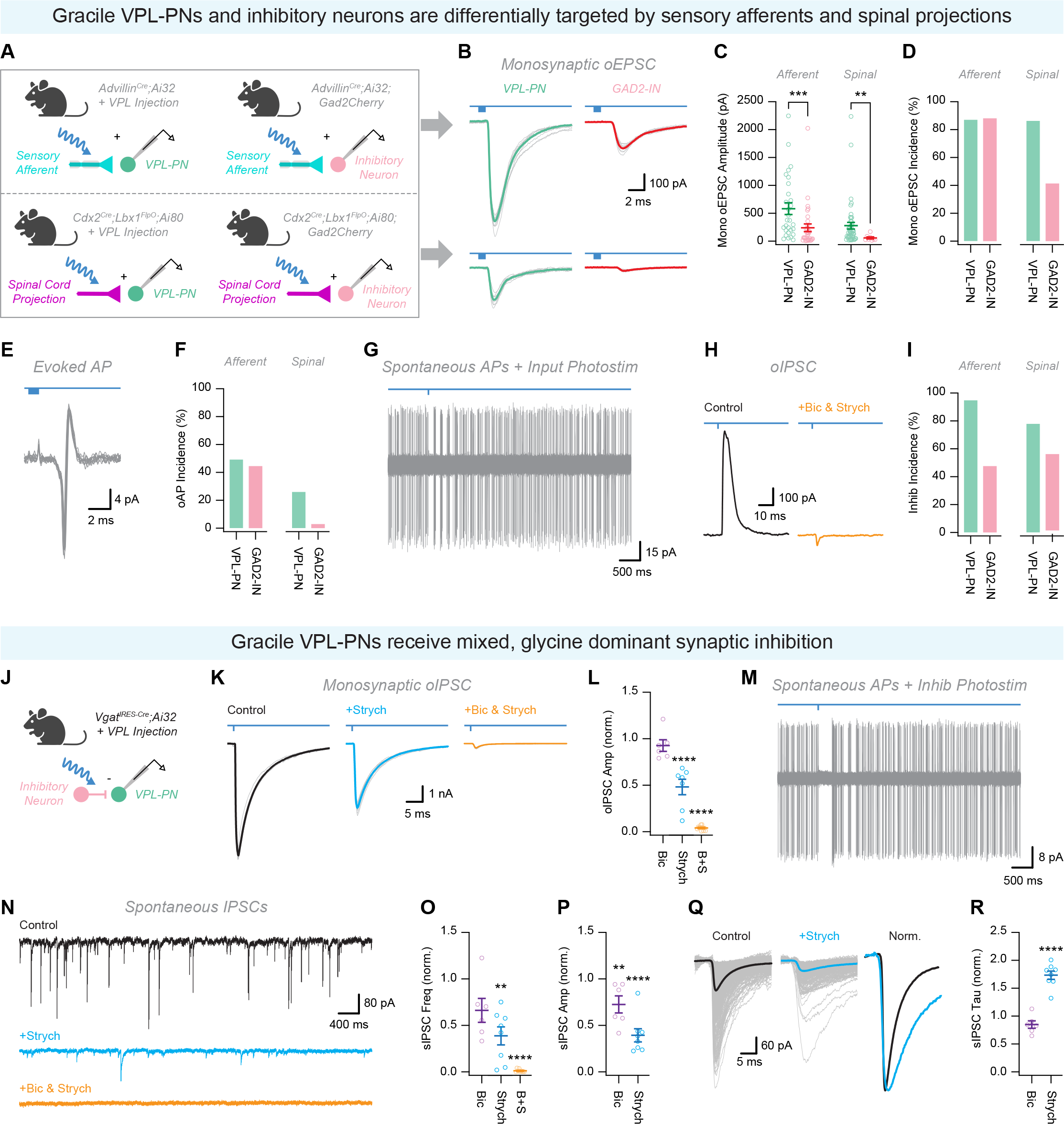
Ascending primary afferent and spinal inputs differentially innervate Gr VPL-PNs and inhibitory neurons and evoke feedforward inhibition A) Strategies to optogenetically activate primary afferent or spinal inputs onto Gr VPL-PNs or inhibitory neurons. B) Whole-cell voltage clamp recordings (-70 mV) of optically-evoked EPSCs (oEPSCs) from VPL-PN (green) and inhibitory neuron (red) in response to primary afferent (top) and spinal projection (bottom) photostimulation. 10 consecutive sweeps with average overlaid. C) Quantification of monosynaptic oEPSC amplitude. D) Percentage of neurons exhibiting monosynaptic oEPSCs. E) Cell-attached voltage clamp recording showing optically evoked AP discharge (sAP). 10 consecutive sweeps overlaid. F) Percentage of neurons exhibiting sAP discharge. G) Cell-attached voltage clamp recording showing the impact of optogenetic activation of ascending inputs on spontaneous AP discharge. 10 consecutive sweeps overlaid. H) Whole-cell voltage clamp recordings (-20 mV) of optically-evoked IPSCs (oIPSCs) in control (black) and following application of bicuculline (10 µM) and strychnine (1 µM; orange) in response to optogenetic activation of ascending inputs. 10 consecutive sweeps with average overlaid. I) Percentage of neurons exhibiting oIPSCs. J) Strategy to optogenetically activate inhibitory inputs onto Gr VPL-PNs. K) Whole-cell voltage clamp recordings (-70 mV) of optically-evoked IPSCs (oIPSCs) in control (black) and following sequential application of strychnine (1 µM; blue) and bicuculine (10 µM; orange) in response to optogenetic activation of inhibitory neurons. 10 consecutive sweeps with average overlaid. L) Normalized impact of bicuculine (10 µM), strychnine (1 µM), and bicuculine+strychnine application on oIPSC amplitude. M) Cell-attached voltage clamp recording showing the impact of optogenetic activation of inhibitory inputs on spontaneous AP discharge. 10 consecutive sweeps overlaid. N) Whole-cell voltage clamp recordings (-70 mV) of sIPSCs from VPL-PN in control (black) and and following sequential application of strychnine (1 µM; blue) and bicuculine (10 µM; orange). O) Normalized impact of bicuculine (10 µM), strychnine (1 µM), and bicuculine+strychnine application on sIPSC frequency. P) Normalized impact of bicuculine (10 µM), strychnine (1 µM), and bicuculine+strychnine application on sIPSC amplitude. Q) Captured sIPSCs from representative cells shown in (N). Right trace shows amplitude normalized overlay. R) Normalized impact of bicuculine (10 µM), strychnine (1 µM), and bicuculine+strychnine application on sIPSC decay.

Correspondingly, activation of primary afferent terminals was more likely than spinal projections to evoke action potential discharge in VPL-PNs and inhibitory neurons **(Figure 3E-F)**. When comparing inputs from primary afferents or spinal projections to VPL-PNs and inhibitory neurons, we found VPL-PNs received larger inputs following both primary afferent and spinal projection stimulation **(Figure 3C)**. Additionally, we often observed a pause in sAP discharge following primary afferent and spinal projection photostimulation, suggesting a feedforward inhibitory circuit **(Figure 3G)**. In-turn, when cells were held at -20 mV, we also observed longer latency bicuculline/strychine inhibitory currents in response to primary afferent and spinal projection photostimulation **(Figure 3H)**. Interestingly, the incidence of these inhibitory currents was substantially greater in the VPL-PNs than inhibitory neurons, suggesting that inhibitory neurons preferentially target the Gr core **(Figure 3I)**. Together, these data suggest a circuit organization where primary afferent inputs play a more substantial role in activating DCN circuits than spinal inputs, and VPL-PNs receive stronger inputs than Gr inhibitory neurons. Further, inhibitory neurons appear to preferentially target the Gr core, where the great majority of VPL-PNs and small portion of inhibitory neurons reside. This arrangement further supports the transmission of sensory information through the Gr and to the VPL by VPL-PNs, particularly driven by primary afferent input, and highlights a potentially interesting role for feedforward inhibitory regulation of VPL-PNs activity.

After identifying a prominent feedforward inhibition of VPL-PNs, we sought to further examine this inhibitory circuit. To determine the functional connectivity between inhibitory neurons and VPL-PNs, we generated *Vgat^IRES-Cre^*;Ai32 mice, where channelrhodopsin was restricted to inhibitory neurons, and injected a retrograde-mCherry virus into the VPL to label VPL-PNs **(Figure 3J)**. Optogenetic activation of inhibitory neurons evoked substantial monosynaptic inhibitory postsynaptic currents (oIPSCs) in all VPL-PN recordings. Notably, we observed optically-evoked monosynaptic inhibitory postsynaptic currents (oIPSCs) in all VPL-PN recordings **(Figure 3K)**. Application of strychnine and bicuculline significantly attenuated oIPSCs, with strychnine having a more pronounced effect on oIPSC amplitude, indicating predominant glycine-mediated inhibition **(Figure 3K-L)**. Interestingly, both blockers had a larger impact of oIPSC amplitude when applied sequentially following application of the other. This interaction suggests a potential role for local disinhibition (pre- or postsynaptic) capable of modulating direct inhibition of VPL-PNs. Despite this potential disinhibitory circuit, optogenetic activation of inhibitory neurons robustly inhibited VPL-PN sAP discharge, highlighting the potential of local inhibitory circuits to reduce VPL-PN output **(Figure 3M)**. Additionally, VPL-PNs received robust spontaneous inhibitory input, which was abolished by the application of strychnine and bicuculline **(Figure 3N)**. Consistent with optogenetically evoked inhibition, strychnine had a more pronounced impact on sIPSC frequency **(Figure 3O)**, amplitude **(Figure 3P)**, and kinetics **(Figure 3Q-R)**, indicating the dominance of glycine-mediated inhibition.

Encoding of sensory stimuli in the DCN is further modulated by presynaptic inhibition^85^, which is suggested to play a role in sensory-induced surround inhibition^84,86^. In the spinal cord dorsal horn, primary afferents are more likely to be apposed to presynaptic inhibitory contacts than cortical afferents^61^. However, the organization of presynaptic inhibition relative to DCN afferent inputs is not well characterized. Similar to spinal circuits, we hypothesized that presynaptic inhibition in the Gr would favor ascending over descending pathways, allowing for better cortical control of tactile information before it reaches the DCN. To investigate presynaptic contacts onto primary afferent and cortical terminals in the Gr, we generated either *Advillin^Cre^*;Ai34 mice to visualize primary afferent terminals **(Figure S3A)**, or *Emx1^Cre^*;Ai34 mice to visualize cortical terminals **(Figure S3B)** and utilized immunolabeling for vGAT to visualize inhibitory contacts^6,61,87^. In the Gr, primary afferent terminals were more likely to receive axoaxonic contacts than cortical afferent terminals **(Figure S3C)**, and had more associated axoaxonic contacts per afferent terminal **(Figure S3D)**, consistent with observations in the spinal cord dorsal horn^61^. Lastly, we investigated whether presynaptic inhibition in the DCN could be a conserved mechanism across species. We performed immunostaining in the macaque Gr for vGLUT1 (presumed primary afferent terminals) and vGAT (putative inhibitory contacts). We found evidence of putative axoaxonic contacts in the macaque Gr **(Figure S3E)**, suggesting this mechanism is conserved in the primate. Therefore, presynaptic inhibition in the Gr displays a similar preference towards primary afferents over cortical afferents as in the spinal cord, and may undergo a similar dysfunction as spinal inhibition following nerve injury^44,48^ to contribute to altered tactile signaling in the DCN.

In summary, these results reveal that ascending somatosensory pathways drive both VPL-PNs and inhibitory neurons, with primary afferent terminals playing a more substantial role. Moreover, a pronounced feedforward inhibitory circuitry influences VPL-PN activity, underscoring the intricate balance between excitatory and inhibitory circuits within the Gr. These circuits are poised to modulate ascending tactile signals before they reach the thalamus, both in normal tactile processing as well as during neuropathic pain.

### Manipulation of Gr VPL-PNs and inhibitory neurons bidirectionally scales tactile sensitivity and intensity of withdrawal response to innocuous tactile stimuli

Broad manipulations, such as dorsal column lesions or DCN lesions, result in deficits to mechanical sensation including two-point sensitivity and grasp-related tactile discrimination^88–91^, as well as reduction of mechanical allodynia^18,92^. However, dorsal column lesions do not fully eliminate the ability to sense tactile stimuli, which is suggested to have a redundant pathway through spinothalamic neurons traveling via the anterolateral tract^88,93,94^. Therefore, the extent to which the DCN contributes to tactile perception, and which specific stimuli rely on this pathway for sensory perception, is not well characterized. Lesions of the dorsal column or DCN influence a combination of distinct excitatory and inhibitory subpopulations which may differentially contribute to tactile sensitivity. In the spinal cord, inhibition plays a key role in setting pain thresholds^57^, and prevents innocuous information from engaging nociceptive pathways^49,95^. However, spinal mechanisms such as gating of tactile inputs from engaging nociceptive projection neurons are likely not conserved in the DCN given the absence of any major DCN outputs to nociceptive regions. Interestingly, dorsal horn circuits exhibit intensity coding, where the strength of afferent activation determines whether a stimulus is interpreted as itch or pain^96^. Therefore, a similar mechanism of intensity coding in the DCN may underlie functional contributions of the DCN to tactile-evoked phenotypes of neuropathic pain.

Given that both VPL-PNs and inhibitory neurons are innervated by primary afferents and spinal projections **(Figure 3B-F)**, they may collectively contribute to DCN outputs which influence tactile perception. Therefore, functional changes to either population are likely to alter tactile responses. Therefore, we investigated responses of mice to sensory stimuli applied to the hindpaw while manipulating either VPL-PNs or inhibitory neurons. To manipulate VPL-PN activity, we injected a retrograde Cre virus into the VPL and a Cre-dependent hM4Di-DREADD virus into the Gr **(Figure 4A)** to express hM4Di in VPL-PNs **(Figure 4B)**. Administration of CNO resulted in decreased tactile sensitivity to lower and mid-force von Frey filaments (0.4-1.0g), while responses to higher force filaments (1.4-4g) were unaffected **(Figure 4C)**. Additionally, CNO administration reduced sensitivity to dynamic brush stimulation **(Figure 4D)**. Interestingly, CNO administration did not affect sensitivity to a noxious mechanical pinprick stimulus **(Figure S4B)** or radiant thermal sensitivity using the Hargreaves assay **(Figure S4C)**. Collectively, these results suggest that VPL-PNs mediate responses to low-threshold, innocuous stimuli, but not to noxious mechanical or thermal stimuli.

**Figure 4.**
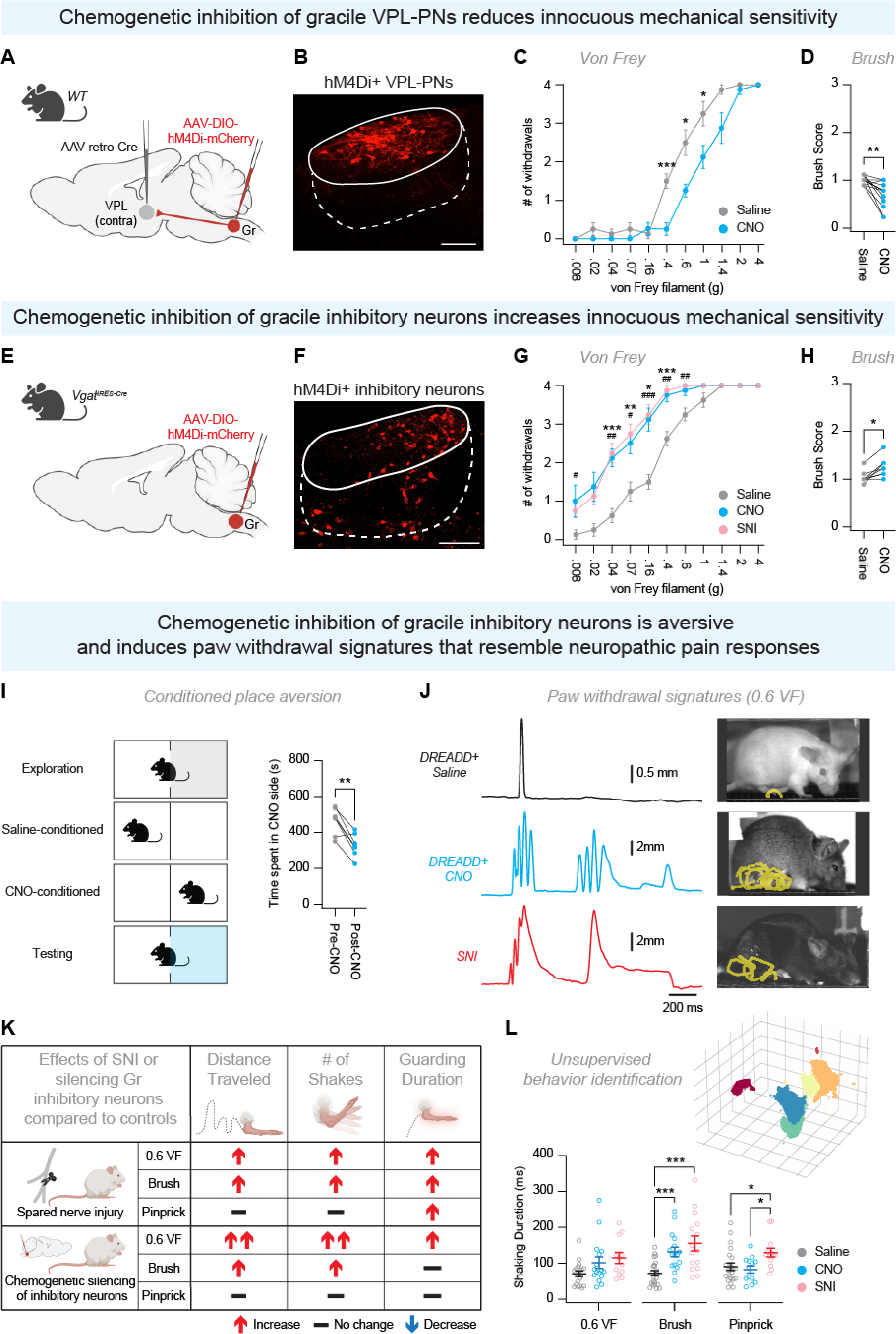
Manipulation of Gr VPL-PNs and inhibitory neurons bidirectionally scales tactile sensitivity and intensity of withdrawal response to innocuous tactile stimuli A,B) To silence Gr VPL-PNs, we injected a retrograde-Cre virus into the VPL, and injected a Cre-dependent inhibitory hM4Di DREADD virus (red) into the Gr. VPL-PNs were silenced by administering CNO, while saline was used as a control. Scale bar = 100 um. C) Punctate mechanical sensitivity in response to von Frey stimulation in mice given CNO (blue) or saline (gray). D) Dynamic mechanical sensitivity in response to a brush stimulation in mice given CNO (blue) or saline (gray). E,F) To silence Gr inhibitory neurons, we injected a Cre-dependent inhibitory hM4Di DREADD virus (red) into the Gr of *Vgat^IRES-Cre^* mice. Gr inhibitory neurons were silenced by administering CNO, while saline was used as a control. Scale bar = 100 um. G) Punctate mechanical sensitivity in response to von Frey stimulation in mice given CNO (blue) or saline (gray), or mice with an SNI model of neuropathic pain (red). *=CNO vs. Saline, ^#^=SNI vs. Saline. H) Dynamic mechanical sensitivity in response to a brush stimulation in mice given CNO (blue) or saline (gray). I) Conditioned place preference assay to assess appetitive/aversive nature of CNO administration. Time spent was measured in the CNO-paired side pre-conditioning (gray) and post-conditioning (blue). J) DeepLabCut was used to track the coordinates of the hindpaw in response to a mechanical stimulus. Representative traces of metatarsophalangeal joint (MTP) y-coordinates during withdrawal responses to a 0.6g von Frey filament for control mice (black), mice with chemogenetic silencing of Gr inhibitory neurons (blue), and SNI mice (red). K) Pain Assessment at Withdrawal Speeds (PAWS) software was used to extract features of the paw withdrawal response including distance traveled, number of shakes, and guarding duration. Further explanation of how this software extracts relevant features can be found in Jones et al., 2020^102^. Features for SNI mice and mice where Gr inhibitory neurons were silenced were compared to saline controls. ↑ represents an increase compared to control mice, ↓ represents a decrease compared to control mice, and - represents no change. L) First three dimensions of UMAP (uniform manifold approximation and projection) embedding, showing in low-dimensional space a projection of extracted behavioral features. Colors represent behavioral clusters identified by HDBSCAN (hierarchical density-based spatial clustering of applications with noise), including a cluster corresponding to shaking behavior (turquoise). Graph shows comparison of normalized duration of shaking behavior in saline, CNO, and SNI mice. Data are reported as mean values ± SEM. *p ≤ 0.05, ∗∗p ≤ 0.01 , ∗∗∗p ≤ 0.001, ∗∗∗∗p ≤ 0.0001. For further details on genetic crosses and statistical tests, see STAR Methods.

Blocking spinal inhibition generates exaggerated responses to tactile stimuli, resembling mechanical allodynia^97^. This suggests that inhibition exerts a control over tactile signals to prevent them from being perceived as something more noxious. Although inhibition in the DCN has been associated with sensory feedback during dextrous forelimb reaching behavior^43^, the role of DCN inhibition in setting normal tactile sensitivity is not well understood. Our electrophysiological data highlighted robust inhibitory control of VPL-PN activity **(Figure 3G-N).** Therefore, we selectively manipulated Gr inhibition to better understand how exaggerated tactile signals in the DCN influence tactile perception. To selectively manipulate Gr inhibition, we injected a Cre-dependent hM4DI-DREADD virus into the Gr of *Slc32a1^Cre^* mice (aka *Vgat^IRES-Cre^* mice) **(Figure 4E)**, resulting in hM4Di expression in Gr inhibitory neurons **(Figure 4F)**. Reducing Gr inhibition produced tactile hypersensitivity to lower force von Frey filaments, similar to mice following spared nerve injury **(Figure 4G)**, as well as increased intensity of response to a dynamic brush stimulus **(Figure 4H)**. Similar to manipulations of VPL-PNs, CNO did not affect sensitivity to a noxious mechanical pinprick stimulus **(Figure S4E)** or radiant thermal sensitivity using the Hargreaves assay **(Figure S4F)**. Given that CNO-induced tactile hypersensitivity resembled neuropathic pain, we next whether reducing Gr inhibition was intrinsically aversive in the absence of applied mechanical stimulation. To explore this, we performed condition place preference/aversion tests in *Vgat^IRES-Cre^*, associating one chamber side with CNO treatment (silencing Gr inhibition). Mice spent less time on the CNO-conditioned side post treatment, suggesting aversion to the chemogenetic silencing of Gr inhibition **(Figure 4I)**. As a control for off-target effects of CNO, we injected a Cre-dependent GFP virus (pAAV-hSyn-DIO-EGFP) into *Vgat^IRES-Cre^*mice **(Figure S4G-H)**. We found no significant differences between CNO and saline treatment for von Frey **(Figure S4I)**, dynamic brush **(Figure S4J)**, pinprick **(Figure S4K)**, Hargreaves **(Figure S4L)**, or place preference **(Figure S4M)**, suggesting that CNO alone does not have any effect on tactile or thermal sensitivity, and is not generally preferred or aversive. These results suggest that disruption of Gr inhibition generates a tactile hypersensitivity to mechanical low-threshold stimuli, but does not affect noxious mechanical or thermal responses.

Silencing Gr inhibition heightened tactile sensitivity to mechanical low-threshold stimuli. However, it is unclear whether these observed behaviors represent sensory-evoked responses of animals in a pain condition. A loss of spinal inhibition produces exaggerated paw withdrawal responses, such as shaking and guarding, which mirror mechanical allodynia following neuropathic pain^46,97–100^. In order to investigate how neuropathic pain or silencing Gr inhibition alters the paw withdrawal response, we utilized high speed videography paired with machine learning to record and quantify paw withdrawal to different mechanical stimuli. We used DeepLabCut^101^ for markerless pose estimation of the metatarsophalangeal joint (MTP) **(Figure 4J)** and utilized Paw Assessment at Withdrawal Speeds (PAWS^102^) to extract relevant features of the response, including shaking and guarding, which are indicative of pain^102^. First, we compared withdrawal responses to a 0.6 VF filament, a dynamic brush, or a pinprick in uninjured mice and mice with an SNI model of neuropathic pain^103^. Following SNI, mice developed tactile hypersensitivity which closely resembled the tactile hypersensitivity induced by silencing Gr inhibitory neurons **(Figure 4G)**. Additionally, SNI mice exhibited altered paw withdrawal responses when stimulated with either innocuous (0.6 VF, brush) or noxious (pinprick) stimuli **(Figure 4K, Supplemental Table 1)**, representing features of both mechanical allodynia and hyperalgesia. Comparatively, reducing Gr inhibition enhanced responses to 0.6 VF and brush, but not to pinprick **(Figure 4K, Supplemental Table 2)**, resembling mechanical allodynia, but not hyperalgesia. These results suggest that silencing Gr inhibitory neurons specifically enhances responses to tactile stimuli, which is also observed during neuropathic pain, but does not affect responses to noxious mechanical stimuli.

Next, we took an unbiased approach to identify key paw withdrawal signatures that were induced by either SNI or chemogenetic silencing of Gr inhibitory neurons. We used an unsupervised machine learning approach known as B-SOiD^104^, which has been previously utilized to identify pain-related paw withdrawal signatures of inflammatory pain^105^. As inputs, we utilized our tracked MTP data as well as two reference points (initial MTP coordinate, MTP coordinate at maximum height of withdrawal) from mice where Gr inhibitory neurons were silenced as well as mice with SNI **(Figure 4J,K)**. B-SOiD then extracted relevant behavioral clusters of movements utilized across the dataset **(Figure 4L)**. Out of 9 identified clusters, about half represented stationary periods within the withdrawal while the others exhibited low usage across all groups. However, one cluster consistently identified a rapid shaking behavior consisting of multiple shakes. We quantified the usage of this “shaking” syllable across all groups in response to different stimuli, and found that in response to a brush stimulus, both SNI mice and mice where Gr inhibitory neurons were silenced showed an increased usage of this shaking syllable **(Figure 4L)**. Comparatively, in response to pinprick only SNI mice showed an increase in the usage of the shaking module, while silencing Gr inhibition had no effect **(Figure 4L)**. These results suggest that manipulation of Gr inhibition specifically exaggerates responses to innocuous mechanical stimuli and not to noxious mechanical stimuli.

In summary, chemogenetic silencing of Gr VPL-PNs reduced mechanical sensitivity to low-threshold, innocuous stimuli while silencing Gr inhibitory neurons increased mechanical sensitivity to low-threshold stimuli in a manner resembling tactile hypersensitivity during neuropathic pain. Additionally, Gr manipulations did not affect noxious mechanical or thermal sensitivity. These results suggest that Gr VPL-PNs are important for the transmission of tactile information to the brain, while Gr inhibitory neurons set normal levels of tactile sensitivity. Additionally, silencing Gr inhibition results in tactile hypersensitivity resembling SNI-induced mechanical allodynia, but does not affect noxious mechanical or thermal sensitivity underlying SNI-induced hyperalgesia.

### Increased tactile-evoked activity and altered primary afferent drive onto Gr VPL-PNs during neuropathic pain

Silencing Gr inhibition resulted in tactile hypersensitivity resembling neuropathic pain **(Figure 4G,H,K,L)**, suggesting that enhanced Gr output may contribute to tactile-evoked manifestations of neuropathic pain. Additionally, reduction of DCN activity via pharmacological intervention or lesion attenuates mechanical allodynia during neuropathic pain^21,106^. Therefore, we next investigated whether Gr activity is increased during neuropathic pain. We focused on activity evoked by low-threshold mechanical stimuli, which normally does not elicit a painful response^2,67^. Previous studies suggest that electrical stimulation of injured peripheral nerves results in an increased recruitment of Gr neurons compared to stimulation of uninjured nerves^23,107^, with more VPL-PNs especially being recruited^24^. Although this suggests increased sensory-evoked activity in the DCN, these stimulations do not physiologically mimic tactile stimuli. To investigate tactile-evoked Gr activity, we applied a brush stimulation to the plantar hindpaw of anesthetized mice, and performed immunohistochemistry for c-Fos, a marker for neuronal activity^24^. We observed an increased number of Fos+ cells **(Figure 5A-B)** in the Gr of SNI mice compared to sham mice, suggesting increased recruitment of Gr neurons in response to tactile stimuli post-injury. Additionally, we observed robust microglial recruitment in the Gr post-injury **(Figure S5A-B)**, which mirrored spinal glial recruitment associated with the maintenance of neuropathic pain^108,109^.

**Figure 5.**
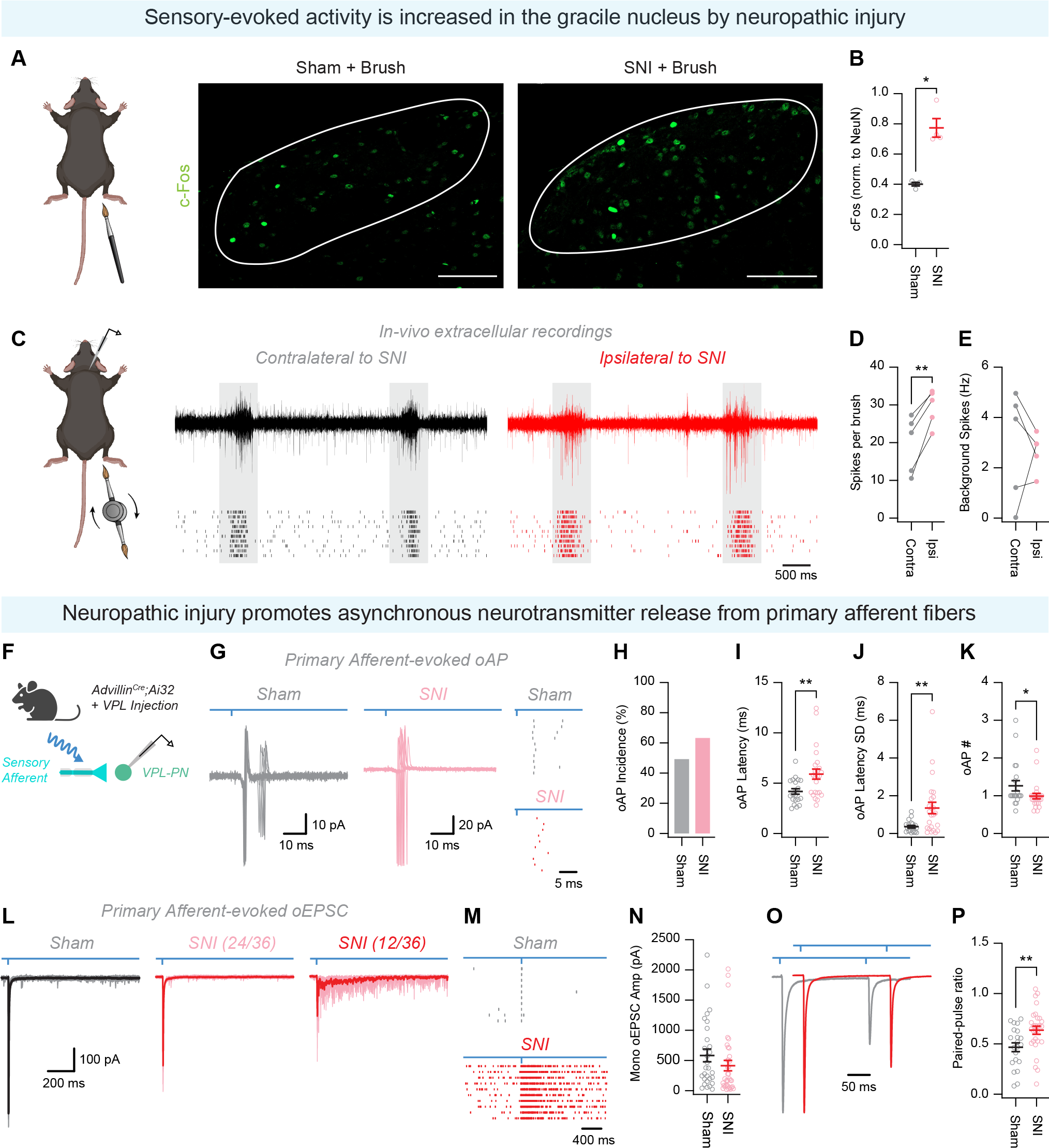
Increased tactile-evoked activity and altered primary afferent drive onto Gr VPL-PNs during neuropathic pain A) c-Fos immunolabeling (green) was used to identify Gr neurons active to a brush stimulus in sham and SNI mice. Scale bar = 100 um. B) Comparison in number of cFos+ cells (normalized to total NeuN) in response to brush stimulation in sham and SNI mice. C) *In vivo* recordings were conducted in anesthetized mice with unilateral SNI. Recording electrodes were placed in the Gr on the side ipsilateral to the injury (Ipsi) and the Gr contralateral to the injury (Contra). Motorized brushes were positioned above the Gr and rotated to apply two brush stimuli per complete rotation. Evoked activity is highlighted by gray boxes, while space between the boxes represents spontaneous activity. D,E) Evoked (D) and spontaneous (E) spiking activity in the ipsilateral and contralateral Gr. G) Cell-attached voltage clamp recording showing optically-evoked AP (oAP) following primary afferent photostimulation in sham (grey) and SNI (red). 10 consecutive sweeps overlaid. Right: raster plot of example recordings. H) Percentage of neurons that exhibit oAPs following primary afferent photostimulation. I) Quantification of oAP latency. J) Quantification of oAP latency standard deviation. K) Quantification of the number of oAPs evoked per stimulus. L) Whole-cell voltage clamp recordings of optically-evoked EPSC (oEPSC) from VPL-PNs in sham (grey) and SNI (red) mice following primary afferent photostimulation. 10 consecutive sweeps with average overlaid. Note, following SNI ⅓ VPL-PNs exhibited prolonged activity following primary afferent photostimulation (right). M) Raster plot of EPSC incidence in sham (top) and SNI (bottom) VPL-PNs. N) Quantification of oEPSC amplitude. O) Whole-cell voltage clamp recording (-70 mV) from VPL-PNs in sham (grey) and SNI (red) mice showing oEPSC paired-pulse response to primary afferent photostimulation. P) Quantification of oEPSC paired-pulse ratio. Data are reported as mean values ± SEM. *p ≤ 0.05, ∗∗p ≤ 0.01 , ∗∗∗p ≤ 0.001, ∗∗∗∗p ≤ 0.0001. For further details on genetic crosses and statistical tests, see STAR Methods.

Increased Gr recruitment suggests an overall increase in tactile-evoked activity in the Gr during neuropathic pain. To investigate this, we performed *in vivo* extracellular recordings in the Gr of anesthetized mice with SNI **(Figure 5C)**, recording the activity of ipsilateral and contralateral Gr in response to dynamic brush of the corresponding hindpaw. Stimulation of the injured hindpaw resulted in significantly higher evoked spiking activity compared to contralateral hindpaw stimulation **(Figure 5D)**. Additionally, there was no difference in spontaneous spiking activity between ipsilateral and contralateral hindpaw **(Figure 5E)**. Collectively, these data suggest that Gr neurons exhibit heightened responses to tactile stimuli during neuropathic pain. Whether this increased Gr excitability is attributed to increased strength of tactile inputs, increased excitability of VPL-PNs, or changes to Gr inhibition is not well characterized.

To better understand Gr hyperactivity during neuropathic pain, we initially characterized the intrinsic properties of Gr VPL-PNs and inhibitory neurons. Surprisingly, we found no difference in responsiveness to depolarizing step currents in either population **(Figure S6A-C, H-P)**. We also examined the properties of the threshold AP in each population **(Figure S6D-M, Q-Z)**, and found that VPL-PNs APs exhibit faster kinetics, and inhibitory neurons APs are unchanged following SNI. While changes to AP kinetics may influence downstream neurotransmitter release^110^, these data suggest unaltered intrinsic excitability of Gr neurons post-injury. Next, we examined the spontaneous activity and excitatory drive to Gr VPL-PNs or inhibitory neurons. In-line with previous work^22^, following SNI, VPL-PNs exhibited increased spontaneous AP discharge frequency **(Figure S7A-C)**, and increased frequency of spontaneous excitatory postsynaptic currents (sEPCS), without changes to amplitude or kinetics **(Figure S7D-I)**. Conversely, inhibitory neurons displayed no changes in spontaneous AP discharge **(Figure S7J-L)**, a reduction in sEPSC amplitude, and no changes in sEPSC frequency or kinetics **(Figure S7G-L)**. These findings align with our earlier observations that the activity of VPL-PNs and inhibitory neurons is governed by distinct presynaptic circuitry **(Figure 3B-I)**. Increased excitatory drive to VPL-PNs, with reduced input to Gr inhibitory neurons has the potential to shift the excitation/inhibition balance of the Gr toward a hyperexcitable state.

To explore the potential contribution of altered input from primary afferents or spinal projections to Gr hyperactivity, we conducted slice electrophysiology in sham or SNI mice, where primary afferent or spinal terminals in the Gr could be optogenetically activated while recording from Gr VPL-PNs or inhibitory neurons. Following SNI, optogenetic activation of primary afferent terminals elicited AP discharge from more VPL-PNs than in sham mice **(Figure 5F-H)**. Intriguingly, we observed an increase in the latency **(Figure 5I)**, and latency standard deviation **(Figure 5J)** of optically-evoked APs post-SNI, alongside a reduction in the number of APs per stimulus **(Figure 5K)**, indicating shift towards decreased reliability of transmission following nerve injury.

In-line with this increased variability, we observed a shift in primary afferent input to VPL-PNs from synchronous to asynchronous release following SNI **(Figure 5L-M)**, accompanied by an increase in paired-pulse ratio (reduced release probability) **(Figure 5O-P)**. This finding is aligned with synchronicity of release being reduced at low release probability synapses^111^. In contrast to the changes observed in primary afferent to VPL-PN synapses, we found no changes in primary afferent to inhibitory neuron **(Figure S8A-F)**, spinal projection to VPL-PNs **(Figure S8G-L)**, or spinal projection to inhibitory neuron **(Figure S8M-R)** synapses following SNI. Interestingly, we observed a mild reduction in primary afferent evoked inhibition onto VPL-PNs post-SNI **(Figure S9B)**, which could unveil polysynaptic excitatory circuitry^95^. Nevertheless, inhibitory receptor antagonists failed to induce prolonged excitatory responses in VPL-PNs **(Figure S9C)**, suggesting that reduced inhibitory input might not account for the observed changes. Additionally, no alterations were observed in sIPSCs recorded from VPL-PNs post-SNI **(Figure S9D-K)**, and optogentically-evoked IPSCs exhibited increased amplitude post-injury, with no changes in GABA/glycine composition **(Figure S9L-O)**.

In summary, we observed an increase in c-Fos+ neurons in the Gr as well as increased Gr spiking activity in response to tactile stimulation during neuropathic pain. Gr VPL-PNs were more likely to fire action potentials in response to primary afferent activation, albeit with increased variability and reduced AP count per stimulus.

This altered Gr signaling was primarily attributed to a shift in primary afferent signaling from synchronous to long duration asynchronous VPL-PN excitation. Together, these results suggest that the observed hyperexcitability within the Gr following SNI stems from an increase in the number of responsive neurons rather than an amplified response of each neuron to tactile stimulation.

### Enhancing Gr inhibition alleviates tactile hypersensitivity and neuropathic-pain induced paw withdrawal signatures

Blockade of DCN activity via pharmacology or lesion ameliorates tactile hypersensitivity during neuropathic pain^21,106^. This work suggests that DCN activity contributes to tactile hypersensitivity during neuropathic pain. However, the specific functional contributions of distinct Gr populations, namely VPL-PNs and inhibitory neurons, to tactile hypersensitivity is not well understood. Since manipulation of Gr VPL-PNs or inhibitory neurons bidirectionally scales tactile responses in naive mice **(Figure 4C,G)**, and can even result in neuropathic pain-like paw withdrawal features **(Figure 4J-L)**, we hypothesized that inhibition of Gr VPL-PNs or activation of Gr inhibitory neurons during neuropathic pain would alleviate tactile allodynia.

Gr VPL-PNs display an increased recruitment to electrical stimulation of a transected peripheral nerve^24^, yet their direct contribution to tactile hypersensitivity is not well characterized. To specifically manipulate VPL-PNs, we injected a retrograde Cre virus into the VPL and injected a Cre-dependent hM4Di inhibitory DREADD into the Gr of SNI mice **(Figure 6A-B)**. SNI mice developed the expected punctate tactile hypersensitivity compared to naive mice, a response that was partially alleviated by administration of CNO **(Figure 6C)**. CNO administration did not affect dynamic mechanical sensitivity to a brush stimulation **(Figure 6C)**. Additionally, CNO did not affect noxious mechanical sensitivity in response to a pinprick **(Figure S10B)**, thermal sensitivity, **(Figure S10C)**, or induce place preference **(Figure S10D)**. Therefore, chemogenetic inhibition of VPL-PNs selectively and partially reduced tactile hypersensitivity during neuropathic pain, but did not affect mechanical or thermal hyperalgesia.

**Figure 6.**
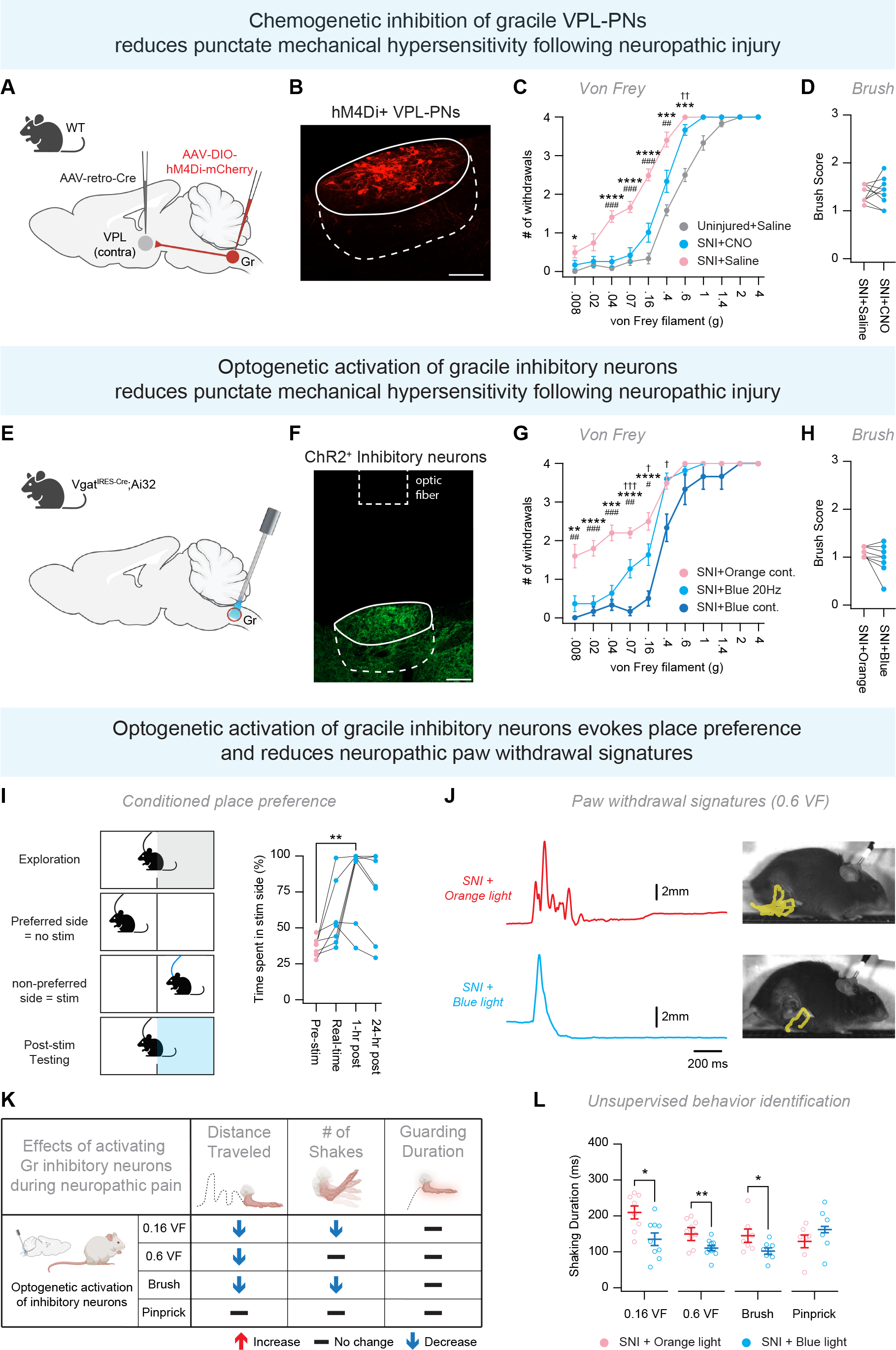
Enhancing Gr inhibition alleviates tactile hypersensitivity and/or neuropathic-pain induced paw withdrawal signatures A,B) To silence Gr VPL-PNs during neuropathic pain, we injected a retrograde-Cre virus into the VPL, and injected a Cre-dependent inhibitory hM4Di DREADD virus (red) into the Gr. Mice then received an SNI procedure to induce neuropathic pain. VPL-PNs were silenced by administering CNO, while saline was used as a control. Scale bar = 100 um. C) Punctate mechanical sensitivity in response to von Frey stimulation in SNI mice given saline (red) or CNO (blue), or uninjured mice given saline (gray). *=Uninjured+Saline vs SNI+Saline, ^#^=SNI+Saline vs SNI+CNO, ^†^=Uninjuired+Saline vs SNI+CNO. D) Dynamic mechanical sensitivity in response to a brush stimulation in SNI mice given saline (red) or CNO (blue). E,F) Strategy to activate inhibition in the Gr. We generated *Vgat^IRES-Cre^*;Ai32 mice, where ChR2 was expressed in inhibitory neurons (green). An optic probe was positioned above the Gr to allow for photostimulation of the Gr. Mice were then given the SNI procedure to induce neuropathic pain. Stimulation with blue light was used for activation of Gr inhibition, while orange light was used as a control. G) Punctate mechanical sensitivity in response to von Frey stimulation in SNI mice given stimulation of continuous orange light (red), 20 Hz blue light (light blue), and continuous blue light (dark blue). *=Orange cont vs Blue cont, ^#^=Orange cont vs Blue 20Hz, ^†^=Blue cont vs Blue 20Hz. H) Dynamic mechanical sensitivity in response to a brush stimulation in SNI mice given stimulation of continuous orange light (red) or continuous blue light (blue). I) Conditioned place preference assay to assess appetitive/aversive nature of activating Gr inhibition in SNI mice. Time spent on the non-preferred side was measured during real-time stimulation, 1 hour post-conditioning, and 24 hours post-conditioning. J) DeepLabCut was used to track the coordinates of the hindpaw in response to a mechanical stimulus. Representative traces of metatarsophalangeal joint (MTP) y-coordinates during withdrawal responses to a 0.6g von Frey filament during stimulation with orange light (red) or blue light (blue). K) Pain Assessment at Withdrawal Speeds (PAWS) software was used to extract features of the paw withdrawal response including distance traveled, number of shakes, and guarding duration. Further explanation of how this software extracts relevant features can be found in Jones et al., 2020^102^. Features for mice with orange light stimulation were compared to features of mice with blue light stimulation. ↑ represents an increase compared to control mice, ↓ represents a decrease compared to control mice, and - represents no change. L) First three dimensions of UMAP (uniform manifold approximation and projection) embedding, showing in low-dimensional space a projection of extracted behavioral features. Colors represent behavioral clusters identified by HDBSCAN (hierarchical density-based spatial clustering of applications with noise), including a cluster corresponding to shaking behavior (turquoise). Graph shows comparison of normalized duration of shaking behavior in mice with orange light stimulation versus blue light stimulation. Data are reported as mean values ± SEM. *p ≤ 0.05, ∗∗p ≤ 0.01 , ∗∗∗p ≤ 0.001, ∗∗∗∗p ≤ 0.0001. For further details on genetic crosses and statistical tests, see STAR Methods.

Enhancing inhibition in the spinal cord is able to alleviate symptoms of neuropathic pain^65,112–116^. Inhibition in the DCN is suggested to become impaired during neuropathic pain^29^, which may contribute to neuropathic pain-like behaviors observed after silencing inhibition in the Gr **(Figure 4E-L)**. Therefore, we hypothesized that enhancing Gr inhibition could ameliorate tactile allodynia during neuropathic pain. We generated *Vgat^IRES-Cre^*,Ai32 mice, where ChR2 was expressed in inhibitory neurons and an optic probe was placed above the Gr to allow for optical stimulation of local inhibitory terminals **(Figure 6E-F)**. To determine if activating Gr inhibition could reduce tactile hypersensitivity during neuropathic pain, we first assessed von Frey sensitivity under different optical paradigms. Given that multiple activation paradigms can be used to increase inhibition^117–120^, we provided either a 20Hz stimulation or continuous stimulation using 470 nm blue light to activate ChR2^121–124^. We found that 20Hz stimulation was less effective at reducing tactile hypersensitivity than constant blue light stimulation **(Figure 6G)**, which specifically reduced tactile stimuli for lower threshold von Frey stimuli (0.008-0.4g). Thus, we utilized continuous blue light stimulation to further assess the functional contributions of DCN inhibition. As a control for channel-independent heating during optical stimulation, we utilized a wavelength of light outside of the excitation range for ChR2^125^. Blue light stimulation did not reduce dynamic mechanical allodynia in response to brush **(Figure 6H)**. Additionally, neither blue nor orange light stimulation affected mechanical hyperalgesia induced by a pinprick **(Figure S10F)**, or thermal hyperalgesia in the Hargreaves assay **(Figure S10G)**. Collectively, activation of Gr inhibition specifically reduced punctate tactile hypersensitivity to lower threshold von Frey stimuli.

Given the effects of Gr manipulations on tactile allodynia, we investigated whether activating Gr inhibition was sufficient to induce place preference, suggestive of an analgesic effect^126^. We used a two-chamber place preference assay where mice were allowed to develop an initial preference, and then optogenetic stimulation was applied on the non-preferred side. We analyzed both real-time place preference for the first stimulation trial, as well as conditioned place preference for time points following the last conditioning trial. Interestingly, mice did not display a real-time place preference, but did display preference for the stimulated side 1 hour following the final conditioning trial **(Figure 6I)**. These results suggest that activation of Gr inhibition is preferred during neuropathic pain and may provide pain relief.

We next investigated whether activating Gr inhibition could alleviate affective features of the paw withdrawal associated with tactile allodynia. We once again used PAWS to track the paw withdrawal response to innocuous or noxious mechanical stimuli to identify shaking and guarding behavior **(Figure 6J)**. Optogenetic stimulation of Gr inhibition resulted in decreased distance traveled for 0.16g VF, 0.6g VF, and brush as well as decreased shaking for 0.6g VF, but did not affect pinprick **(Figure 6K)**. Additionally, optogenetic stimulation reduced the maximum velocity in response to 0.16g and 0.6g VF, but not in response to pinprick **(Supplemental Table 3)**. These results suggest that activation of Gr inhibition specifically reduces the intensity of paw withdrawal responses to tactile stimuli, but not to noxious stimuli. Next, we utilized B-SOiD as an unsupervised approach to investigate whether paw withdrawal signatures of neuropathic pain were reduced by activation of Gr inhibition. We focused on the previously defined “shaking” module **(Figure 4L)**, which had a reduced usage in the blue light stim group compared to the no light group **(Figure 6L)**. Collectively, activation of Gr inhibition resulted in decreased tactile sensitivity specific to lower-threshold von Frey, and reduced affective shaking behavior to innocuous but not noxious mechanical stimuli. In conclusion, activating Gr inhibition specifically alleviated tactile allodynia, but did not affect mechanical or thermal hyperalgesia.

In summary, reducing Gr VPL-PN activity or activating Gr inhibition during neuropathic pain reduces tactile hypersensitivity for lower threshold von Frey stimuli, pointing to specific amelioration of tactile allodynia. Gr manipulations did not affect noxious mechanical or thermal sensitivity, suggesting no effect on hyperalgesia. Additionally, activation of Gr inhibition reduced the severity of paw withdrawal responses to innocuous stimuli and reduced affective shaking behavior associated with pain. Therefore, inhibition in the Gr can be leveraged to reduce tactile hypersensitivity and tactile allodynia during neuropathic pain.

## DISCUSSION

In this study, we found that the gracile nucleus (Gr) of the DCN consists of VPL-projecting neurons (VPL-PNs) as well as local inhibitory neurons which are differentially innervated by ascending primary afferents and spinal projections. Functional manipulations of either Gr population scaled tactile sensitivity: inhibition of Gr VPL-PNs reduced mechanical sensitivity to low-threshold von Frey stimuli, while inhibition of Gr inhibitory neurons increased tactile sensitivity and generated paw withdrawal signatures that resembled responses of mice with tactile allodynia during neuropathic pain. Investigation of the Gr during neuropathic pain revealed increased tactile-evoked activity, coupled with altered excitatory drive from primary afferents onto VPL-PNs. Optogenetic activation of Gr inhibition alleviated tactile allodynia and reduced paw withdrawal features associated with neuropathic pain. These results suggest that manipulation of either Gr VPL-PNs or Gr inhibitory circuits can scale tactile responses in both naive and neuropathic pain conditions. Therefore, the DCN represents a relevant region of study not only for understanding tactile sensitivity, but also for tactile-specific manifestations of neuropathic pain such as tactile allodynia.

### Gr circuits promote VPL-PN signal transduction modulated by sensory-driven feedforward inhibition

Classical work defines the DCN as a tactile region receiving DRG and spinal inputs and projecting to the VPL. Expanding upon this, we used an extensive mouse molecular toolkit to map various DRG inputs into the Gr and found dense contributions from Aβ RA-LTMRs and Aβ Field-LTMRs and relatively minor contributions from Aβ SA-LTMRs and proprioceptors, supporting previous anatomical characterizations^68–70,127^. Overall, primary afferent inputs in the Gr appeared more dense than those from spinal projections, matching other reports^42^. Consistent with this, slice electrophysiology revealed stronger inputs from primary afferents onto VPL-PNs compared to spinal projections. By comparison, Gr inhibitory neurons displayed a similar predominance of primary afferent input over spinal, suggesting tactile responses in the Gr are largely driven by the direct DRG pathway. However, VPL-PNs are more excitable and receive stronger inputs from both primary afferents and spinal projections compared to inhibitory neurons, suggesting that sensory-driven activity in the Gr would predominantly activate VPL-PNs followed by inhibition. This is consistent with work suggesting that Gr VPL-PNs have stronger intensity encoding than Gr inhibitory neurons^37^. Consistent with previous reports^37^, VPL-PNs were more spontaneously active than inhibitory neurons. Therefore, VPL-PNs transmit both spontaneous and sensory-evoked activity to the VPL, while inhibitory neurons are likely to be more dependent on sensory-driven activation. Given that inhibitory neurons largely target the VPL core over the shell, this model supports the idea of sensory-drive feedforward^39^ and surround inhibition of VPL-PNs^37^ in the Gr.

Additionally, feedback mechanisms in the DCN acting via descending cortical projections^39,128^ may further modulate Gr VPL-PN activity in a sensory-driven manner. Within the cuneate nucleus corticofugal neurons target inhibitory neurons to inhibit somatosensory feedback^43^. A similar organization in the Gr may allow top-down pathways to engage Gr inhibition to modulate the coding of bottom-up sensory signals.

### The Gr selectively scales tactile sensitivity

Manipulations of either Gr VPL-PNs or Gr inhibitory neurons altered tactile sensitivity as well as signatures of paw withdrawal responses to tactile stimuli, but did not affect noxious mechanical or thermal sensitivity or paw withdrawal to noxious mechanical stimuli **(Figure 4C,D,G,H, Figure S4B,C,E,H)**. These results indicate that the Gr specifically mediates responses to low-threshold, tactile stimuli. These findings are consistent with evidence that spinal cord circuits mediate the nociceptive reflex response to a noxious pinprick or radiant heat stimuli^129–131^, while spinal cord lamina I projection neurons transmit nociceptive information to supraspinal centers for pain perception^56,132^. Therefore, noxious mechanical and thermal stimuli have anatomical pathways distinct from the DCN, which is why their nociceptive signaling is unaffected by Gr manipulations. During neuropathic pain, silencing VPL-PNs or activating Gr inhibition reduced tactile hypersensitivity, but did not affect noxious mechanical or thermal hypersensitivity **(Figure 6C,D,G,H, Figure S10B,C,D,F,G)**. Additionally, activation of Gr inhibition reduced pain-associated paw withdrawal signatures in response to tactile stimuli, but did not affect responses to noxious stimuli **(Figure 6J.K, Supplemental Table 3)**. These results are consistent with studies leveraging inhibition in the spinal cord to treat chronic pain^65,114–116^, though effects in the Gr were specific to tactile hypersensitivity. Based on these results, it is unlikely that enhanced signaling in the Gr during neuropathic pain contributes to noxious mechanical or thermal sensitivity or withdrawal signatures in response to nociceptive stimuli. Similar to the naive condition, the Gr specifically mediates tactile sensitivity during neuropathic pain, while mechanical and thermal hyperalgesia were unaffected. Consistent with this, administration of lidocaine, a local anesthetic, into the Gr alleviated mechanical hypersensitivity but did not affect thermal hypersensitivity^106^. Additionally, lesion of the dorsal columns can attenuate mechanical allodynia following nerve injury^21^ or spinal hemisection^133^, but does not affect thermal responses. Collectively, these results suggest the Gr is indeed a tactile, mechanical pathway as classically described, but also plays a role in tactile-specific manifestations of neuropathic pain. This is of particular interest for the management of chronic pain, which is maladaptive, while leaving acute, protective pain unaffected. Future work identifying therapeutically tractable targets within the Gr, or examining how different dorsal column/spinal cord stimulation paradigms^134^ may influence Gr neurons activity will be of great interest for the treatment of tactile-specific chronic pain conditions such as tactile allodynia.

### Increased recruitment and altered tactile coding in Gr following nerve injury

Withdrawal thresholds, as well as the intensity of withdrawal responses could be scaled up or down by reducing or enhancing Gr inhibition, respectively. In naive conditions, reducing Gr inhibition closely resembled behavioral phenotypes of nerve injury-induced neuropathic pain, including tactile hypersensitivity and enhanced paw withdrawal signatures. Importantly, reducing Gr inhibition only altered sensitivity and withdrawal signatures to tactile stimuli (allodynia), while nerve injury heightened responses to both innocuous and noxious mechanical stimuli (allodynia and hyperalgesia). Therefore, these findings additionally suggest that Gr signaling is important for the detection and intensity-encoding of tactile stimuli, but is not important for the sensitivity or intensity of responses to noxious mechanical or thermal stimuli. Following nerve injury, enhanced Gr activity is associated with neuropathic pain^22,24^. Consistent with this, post-neuropathic injury c-Fos labeling showed increased neural recruitment, *in-vivo* extracellular recordings showed enhanced tactile-evoked responses, and slice electrophysiology recordings showed increased primary afferent recruitment of VPL-PNs which we attributed to a shift in primary afferent signaling to prolonged asynchronous glutamate release. Along with an increase in the number of responsive neurons, we also observed an increase in the latency to and variability of VPL-PN activation following primary afferent stimulation. Similar asynchronicity in neural activity, increased latency to first spike, and increased jitter in response to tactile stimuli following nerve injury has been reported in the spinal cord dorsal horn ^135^, and increased response latencies to Aβ-LTMR stimulation, as well as afterdischarges in response to mechanical stimulation have been reported in Gr VPL-PNs following peripheral nerve injury^22^. Together, this work suggests neuropathic injury alters the temporal dynamics of tactile signaling and associated activity following neuropathic injury. In-line with previous reports^22^, we also observed an increase in spontaneous discharge in VPL-PNs. This increased background activity, coupled with altered signal reliability and reduced synchronicity could introduce signal noise and lower the fidelity of tactile signaling. Given the known role of the dorsal column nuclei in tactile discrimination^16^, such changes could contribute to altered tactile acuity in chronic pain states^136^. Future work should determine whether these changes in Gr signaling are conserved in the awake, behaving mouse, and how neural responses within the Gr are correlated with specific pain behaviors such as paw withdrawal or shaking.

Chemogenetic silencing of Gr inhibitory neurons resulted in increased tactile sensitivity and exaggerated tactile-evoked paw withdrawal responses. These results are similar to those observed in the spinal cord, where impairing spinal inhibition results in allodynia^97,100,137^. In the spinal cord, reduced inhibition manifests as a reduction in presynaptic inhibition^44^, as well as reduced postsynaptic inhibition of excitatory dorsal horn interneurons and projection neurons^95,138^. As a consequence of reduced inhibition within the spinal cord dorsal horn, innocuous tactile information normally restricted to the deeper lamina engage lamina I nociceptive circuits and facilitate allodynia^6,7,139^. However, this model of allodynia in the spinal cord cannot be applied to the DCN due to the inherent tactile nature of DCN circuits. While we did observe a moderate reduction in the incidence of sensory-evoked inhibition, we found no changes in spontaneous inhibition or inhibitory transmission by inhibitory neurons onto Gr VPL-PNs. Therefore, rather than an “ungating” of tactile information, it is likely that altered DCN activity during neuropathic pain acts on target structures that process touch and pain. A likely candidate is the VPL, which contains neurons that preferentially respond to either innocuous or noxious mechanical information^140^. Recently, the DCN→VPL pathway has been suggested to transmit high-force information coming from PSDCs which are activated by high-threshold mechanoreceptor (HTMR) stimulation^37^. Additionally, enhanced sensory activity in the VPL during neuropathic pain can be attenuated by lesioning the dorsal columns^141^. Therefore, enhanced tactile-evoked activity in the DCN during neuropathic pain may result in increased signaling to the VPL, which may be perceived as high-force or noxious.

Additionally, DCN projections to the periaqueductal gray (PAG)^142^ and spinal cord^143^ have been described, suggesting the DCN potentially has alternate routes for controlling tactile and nociceptive information.

### Redundant pathways for the modulation of neuropathic pain

Classical views on sensory processing suggest divergent pathways for pain and touch: the anterolateral tract transmits noxious mechanical and thermal information^144^ while the dorsal column medial lemniscal tract transmits innocuous touch^145^. Although lesions of the dorsal columns are generally associated with impairments in fine tactile perception^89^, general tactile perception is still retained following lesions where only the anterolateral tract is retained^146^. Further, the Calca neurons within the parabrachial nucleus have been shown to be necessary and sufficient for the establishment and maintenance of mechanical allodynia^94^, while manipulation of excitatory neurons within the parabrachial nucleus can induce or alleviate allodynic behavior^8^. This suggests that tactile information can travel through multiple routes to get to the brain. Therefore, rather than considering the anterolateral and dorsal column medial lemniscal pathways as distinct, one can consider a more comprehensive model where the combined activity of individual pathways contribute to overall sensory state. This idea is supported by studies showing that the parabrachial nucleus receives both tactile and nociceptive information and is a key regulator of neuropathic pain^8,9,147^. Therefore, there are likely to be multiple supraspinal pathways involved in the manifestation of neuropathic pain. Further upstream, both spinothalamic and DCN projections target the same area of the VPL^148^, which receives convergent innocuous and noxious information. Importantly, the lateral thalamus has been strongly implicated in chronic pain processing^149^, and exhibits plasticity following nerve injury^150^. Therefore, neuroplasticity in the VPL during pathological conditions^79,150,151^ may alter how incoming tactile information is processed, which may aberrantly contribute to tactile-evoked pain perception. Further studies on how DCN input to the VPL is interpreted during naive and chronic pain conditions are required.

## Limitations of the study

While our study utilized a range of techniques including in-vivo extracellular recordings, slice patch-clamp recordings, and optogenetic/chemogenetic manipulations to elucidate the circuits and functional role of the Gr in scaling tactile sensitivity in both naive and neuropathically injured mice, several limitations should be acknowledged. Firstly, *in-vivo* extracellular recordings and c-Fos activity measurements were conducted in anesthetized mice, and it’s well-established that anesthetics can significantly influence pain circuits.

Additionally, patch-clamp recordings were performed in a reduced acute slice preparation, which may not fully capture the complexities of neural activity in intact systems. Future studies employing recordings in awake, behaving animals would provide a more physiologically relevant assessment of Gr neural activity in response to sensory stimuli. Moreover, the optogenetic approaches utilized in slice electrophysiology and behavior have inherent limitations. It’s improbable that in a physiological setting, large populations of neurons are synchronously activated to the extent achieved by ChR2-mediated activation. Lastly, our study focused on a specific injury model (SNI), which affects sensory coding of stimuli originating from the injured hindlimb.

However, it’s important to note that the Gr receives sensory input from a broader region spanning T6-S5. Therefore, the changes observed following neuropathic injury may not fully represent hindlimb-specific circuit alterations but offers insights into changes in excitatory input to the Gr from the lower body. Future studies examining how somatopically distinct regions are impacted by injury will provide further insight into region specific changes and changes to receptive fields.

## ACKNOWLEDGEMENTS

We are grateful to Dr. Sliman Bensmaia for providing macaque brainstem tissue. We would like to thank Hanns Ulrich Zeilhofer and Hendrik Wildner for GlyT2-GFP mouse tissue. We would like to thank Sean O’Leary and Yurdiana Hernandez for their assistance with animal care and genotyping. Financial support was provided by the Pew Charitable Trust (V.E.A.), Rita Allen Foundation (V.E.A), NJ Commission on Spinal Cord Research (M.A.G., V.E.A.); NIH NINDS K01NS116224 (V.E.A.), NIH NINDS R01NS119268 (V.E.A.), NIH NINDS R01NS124799.

## AUTHOR CONTRIBUTIONS

A Upadhyay, M.A. Gradwell, and V.E. Abraira conceptualized the study and designed experiments. A. Upadhyay performed and analyzed histology, viral tracing, and behavioral experiments. M.A. Gradwell performed and analyzed electrophysiology experiments. J. Conner performed and analyzed *in vivo* recordings. K. Patel and C. Azadegan aided with histology and behavioral experiments. F. Imai performed cortical viral injections. A. Sanyal, J.K. Thackray, T.J. Vajtay, and S. Ogundare aided with Deeplabcut tracking and PAWS analysis. J.K. Thackray and A. Sanyal performed B-SOiD analysis. M. Bohic aided with performing injury models. E. Azim, I. Abdus-Saboor, and Y. Yoshida provided feedback and guidance on experimental design and interpretation of results. V.E. Abraira supervised the study. A Upadhyay and M.A. Gradwell wrote the paper with edits from V.E. Abraira. All the authors contributed to its editing.

## STAR+METHODS

### Key Resource table

| REAGENT or RESOURCE | SOURCE | IDENTIFIER |
| --- | --- | --- |
| Antibodies |  |  |
| Rabbit anti-Iba1 (1:1000, IHC) | Abcam | ab178846 |
| Chicken anti-GFAP (1:1000, IHC) | MyBioSource | MBS555403 |
| Chicken anti-GFP (1:1000, IHC) | Aves | GFP 1020 |
| Rabbit anti-GFP (1:1000, IHC) | Invitrogen | A11122 |
| Mouse anti-NeuN (1:2000, IHC) | Millipore | MAB377 |
| Rabbit anti-Neuropeptide Y (1:1000, IHC) | Peninsula Laboratories | T-4070.0050 |
| Rabbit anti-dsRed (1:2000, IHC) | Takara | 632496 |
| Rabbit anti-cfos (1:1000, IHC) | Synaptic Systems | 226003 |
| Goat anti-CTb (1:5000, IHC) | List Biological Labs | 703 |
| Mouse anti-VGAT (1:2000, IHC) | Synaptic Systems | 131011 |
| Guinea pig anti-VGLUT1 (1:2000, IHC) | Millipore | AB5905 |
| Rat anti-substance P (1:1000, IHC) | Bio-Rad | 8450-0505 |
| **RNAScope probes** |  |  |
| vGAT | ACD Biosciences | 319191 |
| vGluT2 | ACD Biosciences | 456751 |
| GlyT2 | ACD Biosciences | 409741 |
| **Viruses** |  |  |
| pAAV-hSyn-DIO-EGFP (AAV1) | Addgene | 50457-AAV1 |
| pAAV1-hSyn-DIO-hM4D(Gi)-mCherry (AAV1) | Addgene | 44362-AAV1 |
| pENN.AAV.hSyn.Cre.WPRE.hGH (AAVrg) | Addgene | 105553-AAVrg |
| AAV phSyn1(S)-FLEX-tdTomato-T2A-SypEGFP-WPRE (AAV1) | Addgene | 51509-AAV1 |
| pENN.AAV.CB7.Cl.eGFP.WPRE.rBG (AAV9) | Addgene | 105542-AAV9 |
| pENN.AAV.CB7.Cl.mCherry.WPRE.RBG (AAV9) | Addgene | 105544-AAV9 |
| **Chemicals** |  |  |
| Strychnine hydrochloride | Sigma | S8753 |
| Bicuculline methochloride | Alomone | B-137 |
| CNQX disodium salt | Alomone | C-141 |
| **Experimental Models: Organisms/Strains** |  |  |
| Mouse: *Advillin*<sup>Cre</sup> | Jackson Laboratories | Jax#032536 |
| Mouse: *Advillin*<sup>FlpO</sup> | Zimmerman et. al., 2019 |  |
| Mouse: *Cdx2*<sup>Cre</sup> | Jackson Laboratories | Jax#009350 |
| Mouse: *Emx1*<sup>IRES-Cre</sup> | Jackson Laboratories | Jax#005628 |
| Mouse: *Gad2*<sup>Cherry</sup> | Jackson Laboratories | Jax#023140 |
| Mouse: *Lbx1*<sup>FlpO</sup> | Duan et. al., 2014 |  |
| Mouse: *Mrgprd*<sup>CreER</sup> | Jackson Laboratories | Jax#031286 |
| Mouse: *Ptgfr*<sup>CreER</sup> | Lehnert et. al., 2021 |  |
| Mouse: *Pvalb*<sup>2aCreER</sup> | Jackson Laboratories | Jax#028580 |
| Mouse: *Rbp4*<sup>Cre</sup> | Harris et al., 2019 |  |
| Mouse: *Ret*<sup>CreER</sup> | Luo et. al., 2009 |  |
| Mouse: *Rosa26*<sup>LSL-FSF-tdTomato</sup> | Jackson Laboratories | Ai65, Jax#021875 |
| Mouse: *Rosa26*<sup>LSL-FSF-synaptophysin-GFP</sup> | Jackson Laboratories | RC::FPSit, FPSit<br>Jax#030206 |
| Mouse: *Rosa26*<sup>LSL-SynGFP</sup> | This paper | Derived from<br>Jax#030206 |
| Mouse: *Rosa26*<sup>LSL-synaptophysin-tdTomato</sup> | Jackson Laboratories | Ai34, Jax#012570 |
| Mouse: *Rosa26*<sup>LSL-ChR2-YFP</sup> | Jackson Laboratories | Ai32, Jax#024109 |
| Mouse: *Rosa26*<sup>LSL-FSF-ChR2-YFP</sup> | Jackson Laboratories | Ai80, Jax#025109 |
| Mouse: *Split*<sup>Cre</sup> | Abraira et. al., 2017 |  |
| Mouse: *Nrtk2*<sup>CreER</sup> (aka *Trkb*<sup>CreER</sup>) | Jackson Laboratories | Jax#027214 |
| Mouse: *Nrtk3*<sup>CreER</sup> (aka *Trkc*<sup>CreER</sup>) | Jackson Laboratories | Jax#030291 |
| Mouse: *Trpv1*<sup>Cre</sup> | Jackson Laboratories | Jax#017769 |
| Mouse: *Th*<sup>2aCreER</sup> | Jackson Laboratories | Jax#025614 |
| Mouse: *Vgat*<sup>IRES-Cre</sup> | Jackson Laboratories | Jax#028862 |
| **Software and algorithms** |  |  |
| AxoGraph |  | https://axograph.com/ |
| p-CLAMP 11 | Molecular Devices | https://www.moleculardevices.com/products/axon-patch-clamp-system/acquisition-and-an |
|  |  | alysis-software/pclamp-software-suite |
| ImageJ |  | https://imagej.nih.gov/ij/ |
| IMARIS | Oxford Instruments | https://imaris.oxinst.com/ |
| Adobe Illustrator | Adobe | https://www.adobe.com/ |
| Prism | GraphPad | www.graphpad.com |
| DeepLabCut | Mathis et al., 2018 | https://github.com/DeepLabCut/DeepLabCut |
| PAWS | Jones et al., 2020 | https://github.com/crtwomey/paws |
| B-SOiD | Hsu and Yttri, 2021 | https://github.com/YttriLab/B-SOiD |

## LEAD CONTACT AND MATERIALS AVAILABILITY

Further information and requests for resources should be directed to and will be fulfilled by the Lead Contact, Victoria E. Abraira.

## EXPERIMENTAL MODEL AND SUBJECT DETAILS

Mouse lines used to target DRG neuron populations include *Advillin^Cre^* or *Advillin^FlpO^* for all DRG neurons^152,153^, *TH^2aCreER^* for C-LTMRs^61^, *TrkB^CreER^* for AD-LTMRs^154^, *Ret^CreER^* for Aβ RA-LTMRs^69,155^, *TrkC^CreER^* for Aβ SA1-LTMRs^127^, *PTGFR^CreER^* for Aβ Field-LTMRs^68^, *PV^2aCreER^* for proprioceptors^69^, *Mrgprd^CreER^*for nonpeptidergic nociceptors^156^, and *Trpv1^Cre^* for peptidergic nociceptors^157^. Mouse lines used to target the spinal cord include *Cdx2^Cre^* for CNS tissue below spinal segment C2^158^, and *Lbx1^FlpO^* for dorsal horn neurons^159^. Mouse lines used to target inhibitory interneuron populations include *Vgat^IRES-Cre^* and *Gad2^Cherry^*for GABAergic inhibitory neurons^160,161^. *Emx1^IRES-Cre^* and *Rbp4^Cre^* were used to target cortical neurons^162,163^. For visualization or manipulation of Cre and/or Flp-dependent populations, we utilized the following mouse lines: *_Rosa26_LSL-FSF-tdTomato* _(Ai65)_^1^6^4^_, *Rosa26*_*LSL-FSF-synaptophysin-GFP* _(FPSit)_^1^6^5^_, R*osa*26_LSL-SynGFP _(LSL-SynGFP, derived from FPSit_^1^6^5^_), *Rosa26*_*LSL-*synaptophysin-tdTomato _(Ai34), *Rosa26*_LSL-ChR2-YFP _(Ai32)_^1^6^6^_, and *Rosa26*_LSL-FSF-ChR2-YFP _(Ai80)_^1^6^7^. Transgenic mouse strains were used and maintained on a mixed genetic background (129/C57BL/6). Experimental animals used were of both sexes. With the exception of *in-vivo* recordings, housing, surgery, behavioral experiments and euthanasia were performed in compliance with Rutgers University Institutional Animal Care and Use Committee (IACUC; protocol #201702589). All mice used in experiments were housed in a regular light cycle room (lights on from 08:00 to 20:00) with food and water available ad libitum. *In-vivo* recordings were performed at the lab of Dr. Eiman Azim according to the Animal Care and Use Committee (IACUC; protocol #16-00011) at the Salk Institute. Cortical injections were performed at the lab of Dr. Yutaka Yoshida according to the Animal Care and Use Committee (IACUC; protocol #2018-0045) at the Burke Neurological Institute. All mice used in experiments were housed in a regular light cycle room (lights on from 07:00 to 19:00) with food and water available ad libitum.

### METHOD DETAILS

### Genetic crosses and statistical methods related to individual figures

#### Genetic crosses and statistical methods related to **Figure 1**

**(A)** Primary afferents were labeled using *Advillin^Cre^* mice, where Cre expression is restricted to the DRG^152^, crossed with an *Rosa26^LSL-Synaptophysin-GFP^* mouse derived from the FPSit line^165^, which allows for Cre-dependent visualization of synaptic terminals. **(B)** Spinal afferents were labeled using a genetic intersection of *Cdx2^Cre^*, where Cre expression was restricted to the caudal nervous system^158^, and the *Lbx1^FlpO^* line, where Flp expression is restricted to the spinal dorsal horn^159^. These mice were crossed to the FPSit reporter line^165^ to allow for Cre- and Flp-dependent visualization of spinal synaptic terminals. **(F,G)** Quantification of excitatory and inhibitory markers in the Gr core (F) and shell (G) normalized to total DAPI. The Gr core had a significantly higher number of VGlut2+ cells compared to vGAT+ or GlyT2+ cells. N=4 mice per group, One-way ANOVA, Tukey’s multiple comparisons, VGlut2 vs. VGAT **p=0.0049, VGlut2 vs Glyt2 **p=0.001, VGAT vs Glyt2 p=0.5127. The Gr shell had a more even distribution of excitatory and inhibitory neurons. N=4 mice per group, One-way ANOVA, Tukey’s multiple comparisons, VGlut2 vs. VGAT p=0.9655, VGlut2 vs Glyt2 p=0.2565, VGAT vs Glyt2 p=0.3583. Error bars represent SEM. **(K,L)** Quantification of Gr VPL-PNs and GAD2+ inhibitory neurons in the core (H) and shell (I). The Gr core had similar numbers of VPL-PNs and inhibitory neurons. N=3 mice per group, unpaired t-test, p=0.6502. The Gr shell had significantly more inhibitory neurons than VPL-PNs. N=3 mice per group, unpaired t-test, ***p=0.0005. Error bars represent SEM.

#### Statistical methods related to Figure 2

**(A)** To label VPL-PNs, we injected ∼420 nL of pENN.AAV.CB7.CI.eGFP.WPRE.rBG (AAV9) into the VPL, and used *Gad2^Cherry^* mice to visualize inhibitory neurons^161^. **(C)** Characterization of sAP incidence: n = 91 VPL-PNs from 9 mice, and n = 82 inhibitory neurons from 6 GAD2^Cherry^ mice. **(D)** Characterization of sAP discharge frequency: n = 81 VPL-PNs from 9 mice, and n = 32 inhibitory neurons from 6 GAD2^Cherry^ mice. VPL-PNs exhibit higher sAP discharge frequency: Mann-Whitney test, ****p<0.0001. **(E-H)** Characterization of Gr VPL-PN and Gr inhibitory neurons response to depolarizing current injection: n = 32 VPL-PNs from 5 mice, and n = 41 inhibitory neurons from 6 GAD2^Cherry^ mice. **(F)** VPL-PNs fire more APs at 10 pA, 20 pA, and 30 pA current injection steps: Mixed-effects model with Geisser-Greenhouse correction and Sidak’s multiple comparisons test, 10 pA **p = 0.0014; 20 pA **p=0.0024; 30 pA *p=0.0190; 40 pA, p=0.0763; 50 pA, p=0.3036; 60 pA, p=0.8957; 70 pA, p=0.9952; 80pA, p>0.9999; 90pA, p>0.9999; 100pA, p=0.9997; 110pA, p=0.9872; 120pA, p=0.9114. **(G)**, lower rheobase: Mann-Whitney test, **p = 0.0026 **(H)**, and a more hyperpolarized AP threshold: Unpaired T-test, ****p<0.0001. **(K-N)** Characterization of sEPSCs: n = 79 VPL-PNs from 10 mice, and n = 47 inhibitory neurons from 7 GAD2^Cherry^ mice. **(K)** VPL-PNs receive higher frequency sEPSC input: Mann-Whitney test, ***p=0.0005, **(L)** with larger amplitude: Mann-Whitney test, ****p<0.0001, **(M)** similar rise time: Mann-Whitney test, p=0.4038, **(N)** and similar decay: Mann-Whitney test, p=0.2976.

#### Genetic crosses and statistical methods related to Figure 3

**(A)** To activate primary afferent inputs onto VPL-PNs, we generated *Advillin^Cre^*;*Rosa26*^LSL-ChR2-YFP^(Ai32)^166^ mice, where ChR2 expression was restricted to primary afferent terminals, and injected ∼420 nL of pENN.AAV.CB7.CI.mCherry.WPRE.RBG into the VPL to visualize Gr VPL-PNs. To activate spinal inputs onto VPL-PNs, we generated *Cdx2^Cre^*;*Lbx1^FlpO^*;*Rosa26*^LSL-FSF-CatCh-eYFP^(Ai80)^168^ mice, where a calcium translocating channelrhodopsin (CatCh) was expressed in spinal afferent terminals in the Gr, and injected ∼420 nL of pENN.AAV.CB7.CI.mCherry.WPRE.RBG into the VPL. To activate primary afferent or spinal inputs onto Gr inhibitory neurons, we generated *Advillin^Cre^*;Ai32;*Gad2^Cherry^* mice to activate primary afferents or *Cdx2^Cre^*;*Lbx1^FlpO^*;Ai80;*Gad2^Cherry^*mice to activate spinal inputs onto fluorescently labeled inhibitory neurons^161^.

**(B)** Amplitude of monosynaptic responses to photostimulation of primary afferent terminals (*Advillin^Cre^*;Ai32) recorded in 29 VPL-PNs from 5 mice; 32 inhibitory neurons from 4 mice, and spinal projection terminals (Cdx2^Cre^;Lbx1^FlpO^;Ai80) recorded in 47 VPL-PNs from 5 mice; 8 inhibitory neurons from 2 mice. VPL-PNs receive larger input than inhibitory neurons from primary afferents: Mann-Whitney test, ***p = 0.0005, and spinal projections: Mann-Whitney test, **p = 0.0039. **(J)** To activate inhibitory inputs onto VPL-PNs, we generated *Vgat^IRES-Cre^*;*Rosa26*^LSL-ChR2-YFP^(Ai32)^166^ mice, where ChR2 expression was restricted to inhibitory terminals, and injected ∼420 nL of pENN.AAV.CB7.CI.mCherry.WPRE.RBG into the VPL to visualize Gr VPL-PNs. **(K-L)** Optically-evoked IPSCs were recorded from 34 VPL-PNs in 6 mice. Bicuculline (10 µM) was applied first in 6 recordings, strychnine (1 µM) was applied first in 7 recordings, and bicuculline and strychnine were sequentially applied in 11 recordings from 5 mice. One-Way ANOVA with Dunnett’s multiple comparisons test: Control vs. Bic, p=0.4456, Control vs. Strych, ****p<0.0001, Control vs. BS, ****p<0.0001. **(N-R)** Spontaneous IPSCs were recorded from 32 VPL-PNs in 6 mice. Bicuculline (10 µM) was applied first in 6 recordings, strychnine (1 µM) was applied first in 8 recordings, and bicuculline and strychnine were sequentially applied in 9 recordings from 6 mice. **(O)** Normalized sIPSC Frequency: Kruskal-Wallis with Dunn’s multiple comparisons test; Control vs. Bic, p=0.2388, Control vs. Strych, **p=0.0056, Control vs. BS, ****p<0.0001. **(P)** Normalized sIPSC amplitude: Kruskal-Wallis with Dunn’s multiple comparisons test; Control vs. Bic, **p=0.0078, Control vs. Strych, ****p<0.0001. **(R)** Normalized sIPSC Decay: One-way ANOVA with Dunnett’s multiple comparisons test, with single pooled variance; Control vs. Bic, p=0.0538, Control vs. Strych, ****p<0.0001.

#### Statistical methods related to Figure 4

**(A,B)** To restrict DREADD expression to VPL-PNs we injected a retrograde Cre-virus (pENN.AAV.hSyn.Cre.WPRE.hGH AAVrg) into the VPL, and injected a Cre-dependent hM4Di DREADD virus (pAAV1-hSyn-DIO-hM4D(Gi)-mCherry AAV1) into the Gr. **(C)** CNO-treated mice exhibited a decreased tactile sensitivity to von Frey filaments compared to saline-treated mice. N=12 mice, 2-way ANOVA with repeated measures, Sidak’s multiple comparisons, 0.4 VF ***p=0.0008, 0.6 VF *p=0.0101, 1 VF *p=0.0346. Error bars represent SEM. **(D)** CNO mice exhibited a decreased average brush score compared to saline mice. N=12 mice, Wilcoxon test, *p=0.0469. **(E,F)** To restrict DREADD expression to Gr inhibitory neurons, we injected a Cre-dependent hM4Di DREADD virus (pAAV1-hSyn-DIO-hM4D(Gi)-mCherry) into the Gr of *Vgat^IRES-Cre^* mice.

**(G)** CNO-treated mice and SNI mice had significantly increased von Frey sensitivity compared to saline controls. N=8 mice, 2-way ANOVA with repeated measures, Tukey’s multiple comparisons, *=CNO vs. Saline, 0.04 VF ***p=0.0002, 0.07 VF **p=0.0039, 0.16 VF *p=0.0148, 0.4 VF ***p=0.0001, ^#^=SNI vs. Saline, 0.008 VF ^#^p=0.0266, 0.04 VF ^##^p=0.0011, 0.07 VF ^#^p=0.0336, 0.16 VF ^###^p=0.0005, 0.4 VF ^##^p=0.0039, 0.6 VF ^##^p=0.0062. Error bars represent SEM. **(H)** CNO mice had significantly increased average brush score compared to saline mice. N=8 mice, Wilcoxon test, *p=0.0156. **(I)** Chemogenetic silencing of Gr inhibitory neurons induced a conditioned place aversion. N=8 mice, paired t-test, **p=0.003. **(K)** Comparison of PAWS features of SNI or CNO mice compared to saline controls. N=16 mice, symbols indicate how parameters change after either SNI or chemogenetic manipulation compared to control: **↑** = increase, **↓** = decrease, **-** = no change. ↑ = p<.05, ↑↑ = p<.005. See **Supplemental Table 1,2** for statistics. **(L)** Usage of the shaking module by SNI and CNO mice compared to saline controls. N=23 mice (23 saline videos, 16 CNO videos, and 15 SNI videos), Kruskal-Wallis test, Dunn’s multiple comparisons. There were no differences amongst groups in response to a 0.6 VF stimulus. Saline vs CNO p=0.491, Saline vs SNI p=0.0825, CNO vs SNI p>0.9999. CNO and SNI mice had increased shaking in response to brush stimuli compared to controls. Saline vs CNO **p=0.0014, Saline vs SNI ***p=0.0003, CNO vs SNI p>0.9999. In response to a noxious pinprick, SNI mice had a significantly increased usage compared to saline and CNO mice. Saline vs CNO p>0.9999, Saline vs SNI *p=0.0174, CNO vs SNI *p=0.0215.

#### Statistical methods related to Figure 5

**(B)** Brush stimulation of SNI mice resulted in a significant increase in the number of c-Fos+ cells in the Gr compared to sham controls. N=4 mice per group, Mann-Whitney test, *p=0.0286. **(D)** Evoked Gr spiking activity on the hindpaw ipsilateral to SNI was significantly increased compared to stimulation on the contralateral uninjured side. N=5 mice, paired t-test, **p=0.0048. **(E)** Spontaneous Gr spiking activity was not significantly different between ipsilateral and contralateral sides. N=5 mice, paired t-test, p=0.7773. **(F-P)** Strategy to investigate primary afferent inputs onto VPL-PNs is the same as **Figure 3A**, with the addition of the SNI procedure to induce neuropathic pain. **(G-K)** Primary afferent-evoked AP discharge was recorded in 40 VPL-PNs from 5 sham mice and 38 VPL-PNs from 4 SNI mice. SNI mice exhibited **(I)** increased oAP latency: Mann-Whitney test, **p=0.0040, **(J)** oAP latency SD: Mann-Whitney test, **p=0.0056, and **(K)** AP number per stimulus: Mann-Whitney test, *p=0.0336. **(L-N)** Primary afferent-evoked EPSCs were recorded in 39 VPL-PNs from 5 sham mice and 36 VPL-PNs from 4 SNI mice. **(N)** We found no change in monosynaptic oEPSC amplitude: Mann-Whitney test, p=0.0900. **(P)** Paired-pulse ratio was examined in 21 VPL-PNs from 4 sham mice and 29 VPL-PNs from 4 SNI mice. SNI exhibited reduced paired-pulse ratio: Unpaired t-test, **p=0.0076.

#### Statistical methods related to Figure 6

**(A,B)** To restrict DREADD expression to VPL-PNs we injected a retrograde Cre-virus (pENN.AAV.hSyn.Cre.WPRE.hGH AAVrg) into the VPL, and injected a Cre-dependent hM4Di DREADD virus (pAAV1-hSyn-DIO-hM4D(Gi)-mCherry AAV1) into the Gr, after which mice received an SNI procedure to induce neuropathic pain. **(C)** SNI mice developed hypersensitivity to von Frey stimulation compared to uninjured mice, which was partially rescued by chemogenetic inhibition of VPL-PNs. N=12 mice, 2-way ANOVA with repeated measures, Tukey’s multiple comparisons, *=Uninjured+Saline vs SNI+Saline, 0.008 VF *p=0.0174, 0.04 VF ****p<0.0001,.0.07 VF ****p<.0.0001, 0.16 VF ****p<.0.0001, 0.4 VF ***p=0.0002, 0.6 VF ***p=0.0001. ^#^=SNI+Saline vs SNI+CNO, 0.04 VF ^###^p=0.0004, 0.07 VF ^###^p=0.0003, 0.16 VF ^###^p=0.0001, 0.4 VF ^##^p=0.0041. ^†^=Uninjuired+Saline vs SNI+CNO, 0.6 VF ^††^p=0.0014. **(D)** Chemogenetic inhibition of VPL-PNs during SNI did not significantly affect average brush score compared to saline-treated mice. N=12 mice, Wilcoxon test, p=0.5688. **(E,F)** We generated *Vgat^IRES-Cre^*;Ai32 mice to optogenetically activate inhibitory terminals in the Gr. **(G)** Optogenetic activation of Gr inhibition using continuous blue light stimulation was more effective than 20 Hz blue light stimulation at reducing hypersensitivity to von Frey stimulation in SNI mice compared to a control continuous orange light stimulation. N=10 mice, Two-way ANOVA with repeated measures, Tukey’s multiple comparisons. *=Orange cont vs Blue cont, 0.008 VF **p=0.001,0.02 VF ****p<0.0001, 0.04 VF ***p=0.0002, 0.07 VF ****p<0.0001, 0.16 VF ****p<0.0001. ^#^=Orange cont vs Blue 20Hz, 0.008 VF ^##^p=0.0047, 0.02 VF ^###^p=0.001, 0.04 VF ^###^p=0.005, 0.07 VF ^##^p=0.004, 0.16 VF ^#^p=0.0109. ^†^=Blue cont vs Blue 20Hz, 0.07 VF ^†††^p=0.0009, 0.16 VF ^†^p=0.0255, 0.4 VF ^†^p=0.0221. **(H)** Blue light stimulation did not significantly affect average brush score compared to orange light stimulation in SNI mice. N=8 mice, Wilcoxon test, p=0.1563.**(I)** Blue light stimulation induced a conditioned place preference 1 hour post conditioning in SNI mice. N=7 mice, Friedman’s test, Dunn’s multiple comparisons, pretest vs real-time p=0.1153, pretest vs 1 hr post **p=0.0019, pretest vs 24 hrs post p=0.0518. **(K)** Comparison of PAWS features of blue light stimulation versus orange light stimulation in SNI mice. **↑** = increase, **↓** = decrease, **-** = no change. **↓** = p<.05. See **Supplemental Table 3** for statistics. **(L)** Usage of the shaking module in the blue stimulation group compared to orange stimulation. N=10 mice (10 Blue stim videos, 9 orange stim videos), unpaired t-test for each comparison, 0.16 VF *p=0.0103, 0.6 VF **p=0.0056, Brush *p=0.0475, Pinprick p=0.2956.

#### Tamoxifen treatment

Tamoxifen was dissolved in ethanol (20 mg/ml), mixed with an equal volume of sunflower seed oil (Sigma), vortexed for 5-10 min and centrifuged under vacuum for 20-30 min to remove the ethanol. The solution was kept at −80°C and delivered via oral gavage to pregnant females for embryonic or postnatal treatment. For all analyses, the morning after coitus was designated as E0.5 and the day of birth as P0.

#### Immunohistochemistry

Male and female P30-P37 mice were anesthetized with isoflurane and perfused transcardially (using an in-house gravity driven-perfusion system) with heparinized-saline (∼30 sec) followed by 15 minutes of 4% paraformaldehyde (PFA) in PBS at room temperature (RT). Brains were dissected and post-fixed in 4% PFA at 4°C for 2-24 hr. Sections were collected using a vibrating microtome (Leica VT1200S) and processed for immunohistochemistry (IHC) as described previously^169^. When required, a small incision was made to the ventral brainstem to denote the side contralateral to the SNI or sham surgery. Transverse sections (50 um) were taken of the caudal medulla containing the DCN. Free floating sections were rinsed in 50% ethanol (30 min) to increase antibody penetration, followed by three washes in a high salt PBS buffer (HS-PBS), each lasting 10 min. The tissue was then incubated in a cocktail of primary antibodies made in HS- PBS containing 0.3% Triton X-100 (HS-PBSt) for 48 at 4°C. Next, tissue was washed 3 times (10 minutes each) with HS-PBSt, then incubated in a secondary antibody solution made in HS-PBSt for 24 hr at 4°C. Immunostained tissue was mounted on positively charged glass slides (41351253, Worldwide Medical Products) and coverslipped (48393-195, VWR) with Fluoromount-G mounting medium (100241-874, VWR).

### Fluorescence *in situ* hybridization (FISH;RNAscope)

Mice were anesthetized with isoflurane and perfused transcardially as described above. Medullas containing the DCN were dissected and post-fixed in 4% PFA at 4°C for 2-24 hr. Samples were cryosectioned at 20 µm and processed using the RNAscope Multiplex Fluorescent v2 Assay (Advanced Cell Diagnostics, 323110). Tissue was placed in 1xPB for 5 mins at room temperature then dehydrated through successive EtOH steps (50%, 70%, 100%, 100% – 5 min) and then dried at room temperature. Tissue was then treated with hydrogen peroxide for 10 min at RT, washed in water, treated with target retrieval reagents (322000) in a steamer for 5 min, and then treated with 100% EtOH for 3 min at RT. Tissue was then treated with protease III (322337) for 8 min at 40°C before being rinsed with water. Probes for Slc32a1/vGAT (Mm-Slc32a1, 319191), Slc6a5/vGlyT2 (Mm- Slc6a5, 409741), and Slc17a6/vGluT2 (Mm-Slc17a6, 456751) were hybridized for 2 hrs at 40°C in a humidified oven, rinsed in wash buffer, and then stored overnight in 5x saline sodium citrate. After rinsing with wash buffer, a series of incubations was performed to amplify and develop hybridized probe signals. Briefly, AMP1 (323101), AMP2 (323102), and AMP3 (323103) were successively applied for 30 min, 30 min, and 15 min at 40°C in a humidified oven, respectively, with a buffer wash in-between. For each channel, HRPC1 (323104) and/or HRPC2 (323105) and/or HRPC3 (323106) were applied for 15 min at 40°C in a humidified oven followed by a buffer wash, then treated using TSA Plus Fluorophores (PerkinElmer; NEL741001KT, NEL744001KT, and NEL745001KT), followed by 15 min of HRP blocker (323107) at 40°C in a humidified oven. Slides were then mounted with Fluoromount-G mounting medium (100241-874, VWR) with DAPI.

### Image acquisition and analysis

Images were captured with a Zeiss LSM 800 confocal or a Zeiss axiovert 200M fluorescence microscope. Images for cell counts were taken with 10x or 20x air objectives, and images of synaptic contacts were taken using a 40x oil objective. ImageJ (cell count plug-in) was used for colocalization analysis of cell bodies. Imaris (spot detection plug-in) was used for synaptic analysis, quantification of synaptic terminals, and generation of contour density plots.

### C-fos induction

Mice received either a SNI surgery to induce neuropathic pain, or a sham surgery as a control. 14 days later, mice were anesthetized (5% induction, 1% maintenance) in a supine position with the plantar surface of their hindpaw exposed, and were allowed to rest for 15 minutes. A paintbrush (Artlicious Paint Brushes) was applied heel-to-toe with the following paradigm: 10 brushes within 10 seconds, every 30 seconds, for 30 minutes. 1 hour later, mice were perfused and processed for immunohistochemistry. Transverse DCN sections (∼50 um) were taken and stained with rabbit anti-cFos (Synaptic Systems, 226003), mouse anti-NeuN (Millipore MAB377), and guinea pig anti-vGluT1 (Millipore AB5905). 20x images of the Gr were acquired using the Zeiss LSM 800 confocal microscope, with vGlut1 immunostaining used to identify the Gr boundaries. 4 images were analyzed per condition. c-Fos+ cells were counted, normalized to total NeuN+ cells per image.

### CTb injections

#### Hindpaw

Mice were anesthetized (5% induction, 2-3% maintenance). Injections of 1% unconjugated CTb (∼7µL across four sites) were injected into the ventral paw of C57BL/6 wild type mice to label paw sensory input. Animals were perfused 5 days after CTb injection and tissue was processed for IHC as described above. Free floating sections were immunostained with goat-CTb (list-labs). 5x images of CTb-labeled afferent inputs were taken at the Zeiss LSM 800 confocal microscope and ImageJ was used for synaptic analysis.

### Stereotaxic injections

#### Ventral posterolateral thalamus (VPL)

Mice were anesthetized with isoflurane (5% induction, 1.5-2% maintenance) and placed onto a stereotaxic frame (Stoelting) that had a feedback controlled heating blanket maintained at 36°C (FHC) on the base. The scalp was cleaned with Betadine (Purdue Products) followed by 70% ethanol (Fisher) three times. Bupivacaine (0.03 mg/kg) was injected subcutaneously into the scalp, and Meloxicam (5 mg/kg) was injected subcutaneously into the flank. A midline incision was made and the skull was exposed. The skull was leveled in the dorsoventral plane by ensuring equal bregma and lambda coordinates. Craniotomy and Injections were made following coordinates with respect to bregma: anteroposterior (AP)= -1.5 mm, mediolateral (ML)= +1.8 mm, dorsoventral (DV)= -3.5 mm. For retrograde labeling of VPL-PNS for electrophysiology and anatomical experiments, 420 nL of either pENN.AAV.CB7.CI.eGFP.WPRE.rBG (AAV9) or pENN.AAV.CB7.CI.mCherry.WPRE.RBG (AAV9) was injected into the contralateral thalamus. For retrograde targeting of VPL-PNs for chemogenetic manipulation, 420 nL of pENN.AAV.hSyn.Cre.WPRE.hGH (AAVrg) was injected into the contralateral thalamus. For all VPL injections, the micropipette was lowered down to the target depth and allowed to sit for 8 minutes. Virus was pressure injected over 10 minutes. 210 nL of the virus was injected at DV= -3.5 mm, then the micropipette was raised to DV= -3.3 mm, where the remaining 210 nL of the virus was injected. This was followed by a 8 minute period of rest to allow for diffusion of the virus into the surrounding tissue. The micropipette was then slowly raised out of the tissue. Following injections, the overlying muscle and the skin incision was sutured, and Ethiqa XR (3.25 mg/kg) was injected subcutaneously into the flank. Mice were allowed to recover from anesthesia on a heating pad, and were returned to their home cages. 2-4 weeks following the injection, mice were either perfused, and their tissue was processed for immunohistochemistry, or they were utilized for electrophysiological or behavioral experiments.

#### Cortex (M1, S1-Forelimb, S1-Hindlimb)

Following the procedures described above, a craniotomy and injection were made at the following coordinates with respect to bregma: For <u>M1</u>: AP= +1.0 mm, ML= +1.7 mm, DV= -1.0 mm. For <u>S1-forelimb</u> (S1-FL): AP= 0 mm, ML= +2.0 mm, DV= -1.0 mm. For <u>S1-hindlimb</u> (S1-HL): AP= -0.5 mm, ML= +1.4, DV= -1.0 mm. For each target, 60-100 nL of AAV-phSyn1(S)-FLEX-tdTomato-T2A-SypEGFP-WPRE (Addgene, 51509) was injected at the designated coordinates. Virus was pressure injected over 10 minutes, and was allowed to rest for 8 minutes to allow for diffusion of the virus into the surrounding tissue.

#### Gracile nucleus (Gr)

Following the procedures described above, the skin and underlying muscle surrounding the base of the skull was removed. For greater consistency of Gr injections, AP and ML coordinates were taken with respect to the caudal base of the skull, while DV coordinates were taken with respect to the cerebellar surface. A craniotomy was made at the following coordinates relative to the midline at the caudal base of the skull: AP= +0.3 mm, ML= -0.15 mm, revealing the caudal surface of the cerebellum. The micropipette was lowered to the following coordinates: AP= +0.3 mm, ML= -0.15 mm, DV= -2.00 mm and allowed to rest for 5 minutes. Injections of 25 nL each were made at four DV coordinates: DV= -2.00 mm, -1.95 mm, -1.90 mm, -1.85 mm, with the micropipette slowly being raised dorsally over the course of injection. Virus was pressure injected over 6 minutes, and allowed to rest for 8 minutes to allow for diffusion of the virus into the surrounding tissue. For targeting of retrogradely-labeled VPL-PNs for chemogenetics, 100 nL of pAAV1-hSyn-DIO-hM4D(Gi)-mCherry (AAV1) was injected into the ipsilateral Gr, alongside injection of 420 nL of pENN.AAV.hSyn.Cre.WPRE.hGH (AAVrg) into the contralateral VPL (described above). For chemogenetic targeting of local Gr inhibitory neurons, 100nL of pAAV1-hSyn-DIO-hM4D(Gi)-mCherry was injected into the Gr of *Vgat^IRES-Cre^* mice.

### Optic probe implantation

#### Gracile nucleus (Gr)

The skull was positioned at a 20 degree decline to gain better optical access to the brainstem. The skin and underlying muscle surrounding the entire skull was removed. Connective tissue was removed from the skull by gentle scraping and a blue light curable bonding agent (iBond, Heraeus Kulzer) was applied to the skull to provide a better surface for dental composite to bind to. A dental drill was used to remove the caudal occipital bone, revealing the caudal cerebellum. To gain optical access to the DCN, a needle was used to puncture the dura surrounding the cerebellum. A small portion of the caudal base of the cerebellum was aspirated to reveal the Gr. An optic cannula (ThorLabs, CFML52U-20, 4 mm length, Ø200 µm Core, 0.50 NA) was lowered to the following coordinates with respect to the obex: AP= 0 mm, ML= -0.4 mm, DV= +0.5 mm. Surrounding muscle was sutured around the cannula, and tissue glue was used to secure the optic cannula in place. Dental composite (Tetric Evoflow, Ivoclar Vivadent) was then applied to the optic cannula as well as the skull surface. Skin was sutured around the dental composite and stabilized with tissue glue. This procedure was done in *Vgat^RES-Cre^*;Ai32 mice.

### Spared nerve injury (SNI)

To study the function of the DCN in mediating allodynia and hyperalgesia, we utilized the spared nerve injury (SNI) procedure to induce chronic neuropathic pain^170^. Mice were anesthetized and the hair surrounding the hindlimb was shaved. An incision was made on the right left below the knee, and another incision was made in the muscle to reveal the sciatic nerve. A ligature using silk suture (Braintree Scientific, SUT-S 103) was made around the peroneal and tibial nerve branches, leaving the sural nerve intact. A 2 cm portion of the peroneal/tibial nerve was cut and removed. Special care was taken to ensure that the sural nerve was not damaged during the procedure. Surrounding musculature was sutured around the nerve, and then the wound was secured using surgical wound clips (Kent Scientific, INS750344-2). Tactile hypersensitivity was determined using von Frey testing, and animals exhibiting any paralysis or desensitization were excluded from experiments.

### Electrophysiology

Recordings were made from male (n = 23) and female (n = 20) mice (10.7 +/- 0.4 wks). Mice used for electrophysiology were all tested for tactile hypersensitivity prior to sacrifice 11-28 days post injury (sham 15.9 +/- 0.9 days and SNI 15.1 +/- 0.8 days); Sham = 13 male and 10 female mice, SNI = 10 male and 10 female mice. Mice were anesthetized with ketamine (100 mg/kg i.p), decapitated, and the lumbar enlargement of the spinal cord rapidly removed in ice-cold sucrose substituted artificial cerebrospinal fluid (sACSF) containing (in mM): 250 sucrose, 25 NaHCO3, 10 glucose, 2.5 KCl, 1 NaH2PO4, 6 MgCl2, and 1 CaCl2. Sagittal or transverse slices (200µm thick) were prepared using a vibrating microtome (Leica VT1200S). Slices were incubated for at least 1 hour at 22-24°C in an interface chamber holding oxygenated ACSF containing (in mM): 118 NaCl, 25 NaHCO3, 10 glucose, 2.5 KCl, 1 NaH2PO4, 1 MgCl2, and 2.5 CaCl2.

Following incubation, slices were transferred to a recording chamber and continually superfused with ACSF bubbled with Carbogen (95% O2 and 5% CO2) to achieve a pH of 7.3-7.4. All recordings were made at room temperature (22-24°C) and neurons visualized using a Zeiss Axiocam 506 color camera. Patch pipettes (3-7 MΩ) were filled with a potassium gluconate-based internal solution containing (in mM): 135 C6H11KO7, 8 NaCl, 10 HEPES, 2 Mg2ATP, 0.3 Na3GTP, and 0.1 EGTA, (pH 7.3, adjusted with KOH, and 300 mOsm) to examine AP discharge and excitatory synaptic transmission. Recordings were acquired in cell-attached (holding current = 0mV), voltage-clamp (holding potential -70mV), or current-clamp (maintained at −60mV) configuration. The incidence of oIPSCs was also assessed using the potassium gluconate-based internal solution by holding the cell at a more depolarized potential in voltage clamp (-20 mV) to reveal hyperpolarizing potentials. A cesium chloride-based internal solution containing (in mM): 130 CsCl, 10 HEPES, 10 EGTA, 1 MgCl2, 2 Na-ATP, 2 Na-GTP, and 5 QX-314 bromide (pH 7.3, adjusted with CsOH, and 280 mOsm) was used to record inhibitory synaptic transmission in all other occasions. All recordings of sIPSCs and oIPSCs were performed in the presence of CNQX (10µM). No liquid junction potential correction was made, although this value was calculated at 14.7 mV (22°C). Slices were illuminated using an X-CITE 120LED Boost High-Power LED Illumination System that allowed visualization of mCherry (VPL-PNs, GAD2+ inhibitory neurons) with TRITC filters, and visualization of GFP fluorescence (VPL-PNs) and ChR2 photostimulation using FITC filter set.

AP discharge patterns were assessed in the current clamp from a membrane potential of −60 mV by delivering a series of depolarizing current steps (1 s duration, 20 pA increments, 0.1 Hz), with rheobase defined as the first current step that induced AP discharge. Spontaneous action potential discharge was assessed using cell-attached voltage clamp recordings, with minimal holding current (< 20 pA).

For optogenetic circuit mapping, photostimulation intensity was suprathreshold (24 mW), duration 1 ms (controlled by transistor-transistor logic pulses), 0.1 Hz. This ensured generation of action potential discharge in presynaptic populations and allowed confident assessment of postsynaptic currents in recorded neurons. When examining paired pulse ratio, an interval of 200 ms was chosen for oEPSCs and 1 s for oIPSCs.

All data were amplified using a MultiClamp 700B amplifier, digitized online (sampled at 20 kHz, filtered at 5 kHz) using an Axon Digidata 1550B, acquired and stored using Clampex software. After obtaining the whole-cell recording configuration, series resistance, input resistance, and membrane capacitance were calculated (averaged response to -5mV step, 20 trials, holding potential -70mV). A series resistance of <30 MΩ was set as a requirement for inclusion in data analysis. This resulted in the exclusion of one VPL-PN recording from a sham mouse, four inhibitory neuron recordings from sham mice, and one inhibitory neuron recording from an SNI mouse.

### Electrophysiology Analysis

All data were analyzed offline using AxoGraph X software.

APs were captured using a derivative threshold with AP threshold defined as the inflection point during spike initiation (dV/dt ≥15 V/s). AP latency was measured as the time from depolarization onset to AP threshold, AP amplitude was measured as the difference between peak amplitude and threshold, AP rise time (10-90% of AP peak), AP decay time (90-10% of AP peak), AP half-width (50% of AP peak), AHP amplitude as the difference between AP threshold and AHP peak, and AHP half-width (50% of AHP peak).

Optogenetically-evoked excitatory postsynaptic currents (oEPSCs) and inhibitory postsynaptic currents (oIPSCs) were measured from baseline, just before photostimulation. The peak amplitude of responses was calculated from the average of 10 successive trials. Several parameters were considered for determining whether a photostimulation-evoked synaptic input was monosynaptic or polysynaptic. The latency of oPSCs was measured as the time from photostimulation to the onset of the evoked current. The “jitter” in latency was measured as the standard deviation in latency of 10 successive trials. Importantly, the latency of monosynaptic inputs was shorter, there was minimal jitter in the onset of responses between trials, and reliability (percentage of photostimulation trials to evoke a response) was higher than those deemed polysynaptic inputs. Paired pulse ratio was determined by dividing the peak amplitude of the second pulse response by the peak amplitude of the first pulse response. The number of evoked APs was calculated as the number of APs within the first 20 ms following photostimulation. The latency and jitter of evoked APs was calculated from the first evoked AP following photostimulation.

sEPSCs and sIPSCs were detected using a sliding template method (a semi-automated procedure in the Axograph package). Frequency was determined from at least 30s of recording at a membrane potential of -70 mV. Peak amplitude, rise time (10-90% of peak) were measured from captured events, and decay time constant (fitted over 10-90% of the decay phase) was measured from the averaged sEPSC/sIPSC for each neuron. When measuring the contributions of different neurotransmitters to sIPSC frequency, sIPSC and oIPSC amplitude, and sIPSC decay all drug responses were normalized to control conditions.

### In-vivo recordings

#### Animal preparation and electrode implantation

C57BL/6J mice (Jax #000664) were given SNI (described above) and sent to the Salk Institute. Mice were then anesthetized with isoflurane and placed in a stereotaxic frame (Kopf Instruments) in a flat skull position. The skin and overlying fascia were incised and retracted, and the skull was cleaned with cotton swabs and dried. A small burr hole was drilled above the cerebellar vermis and a 127 µm silver wire (A-M Systems) was lowered ∼1 mm below the surface of the brain and cemented in place to serve as the reference electrode for differential recordings. Next, a headplate was affixed to the skull immediately above bregma with epoxy (Tetric EvoFlow, Ivoclar Vivadent). Mice were then placed in the stereotaxic frame equipped with custom-made ear bars to allow the head to be dorsiflexed 30° from horizontal. From this position, an incision was made at the back of the head and the skin and overlying muscles were retracted to expose the obex. The most caudal portion of the occipital bone was removed to permit access to more rostral aspects of the Gr. Finally, the dura was pierced with a 30½ gauge syringe and retracted to facilitate electrode penetration. The animal was then taken off isoflurane and administered a cocktail containing 1.2 mg/kg urethane and 20 mg/kg xylazine^171^, and transferred to a separate recording rig. The headplate was clamped in place the head was again dorsiflexed ∼30° from horizontal to enable clear access to the obex and the dorsal column nuclei. The surface of the brainstem was kept moist with a layer of sterile saline or paraffin oil. Throughout recordings, animals were kept on a DC-powered heating pad to maintain stable body temperature, and supplemental doses of urethane/xylazine were given to maintain animals in an areflexic state, assessed by foot pinch.

#### Tactile stimuli

For all experiments, the hindlimbs were positioned on rolled gauze pads such that the glabrous footpads were oriented upwards allowing easy access for applying tactile stimuli. Tactile stimuli were applied with 2 small paintbrushes that were oriented 180° from one another and rotated across the surface of the footpad by means of a stepper motor (28BYJ-48m, MikroElektronika) controlled by a microcontroller board (Arduino UNO, Arduino). The board was triggered by a TTL pulse from a Cerebus neural signal processor (Blackrock Microsystems) and was programmed to rotate the brush-wheel at a rate of 60° per sec. There was a 7 sec interval between each complete revolution (1 revolution = 2 tactile stimuli). The duration of each individual brush stimulus was ∼0.7 sec and the duration of a full revolution was 6 sec. The entire stepper motor / brush assembly was attached to a 3-axis micromanipulator (M3301; WPI) that enabled the tactile stimuli to be applied precisely to the footpad. For each recording site within the Gr, responses to a total of 20 tactile stimuli were assessed (10 revolutions x 2 brushstrokes / rev). Stimuli were randomly alternated between the right and left footpad throughout the recording session.

#### Extracellular Recordings

Extracellular recordings were made using carbon fiber electrodes (∼0.8 MOhm, Carbostar-1, Kation Scientific) mounted to a stereotaxic manipulator (Kopf Instruments) and connected to a Cereplex M headstage (Blackrock Microsystems), which interfaced with the Cerebus neural signal processor. Signals were sampled at 30 kHz and band-pass filtered between 250-5000 Hz. Hindlimb responsive neurons were found within the main body of the Gr (measured relative to obex; A/P = -0.25 à +0.2 mm; M/L = 0.0 à ± 0.3mm; D/V= -0.05 à -0.20mm from the surface of the brainstem). The electrode was slowly lowered as tactile inputs were applied to the footpad and responses were monitored on an oscilloscope and through audio feedback. Robust tactile responses were typically found between 50 and 200 µm below the surface of the brainstem. All data were collected within a 4-6 hour recording session and mice were perfused for histological analysis immediately following the experiment. In vivo electrophysiology data consisted of continuous raw data, spiking events (defined as crossing a threshold set well above noise), and analog signals marking motor outputs. Cerebus Central Suite (Blackrock Microsystems) was used to set event thresholds, and NeuroExplorer (Plexon) was used to export recording files to Excel or MATLAB (MathWorks) for further analysis. To quantify tactile evoked responses to brushstrokes applied to the hindlimbs, peri-event raster plots were generated and thresholded spikes were binned into 100 msec intervals, beginning at the initiation of motor revolution, and continuing for 6 sec, encompassing the delivery of 2 brushstroke stimuli. To separate tactile evoked responses from inter-stimulus interval (ISI) spiking, each 6 sec revolution was separated into 2 individual 3.0 sec intervals corresponding to a half revolution of the stepper motor. Thus, a single tactile stimulus always occurred at some point within each 3.0 sec interval. Within each 3.0 second interval, we defined the window of tactile stimulation as a 0.7 sec window centered on bins where the spike frequency rose to at least 200% above baseline activity, which was usually quite low. Background spiking activity was also computed for each recording site by summing the total number of spikes within a 4 second window centered in the 7 second interval between wheel revolutions. The process of computing tactile-evoked activity and background activity was repeated for each recording site. The number of spikes generated at each recording site across all 20 tactile events (10 revolutions x 2 stimuli) were then combined. The evoked and background activity were then summed across all units (both ipsilateral and contralateral to the lesioned sciatic nerve) for each mouse and paired t-tests were used to compare evoked and background activity between ipsilateral and contralateral Gr.

### Behavioral testing

7-14 weeks old male and female mice from a Bl6 (Jax#000664), CD1 (Charles River Strain#22), or mixed background were used for behavior testing .The experimenter was blind to the genotype of the animals. For all behavior assays, at least 7 days prior testing mice were transferred to the holding area adjacent to the behavior rooms. On the day of training/testing, mice were transferred to the behavior room for a 30-minute habituation.

### Behavior test #1: VonFrey

Von frey testing was used to assay tactile sensitivity across various stimulation strengths^67^. The testing protocol was adapted from Murthy et al., 2018^172^. 11 different von Frey filaments, ranging from .008g to 4g, were applied to the plantar hindpaw 4 times each. Withdrawal responses were recorded out of 4 total stimulations.

Responses to the lowest force were recorded first before moving on to the next highest force.

### Behavior test #2: Brush

Dynamic brush testing was used to assay tactile sensitivity at baseline as well as dynamic allodynia during neuropathic pain^173^. A paintbrush (Artlicious Paint Brushes) was applied to the glabrous hindpaw from heel to toe 9 times, and responses were scored from 0-3: 0=no response, 1=withdrawal, 2=guarding, 3=licking/tending. Responses were analyzed as the sum of all scores divided by 9 total stimulations, with a 5 minute break after every 3 trials.

### Behavior test #3: Pinprick

Pinprick testing was used to assess noxious mechanical sensitivity at baseline and mechanical hyperalgesia during neuropathic pain^67^. An insect pin (Austerlitz Premium Insect Entomology Dissection Pins, Size 1) was slowly raised until it made contact with the glabrous hindpaw. The number of withdrawal responses to pinprick were recorded over 10 simulations, with a 5 minute break in the middle.

### Behavior test #4: Hargreaves

Hargreaves testing was used to assess thermal sensitivity at baseline and thermal hyperalgesia during neuropathic pain^174^. Animals were enclosed over a glass plate, allowing for optical access to the hindpaw. A radiant heat source (IITC) was positioned underneath the animal, and aimed at the glabrous surface of the hindpaw. Intensity of the heat source was calibrated as to produce a withdrawal latency of ∼10-15 seconds in naive animals, as to allow for a window to capture both hypo and hyperalgesia, with a cutoff time of 20 seconds to avoid long term skin damage^67^. Latency to withdrawal (seconds) to the radiant heat stimulus was recorded, and averaged over 3 trials, with at least 5 minutes of rest between trials.

### Behavior test #5: Conditioned place preference/aversion (CPP/CPA)

To assess general preference/avoidance behavior of behavioral manipulations in the absence of any evoked sensory responses, I utilized the two-chamber CPP/CPA assay^175^. For chemogenetic experiments, mice were allowed to freely explore both chambers on Day 1 for 10 minutes to establish initial preference. On Days 2-5, chambers were closed off, and mice were conditioned to saline on one side in the morning, and CNO on the other side in the evening, for 30 minutes each. To remove bias of time of day, conditioning to saline/CNO was swapped between morning and evening sessions every day. On Day 6, mice were once again allowed to freely explore both chambers to establish CPP/CPA. For optogenetic experiments, mice were allowed to freely explore the chambers in the morning on Day 1 for 10 minutes to establish initial preference. Following this initial session, mice were optically stimulated with continuous light on the side they preferred less, using a 2 second on, 2 second off paradigm. This first session was used to establish “real-time’ placed preference. This was repeated in the evening on Day 1, as well as a morning and evening session on Day 2, for a total of 4 sessions. On Days 2 and 3, mice were allowed to freely explore the chambers without any optical stimulation to determine if conditioning was retained 1 hour and 24 hours after the final stimulating session.

### Behavior test #5: Data generation for PAWS and B-SOiD

To generate data for PAWS analysis, mice were placed in clear plexiglass chambers positioned perpendicular to a camera (FASTCAM Mini AX100, ZEISS Milvus 100mm f/2M ZF.2 lens). Barriers were placed so that animals were not able to see each other. Mice were habituated for a minimum of 3 days prior to the testing date, and habituated in the chambers for 30 minutes prior to testing. A maximum of two different simulations were applied per day, a minimum of 6 hours apart, to limit the effects of prior stimulations on future responses. Stimulations were also delivered in order from most innocuous to most noxious (0.16 VF → 0.6 VF → brush → pinprick). Mouse behaviors were recorded at 4000 FPS, and video durations were approximately 1.25 seconds. The camera was placed at a 10 degree angle facing downwards towards the animal, approximately 3 feet away from the plexiglass chambers. Infrared lights were used to illuminate the scene.

Videos were processed by DeepLabCut^101^ to predict the location of the toe, metatarsophalangeal join (MTP), heel, and reference positions from the plexiglass box (top left corner and bottom left corner). We utilized DeepLabCut for markerless pose estimation of body parts. The software provides an open-source toolbox that facilitates the extraction of frames, annotation of body parts, training of a deep neural network, and subsequent tracking of body parts using the trained model. The network is finetuned on frames extracted from videos and the accompanying labels marking the locations of the body parts, training the architecture end to end. We extracted 20 frames from a random sample of our collected videos through the DeepLabCut toolbox and annotated the toe, MTP, and the top and the bottom corners on the left side of the box containing the mouse.

We observed the greatest confidence of tracking for the MTP, which we then used for all analyses. The annotated frames were then used to train the pose estimation network and track the body parts on unlabeled frames through the full length of our collected videos. Labeled data was then run through the PAWS^102^ software for feature extraction.

### Analysis of PAWS data

Paw withdrawal tracking data was analyzed using the Pain Assessment at Withdrawal Speeds (PAWS) software^102^. Coordinates for the metatarsophalangeal joint (MTP) were used to extract features of the paw withdrawal, including max height, distance traveled, max velocity, # of shakes, and guarding duration. Data points under 80% tracking confidence were excluded from the analysis, and a rolling average filter was applied for data smoothing.

#### B-SOiD analysis

To quantify behavior following foot stimulation, pose estimation data output by DeepLabCut was imported into the unsupervised behavioral discovery algorithm, B-SOiD, following previously described methods^105^. Briefly, data was first converted from pixel units to millimeter units, and downsampled to 1/14th of the original frame rate. Additionally, two static reference points for each video were computed: a) the initial position of the toe, and b) the toe position at the maximum elevation reached post-stimulation. The coordinates of the toe, MTP, and two reference points were selected for feature extraction, non-linear embedding into UMAP space.

HDBSCAN (parameters: min_samples=10; minimum cluster size range=0.1%-1.0%) was used to identify nine behavioral clusters. Cluster assignments were used to train a random forest classifier to predict behavioral clusters from extracted features, and this model was run on the full dataset. Behavioral clusters were given post-hoc ethological labels by expert raters. Clusters representing less than 1% of all frames were excluded from further analysis. The normalized number of frames labeled by each cluster starting from the initial withdrawal response was computed and plotted using matplotlib.

### Statistical analysis

All data are reported as mean values ± the standard error of the mean (SEM), unless otherwise stated. Statistical analysis was performed in GraphPad Prism (USA) or Matlab. All data was tested for normality using the Shapiro-Wilk test. For all tests *p < 0.05, **p < 0.05, ***p < 0.005, and ****p≤ 0.0005. The means of different data distributions were compared using One-way ANOVA, Tukey’s multiple comparisons (Figure 1F, 1G), Unpaired t-test (Figure 1K, 1L, 2H, 5P, 6L), Mann-Whitney test (Figure 2D, 2G, 2K, 2L, 2M, 2N, 3C, 5B, 5I, 5J, 5K, 5N), Mixed-effects model with Geisser-Greenhouse correction and Sidak’s multiple comparisons test (Figure 2F), One-Way ANOVA with Dunnett’s multiple comparisons test (Figure 3L, 3R), Kruskal-Wallis with Dunn’s multiple comparisons test (Figure 3O, 3P, 4L), Two-way ANOVA with Sidaks multiple comparisons test (Figure 4C), Wilcoxon test (Figure 4D, 4H, 6D, 6H), 2-way ANOVA with Tukey’s multiple comparisons test (Figure 4G, 6C, 6G), Paired t-test (Figure 4I, 5D, 5E), and Friedman’s with Dunn’s multiple comparisons test (Figure 6I). Statistics details for supplemental figures can be found in supplemental figure legends.

## DATA AND CODE AVAILABILITY

Data are available upon request from the Lead Contact, Victoria E. Abraira.

## REFERENCES

1. Lolignier, S., Eijkelkamp, N. & Wood, J. N. Mechanical allodynia. Pflugers Arch. 467, 133–139 (2015).

2. Jensen, T. S. & Finnerup, N. B. Allodynia and hyperalgesia in neuropathic pain: clinical manifestations and mechanisms. Lancet Neurol. 13, 924–935 (2014).

3. Kuner, R. Central mechanisms of pathological pain. Nat. Med. 16, (2010).

4. Gold, M. S. & Gebhart, G. F. Nociceptor sensitization in pain pathogenesis. Nat. Med. 16, 1248 (2010).

5. Gangadharan, V. et al. Neuropathic pain caused by miswiring and abnormal end organ targeting. Nature 606, 137–145 (2022).

6. Boyle, K. A. et al. Defining a Spinal Microcircuit that Gates Myelinated Afferent Input: Implications for Tactile Allodynia. Cell Rep. 28, (2019).

7. Peirs, C. et al. Mechanical allodynia circuitry in the dorsal horn is defined by the nature of the injury. Neuron 109, 73 (2021).

8. Sun, L. et al. Parabrachial nucleus circuit governs neuropathic pain-like behavior. Nat. Commun. 11, 1–21 (2020).

9. Wang, Z. & Xu, Z.-Z. The Parabrachial Nucleus as a Key Regulator of Neuropathic Pain. Neurosci. Bull. 37, 1079 (2021).

10. Miyamoto, K., Kume, K. & Ohsawa, M. Role of microglia in mechanical allodynia in the anterior cingulate cortex. J. Pharmacol. Sci. 134, (2017).

11. Mansikka, H. & Pertovaara, A. Supraspinal influence on hindlimb withdrawal thresholds and mustard oil-induced secondary allodynia in rats. Brain Res. Bull. 42, 359–365 (1997).

12. Sung, B. et al. Supraspinal involvement in the production of mechanical allodynia by spinal nerve injury in rats. Neurosci. Lett. 246, 117–119 (1998).

13. Malan, T. P. et al. Extraterritorial neuropathic pain correlates with multisegmental elevation of spinal dynorphin in nerve-injured rats. Pain 86, 185–194 (2000).

14. Bian, D., Ossipov, M. H., Zhong, C., Malan, T. P. & Porreca, F. Tactile allodynia, but not thermal hyperalgesia, of the hindlimbs is blocked by spinal transection in rats with nerve injury. Neurosci. Lett. 241, (1998).

15. Shankarappa, S. A. et al. Prolonged nerve blockade delays the onset of neuropathic pain. Proc. Natl. Acad. Sci. U. S. A. 109, 17555–17560 (2012).

16. Delhaye, B. P., Long, K. H. & Bensmaia, S. J. Neural Basis of Touch and Proprioception in Primate Cortex. Compr. Physiol. 8, 1575–1602 (2018).

17. Loutit, A. J., Vickery, R. M. & Potas, J. R. Functional organization and connectivity of the dorsal column nuclei complex reveals a sensorimotor integration and distribution hub. J. Comp. Neurol. 529, 187–220 (2021).

18. Sun, H. et al. Nerve injury-induced tactile allodynia is mediated via ascending spinal dorsal column projections. Pain 90, 105–111 (2001).

19. Gmel, G. E. et al. Postsynaptic dorsal column pathway activation during spinal cord stimulation in patients with chronic pain. Front. Neurosci. 17, 1297814 (2023).

20. Schwark, H. D. & Ilyinsky, O. B. Inflammatory pain reduces correlated activity in the dorsal column nuclei. Brain Res. 889, 295–302 (2001).

21. Kim, H. Y., Wang, J. & Gwak, Y. S. Gracile neurons contribute to the maintenance of neuropathic pain in peripheral and central neuropathic models. J. Neurotrauma 29, 2587–2592 (2012).

22. Miki, K. et al. Responses of dorsal column nuclei neurons in rats with experimental mononeuropathy. Pain 76, 407–415 (1998).

23. Shortland, P. & Molander, C. The time-course of abeta-evoked c-fos expression in neurons of the dorsal horn and gracile nucleus after peripheral nerve injury. Brain Res. 810, 288–293 (1998).

24. Persson, J. K., Hongpaisan, J. & Molander, C. c-fos expression in gracilothalamic tract neurons after electrical stimulation of the injured sciatic nerve in the adult rat. Somatosens. Mot. Res. 10, 475–483 (1993).

25. Tsuda, M. Microglia in the spinal cord and neuropathic pain. J. Diabetes Investig. 7, 17–26 (2016).

26. Magnussen, C., Hung, S.-P. & Ribeiro-da-Silva, A. Novel expression pattern of neuropeptide Y immunoreactivity in the peripheral nervous system in a rat model of neuropathic pain. Mol. Pain 11, (2015).

27. Zou, Y. et al. Distinct calcitonin gene-related peptide expression pattern in primary afferents contribute to different neuropathic symptoms following chronic constriction or crush injuries to the rat sciatic nerve. Mol. Pain 12, 1744806916681566 (2016).

28. Hughes, D. I., Scott, D. T., Riddell, J. S. & Todd, A. J. Upregulation of substance P in low-threshold myelinated afferents is not required for tactile allodynia in the chronic constriction injury and spinal nerve ligation models. J. Neurosci. 27, 2035–2044 (2007).

29. Upregulation of the GABA transporter GAT-1 in the gracile nucleus in the spared nerve injury model of neuropathic pain. Neurosci. Lett. 480, 132–137 (2010).

30. Liljencrantz, J. & Olausson, H. Tactile C fibers and their contributions to pleasant sensations and to tactile allodynia. Front. Behav. Neurosci. 8, 69962 (2014).

31. Tashima, R. et al. Optogenetic Activation of Non-Nociceptive Aβ Fibers Induces Neuropathic Pain-Like Sensory and Emotional Behaviors after Nerve Injury in Rats. eNeuro 5, (2018).

32. Dhandapani, R. et al. Control of mechanical pain hypersensitivity in mice through ligand-targeted photoablation of TrkB-positive sensory neurons. Nat. Commun. 9, 1640 (2018).

33. Gautam, M. et al. Distinct Local and Global Functions of Aβ Low-Threshold Mechanoreceptors in Mechanical Pain Transmission. Research square (2023) doi:10.21203/rs.3.rs-2939309/v1.

34. Suresh, A. K. et al. Sensory computations in the cuneate nucleus of macaques. Proc. Natl. Acad. Sci. U. S. A. 118, (2021).

35. Versteeg, C., Rosenow, J. M., Bensmaia, S. J. & Miller, L. E. Encoding of limb state by single neurons in the cuneate nucleus of awake monkeys. J. Neurophysiol. 126, 693–706 (2021).

36. Versteeg, C., Chowdhury, R. H. & Miller, L. E. Cuneate nucleus: The somatosensory gateway to the brain. Curr Opin Physiol 20, 206–215 (2021).

37. Turecek, J., Lehnert, B. P. & Ginty, D. D. The encoding of touch by somatotopically aligned dorsal column subdivisions. Nature 612, 310–315 (2022).

38. Marino, J., Canedo, A. & Aguilar, J. Sensorimotor cortical influences on cuneate nucleus rhythmic activity in the anesthetized cat. Neuroscience 95, 657–673 (2000).

39. Andersen, P., Eccles, J. C., Oshima, T. & Schmidt, R. F. MECHANISMS OF SYNAPTIC TRANSMISSION IN THE CUNEATE NUCLEUS. J. Neurophysiol. 27, 1096–1116 (1964).

40. Canedo, A., Martinez, L. & Mariño, J. Tonic and bursting activity in the cuneate nucleus of the chloralose-anesthetized cat. Neuroscience 84, 603–617 (1998).

41. Dykes, R. W., Rasmusson, D. D., Sretavan, D. & Rehman, N. B. Submodality segregation and receptive-field sequences in cuneate, gracile, and external cuneate nuclei of the cat. J. Neurophysiol. 47, 389–416 (1982).

42. Mariño, J., Martinez, L. & Canedo, A. Sensorimotor Integration at the Dorsal Column Nuclei. News Physiol. Sci. 14, 231–237 (1999).

43. Conner, J. M. et al. Modulation of tactile feedback for the execution of dexterous movement. Science 374, 316–323 (2021).

44. Guo, D. & Hu, J. Spinal presynaptic inhibition in pain control. Neuroscience 283, 95–106 (2014).

45. Sivilotti, L. & Woolf, C. J. The contribution of GABAA and glycine receptors to central sensitization: disinhibition and touch-evoked allodynia in the spinal cord. J. Neurophysiol. 72, 169–179 (1994).

46. Roberts, L. A., Beyer, C. & Komisaruk, B. R. Nociceptive responses to altered GABAergic activity at the spinal cord. Life Sci. 39, 1667–1674 (1986).

47. Enna, S. J. & McCarson, K. E. The role of GABA in the mediation and perception of pain. Adv. Pharmacol.

48. Comitato, A. & Bardoni, R. Presynaptic Inhibition of Pain and Touch in the Spinal Cord: From Receptors to Circuits. Int. J. Mol. Sci. 22, (2021).

49. Baba, H. et al. Removal of GABAergic inhibition facilitates polysynaptic A fiber-mediated excitatory transmission to the superficial spinal dorsal horn. Mol. Cell. Neurosci. 24, 818–830 (2003).

50. Lu, Y. et al. A feed-forward spinal cord glycinergic neural circuit gates mechanical allodynia. J. Clin. Invest. 123, 4050–4062 (2013).

51. Wercberger, R. & Basbaum, A. I. Spinal cord projection neurons: a superficial, and also deep, analysis. Curr Opin Physiol 11, 109–115 (2019).

52. Paixão, S. et al. Identification of Spinal Neurons Contributing to the Dorsal Column Projection Mediating Fine Touch and Corrective Motor Movements. Neuron 104, 749–764.e6 (2019).

53. Browne, T. J. et al. Spinoparabrachial projection neurons form distinct classes in the mouse dorsal horn. Pain 162, 1977–1994 (2021).

54. Todd, A. J. et al. Projection neurons in lamina I of rat spinal cord with the neurokinin 1 receptor are selectively innervated by substance p-containing afferents and respond to noxious stimulation. J. Neurosci. 22, 4103–4113 (2002).

55. Agashkov, K. et al. Distinct mechanisms of signal processing by lamina I spino-parabrachial neurons. Sci. Rep. 9, 19231 (2019).

56. Koch, S. C., Acton, D. & Goulding, M. Spinal Circuits for Touch, Pain, and Itch. Annu. Rev. Physiol. 80, 189–217 (2018).

57. Hughes, D. I. & Todd, A. J. Central Nervous System Targets: Inhibitory Interneurons in the Spinal Cord. Neurotherapeutics 17, 874–885 (2020).

58. Todd, A. J. Identifying functional populations among the interneurons in laminae I-III of the spinal dorsal horn. Mol. Pain 13, 1744806917693003 (2017).

59. Bardoni, R. et al. Pre- and postsynaptic inhibitory control in the spinal cord dorsal horn. Ann. N. Y. Acad. Sci. 1279, 90–96 (2013).

60. Sullivan, S. J. & Sdrulla, A. D. Excitatory and Inhibitory Neurons of the Spinal Cord Superficial Dorsal Horn Diverge in Their Somatosensory Responses and Plasticity in Vivo. J. Neurosci. 42, 1958–1973 (2022).

61. Abraira, V. E. et al. The Cellular and Synaptic Architecture of the Mechanosensory Dorsal Horn. Cell 168, (2017).

62. Moehring, F., Halder, P., Seal, R. P. & Stucky, C. L. Uncovering the Cells and Circuits of Touch in Normal and Pathological Settings. Neuron 100, 349–360 (2018).

63. Tashima, R. et al. A subset of spinal dorsal horn interneurons crucial for gating touch-evoked pain-like behavior. Proc. Natl. Acad. Sci. U. S. A. 118, (2021).

64. Chen, S. et al. A spinal neural circuitry for converting touch to itch sensation. Nat. Commun. 11, 5074 (2020).

65. Foster, E. et al. Targeted ablation, silencing, and activation establish glycinergic dorsal horn neurons as key components of a spinal gate for pain and itch. Neuron 85, 1289–1304 (2015).

66. Escalante, A. & Klein, R. Spinal Inhibitory Ptf1a-Derived Neurons Prevent Self-Generated Itch. Cell Rep. 33, 108422 (2020).

67. Deuis, J. R., Dvorakova, L. S. & Vetter, I. Methods Used to Evaluate Pain Behaviors in Rodents. Front. Mol. Neurosci. 10, 284 (2017).

68. Lehnert, B. P. et al. Mechanoreceptor synapses in the brainstem shape the central representation of touch. Cell 184, 5608–5621.e18 (2021).

69. Niu, J. et al. Modality-based organization of ascending somatosensory axons in the direct dorsal column pathway. J. Neurosci. 33, 17691–17709 (2013).

70. Giuffrida, R. & Rustioni, A. Dorsal root ganglion neurons projecting to the dorsal column nuclei of rats. J. Comp. Neurol. 316, 206–220 (1992).

71. Weisberg, J. A. & Rustioni, A. Differential projections of cortical sensorimotor areas upon the dorsal column nuclei of cats. J. Comp. Neurol. 184, 401–421 (1979).

72. Kim, Y. R. & Kim, S. J. Altered synaptic connections and inhibitory network of the primary somatosensory cortex in chronic pain. Korean J. Physiol. Pharmacol. 26, 69–75 (2022).

73. Seifert, F. & Maihöfner, C. Central mechanisms of experimental and chronic neuropathic pain: findings from functional imaging studies. Cell. Mol. Life Sci. 66, 375–390 (2009).

74. Bak, M. S., Park, H. & Kim, S. K. Neural Plasticity in the Brain during Neuropathic Pain. Biomedicines 9, (2021).

75. Thompson, S. J. et al. Metabolic brain activity suggestive of persistent pain in a rat model of neuropathic pain. Neuroimage 91, 344–352 (2014).

76. Chen, C. et al. Synchronized activity of sensory neurons initiates cortical synchrony in a model of neuropathic pain. Nat. Commun. 14, 689 (2023).

77. Chao, T.-H. H., Chen, J.-H. & Yen, C.-T. Plasticity changes in forebrain activity and functional connectivity during neuropathic pain development in rats with sciatic spared nerve injury. Mol. Brain 11, 55 (2018).

78. Bray, N. Pain: A painful loss of inhibition. Nature reviews. Neuroscience vol. 18 456 (2017).

79. Ab Aziz, C. B. & Ahmad, A. H. The role of the thalamus in modulating pain. Malays. J. Med. Sci. 13, 11–18 (2006).

80. Wang, Z. et al. Altered thalamic neurotransmitters metabolism and functional connectivity during the development of chronic constriction injury induced neuropathic pain. Biol. Res. 53, 36 (2020).

81. Chambers, W. W., Liu, C. N. & McCOUCH, G. P. Inhibition of the dorsal column nuclei. Exp. Neurol. 7, 13–23 (1963).

82. Schwark, H. D., Tennison, C. F., Ilyinsky, O. B. & Fuchs, J. L. Inhibitory influences on receptive field size in the dorsal column nuclei. Exp. Brain Res. 126, 439–442 (1999).

83. Jabbur, S. J. & Banna, N. R. Widespread cutaneous inhibition in dorsal column nuclei. J. Neurophysiol. 33, 616–624 (1970).

84. Andersen, P., Etholm, B. & Gordon, G. Presynaptic and post-synaptic inhibition elicited in the cat’s dorsal column nuclei by mechanical stimulation of skin. J. Physiol. 210, 433–455 (1970).

85. Dawson, G. D. The effect of cortical stimulation on transmission through the cuneate nucleus in the anaesthetized rat. J. Physiol.

86. Valeriani, M., Insola, A. & Mazzone, P. 16. Presynaptic and postsynaptic inhibition in the human dorsal column nuclei. Clin. Neurophysiol. 127, e327 (2016).

87. Chaudhry, F. A. et al. The vesicular GABA transporter, VGAT, localizes to synaptic vesicles in sets of glycinergic as well as GABAergic neurons. J. Neurosci. 18, 9733–9750 (1998).

88. Berkley, K. J. & Hubscher, C. H. Are there separate central nervous system pathways for touch and pain? Nat. Med. 1, 766–773 (1995).

89. Nathan, P. W., Smith, M. C. & Cook, A. W. Sensory effects in man of lesions of the posterior columns and of some other afferent pathways. Brain 109 **( Pt** **5**), 1003–1041 (1986).

90. Vierck, C. J., Jr. Comparison of forelimb and hindlimb motor deficits following dorsal column section in monkeys. Brain Res. 146, 279–294 (1978).

91. Ballermann, M., McKenna, J. & Whishaw, I. Q. A grasp-related deficit in tactile discrimination following dorsal column lesion in the rat. Brain Res. Bull. 54, 237–242 (2001).

92. Houghton, A. K., Hewitt, E. & Westlund, K. N. Dorsal column lesion prevents mechanical hyperalgesia and allodynia in osteotomy model. Pain 82, 73–80 (1999).

93. White, J. C. Neurosurgical treatment of persistent pain. Lancet 2, 161–164 (1950).

94. Condon, L. F. et al. Parabrachial Calca neurons drive nociplasticity. Cell Rep. 43, 114057 (2024).

95. Torsney, C. & MacDermott, A. B. Disinhibition opens the gate to pathological pain signaling in superficial neurokinin 1 receptor-expressing neurons in rat spinal cord. J. Neurosci. 26, 1833–1843 (2006).

96. Sun, S. et al. Leaky Gate Model: Intensity-Dependent Coding of Pain and Itch in the Spinal Cord. Neuron 93, 840–853.e5 (2017).

97. Yaksh, T. L. Behavioral and autonomic correlates of the tactile evoked allodynia produced by spinal glycine inhibition: effects of modulatory receptor systems and excitatory amino acid antagonists. Pain 37, 111–123 (1989).

98. Choi, J.-H. et al. Development of tactile allodynia immediately after spinal anesthesia. Pain Med. 16, 1242–1244 (2015).

99. La, J.-H. & Chung, J. M. Peripheral afferents and spinal inhibitory system in dynamic and static mechanical allodynia. Pain 158, 2285–2289 (2017).

100. Zhang, Z., Hefferan, M. P. & Loomis, C. W. Topical bicuculline to the rat spinal cord induces highly localized allodynia that is mediated by spinal prostaglandins. Pain 92, 351–361 (2001).

101. Mathis, A. et al. DeepLabCut: markerless pose estimation of user-defined body parts with deep learning. Nat. Neurosci. 21, 1281–1289 (2018).

102. Jones, J. M. et al. A machine-vision approach for automated pain measurement at millisecond timescales. Elife 9, (2020).

103. Cichon, J., Sun, L. & Yang, G. Spared Nerve Injury Model of Neuropathic Pain in Mice. Bio Protoc 8, (2018).

104. Hsu, A. I. & Yttri, E. A. B-SOiD, an open-source unsupervised algorithm for identification and fast prediction of behaviors. Nat. Commun. 12, 5188 (2021).

105. Bohic, M. et al. Mapping the neuroethological signatures of pain, analgesia, and recovery in mice. Neuron 111, 2811–2830.e8 (2023).

106. Sun, H. et al. Nerve injury-induced tactile allodynia is mediated via ascending spinal dorsal column projections. Pain 90, 105–111 (2001).

107. Tokunaga, A. et al. Excitability of spinal cord and gracile nucleus neurons in rats with chronically injured sciatic nerve examined by c-fos expression. Brain Res. 847, 321–331 (1999).

108. Pottorf, T. S., Rotterman, T. M., McCallum, W. M., Haley-Johnson, Z. A. & Alvarez, F. J. The Role of Microglia in Neuroinflammation of the Spinal Cord after Peripheral Nerve Injury. Cells 11, (2022).

109. Beggs, S., Trang, T. & Salter, M. W. P2X4R+ microglia drive neuropathic pain. Nat. Neurosci. 15, 1068–1073 (2012).

110. Borst, J. G. & Sakmann, B. Effect of changes in action potential shape on calcium currents and transmitter release in a calyx-type synapse of the rat auditory brainstem. Philos. Trans. R. Soc. Lond. B Biol. Sci. 354, 347–355 (1999).

111. Mendonça, P. R. F. et al. Asynchronous glutamate release is enhanced in low release efficacy synapses and dispersed across the active zone. Nat. Commun. 13, 3497 (2022).

112. Taylor, B. K. Spinal inhibitory neurotransmission in neuropathic pain. Curr. Pain Headache Rep. 13, 208–214 (2009).

113. Vaysse, L. et al. GABAergic pathway in a rat model of chronic neuropathic pain: modulation after intrathecal transplantation of a human neuronal cell line. Neurosci. Res. 69, 111–120 (2011).

114. Jergova, S., Hentall, I. D., Gajavelli, S., Varghese, M. S. & Sagen, J. Intraspinal transplantation of GABAergic neural progenitors attenuates neuropathic pain in rats: a pharmacologic and neurophysiological evaluation. Exp. Neurol. 234, 39–49 (2012).

115. Bráz, J. M. et al. Forebrain GABAergic neuron precursors integrate into adult spinal cord and reduce injury-induced neuropathic pain. Neuron 74, 663–675 (2012).

116. Gwak, Y. S. et al. Activation of spinal GABA receptors attenuates chronic central neuropathic pain after spinal cord injury. J. Neurotrauma 23, 1111–1124 (2006).

117. Hamilton, L. S. et al. Optogenetic activation of an inhibitory network enhances feedforward functional connectivity in auditory cortex. Neuron 80, 1066–1076 (2013).

118. Moon, H. S. et al. Contribution of Excitatory and Inhibitory Neuronal Activity to BOLD fMRI. Cereb. Cortex 31, 4053–4067 (2021).

119. Babl, S. S., Rummell, B. P. & Sigurdsson, T. The Spatial Extent of Optogenetic Silencing in Transgenic Mice Expressing Channelrhodopsin in Inhibitory Interneurons. Cell Rep. 29, 1381–1395.e4 (2019).

120. Gu, L. et al. Pain inhibition by optogenetic activation of specific anterior cingulate cortical neurons. PLoS One 10, e0117746 (2015).

121. Lin, J. Y. A user’s guide to channelrhodopsin variants: features, limitations and future developments. Exp. Physiol. 96, 19–25 (2011).

122. Schoenenberger, P., Gerosa, D. & Oertner, T. G. Temporal control of immediate early gene induction by light. PLoS One 4, e8185 (2009).

123. Ishizuka, T., Kakuda, M., Araki, R. & Yawo, H. Kinetic evaluation of photosensitivity in genetically engineered neurons expressing green algae light-gated channels. Neurosci. Res. 54, 85–94 (2006).

124. Lin, J. Y., Lin, M. Z., Steinbach, P. & Tsien, R. Y. Characterization of engineered channelrhodopsin variants with improved properties and kinetics. Biophys. J. 96, 1803–1814 (2009).

125. Britt, J. P., McDevitt, R. A. & Bonci, A. Use of channelrhodopsin for activation of CNS neurons. Curr. Protoc. Neurosci. **Chapter** 2, Unit2.16 (2012).

126. Sufka, K. J. Conditioned place preference paradigm: a novel approach for analgesic drug assessment against chronic pain. Pain 58, 355–366 (1994).

127. Bai, L. et al. Genetic Identification of an Expansive Mechanoreceptor Sensitive to Skin Stroking. Cell 163, 1783–1795 (2015).

128. Towe, A. L. & Zimmerman, I. D. Peripherally Evoked Cortical Reflex in the Cuneate Nucleus. Nature 194, 1250–1251 (1962).

129. Creed, R. S. & Sherrington, C. S. Observations on concurrent contraction of flexor muscles in the flexion reflex. Proc. R. Soc. Lond. B Biol. Sci. 100, 258–267 (1997).

130. Lundberg, A. Multisensory Control of Spinal Reflex Pathways. in Progress in Brain Research (eds. Granit, R. & Pompeiano, O.) vol. 50 11–28 (Elsevier, 1979).

131. Morgan, M. M. Direct comparison of heat-evoked activity of nociceptive neurons in the dorsal horn with the hindpaw withdrawal reflex in the rat. J. Neurophysiol. 79, 174–180 (1998).

132. Todd, A. J. Neuronal circuitry for pain processing in the dorsal horn. Nat. Rev. Neurosci. 11, 823–836 (2010).

133. Kim, J., Back, S. K., Yoon, Y. W., Hong, S. K. & Na, H. S. Dorsal column lesion reduces mechanical allodynia in the induction, but not the maintenance, phase in spinal hemisected rats. Neurosci. Lett. 379, 218–222 (2005).

134. Sagalajev, B. et al. Absence of paresthesia during high-rate spinal cord stimulation reveals importance of synchrony for sensations evoked by electrical stimulation. Neuron 112, 404–420.e6 (2024).

135. Rankin, G. et al. Nerve injury disrupts temporal processing in the spinal cord dorsal horn through alterations in PV interneurons. Cell Rep. 43, 113718 (2024).

136. Catley, M. J., O’Connell, N. E., Berryman, C., Ayhan, F. F. & Moseley, G. L. Is tactile acuity altered in people with chronic pain? a systematic review and meta-analysis. J. Pain 15, 985–1000 (2014).

137. 137. Soleimani, B., et al. Immunotherapy-Responsive Neuropathic Pain and Allodynia in a Patient With Glycine Receptor Autoantibodies: A Case Report. Neurol Neuroimmunol Neuroinflamm 10, (2023).

138. Lee, K. Y., Ratté, S. & Prescott, S. A. Excitatory neurons are more disinhibited than inhibitory neurons by chloride dysregulation in the spinal dorsal horn. Elife 8, (2019).

139. Gradwell, M. A., Callister, R. J. & Graham, B. A. Reviewing the case for compromised spinal inhibition in neuropathic pain. J. Neural Transm. 127, 481–503 (2020).

140. Guilbaud, G., Peschanski, M., Gautron, M. & Binder, D. Neurones responding to noxious stimulation in VB complex and caudal adjacent regions in the thalamus of the rat. Pain 8, 303–318 (1980).

141. Miki, K. et al. Dorsal column-thalamic pathway is involved in thalamic hyperexcitability following peripheral nerve injury: a lesion study in rats with experimental mononeuropathy. Pain 85, 263–271 (2000).

142. Barbaresi, P. & Mensà, E. Connections from the rat dorsal column nuclei (DCN) to the periaqueductal gray matter (PAG). Neurosci. Res. 109, 35–47 (2016).

143. Burton, H. & Loewy, A. D. Projections to the spinal cord from medullary somatosensory relay nuclei. J. Comp. Neurol. 173, 773–792 (1977).

144. Purves, D., et al. Central Pain Pathways: The Spinothalamic Tract. (Sinauer Associates, 2001).

145. Abraira, V. E. & Ginty, D. D. The sensory neurons of touch. Neuron 79, 618–639 (2013).

146. Kenshalo, D. R. Sensory Functions of the Skin of Humans. (Springer Science & Business Media, 2012).

147. Choi, S. et al. Parallel ascending spinal pathways for affective touch and pain. Nature 587, 258–263 (2020).

148. Ma, W., Peschanski, M. & Besson, J. M. The overlap of spinothalamic and dorsal column nuclei projections in the ventrobasal complex of the rat thalamus: a double anterograde labeling study using light microscopy analysis. J. Comp. Neurol. 245, 531–540 (1986).

149. Lenz, F. A., Weiss, N., Ohara, S., Lawson, C. & Greenspan, J. D. Chapter 6 The role of the thalamus in pain. in Supplements to Clinical Neurophysiology (eds. Hallett, M., Phillips, L. H., Schomer, D. L. & Massey, J. M.) vol. 57 50–61 (Elsevier, 2004).

150. Brüggemann, J., Galhardo, V. & Apkarian, A. V. Immediate reorganization of the rat somatosensory thalamus after partial ligation of sciatic nerve. J. Pain 2, 220–228 (2001).

151. Yan, Y. et al. Thalamocortical Circuit Controls Neuropathic Pain via Up-regulation of HCN2 in the Ventral Posterolateral Thalamus. Neurosci. Bull. 39, 774–792 (2023).

152. Zhou, X. et al. Deletion of PIK3C3/Vps34 in sensory neurons causes rapid neurodegeneration by disrupting the endosomal but not the autophagic pathway. Proc. Natl. Acad. Sci. U. S. A. 107, 9424–9429 (2010).

153. Zimmerman, A. L. et al. Distinct Modes of Presynaptic Inhibition of Cutaneous Afferents and Their Functions in Behavior. Neuron 102, 420–434.e8 (2019).

154. Rutlin, M. et al. The Cellular and Molecular Basis of Direction Selectivity of Aδ-LTMRs. Cell 160, 1027 (2015).

155. Luo, W., Enomoto, H., Rice, F. L., Milbrandt, J. & Ginty, D. D. Molecular identification of rapidly adapting mechanoreceptors and their developmental dependence on ret signaling. Neuron 64, 841–856 (2009).

156. Olson, W. et al. Sparse genetic tracing reveals regionally specific functional organization of mammalian nociceptors. Elife 6, (2017).

157. Cavanaugh, D. J. et al. Trpv1 reporter mice reveal highly restricted brain distribution and functional expression in arteriolar smooth muscle cells. J. Neurosci. 31, 5067–5077 (2011).

158. Hinoi, T. et al. Mouse model of colonic adenoma-carcinoma progression based on somatic Apc inactivation. Cancer Res. 67, 9721–9730 (2007).

159. Bourane, S. et al. Gate control of mechanical itch by a subpopulation of spinal cord interneurons. Science 350, 550–554 (2015).

160. Vong, L. et al. Leptin action on GABAergic neurons prevents obesity and reduces inhibitory tone to POMC neurons. Neuron 71, 142–154 (2011).

161. Peron, S. P., Freeman, J., Iyer, V., Guo, C. & Svoboda, K. A Cellular Resolution Map of Barrel Cortex Activity during Tactile Behavior. Neuron 86, 783–799 (2015).

162. Gorski, J. A. et al. Cortical excitatory neurons and glia, but not GABAergic neurons, are produced in the Emx1-expressing lineage. J. Neurosci. 22, 6309–6314 (2002).

163. Harris, J. A. et al. Hierarchical organization of cortical and thalamic connectivity. Nature 575, 195–202 (2019).

164. Madisen, L. et al. Transgenic mice for intersectional targeting of neural sensors and effectors with high specificity and performance. Neuron 85, 942–958 (2015).

165. Niederkofler, V., et al. Identification of Serotonergic Neuronal Modules that Affect Aggressive Behavior. Cell Rep. 17, 1934–1949 (2016).

166. Madisen, L. et al. A toolbox of Cre-dependent optogenetic transgenic mice for light-induced activation and silencing. Nat. Neurosci. 15, 793–802 (2012).

167. Daigle, T. L. et al. A Suite of Transgenic Driver and Reporter Mouse Lines with Enhanced Brain-Cell-Type Targeting and Functionality. Cell 174, 465–480.e22 (2018).

168. Kleinlogel, S. et al. Ultra light-sensitive and fast neuronal activation with the Ca^2^+-permeable channelrhodopsin CatCh. Nat. Neurosci. 14, 513–518 (2011).

169. Hughes, D. I. et al. Morphological, neurochemical and electrophysiological features of parvalbumin-expressing cells: a likely source of axo-axonic inputs in the mouse spinal dorsal horn. J. Physiol. 590, 3927–3951 (2012).

170. Decosterd, I. & Woolf, C. J. Spared nerve injury: an animal model of persistent peripheral neuropathic pain. Pain 87, 149–158 (2000).

171. Lee, C. & Jones, T. A. Effects of Ketamine Compared with Urethane Anesthesia on Vestibular Sensory Evoked Potentials and Systemic Physiology in Mice. J. Am. Assoc. Lab. Anim. Sci. 57, 268–277 (2018).

172. Murthy, S. E. et al. The mechanosensitive ion channel Piezo2 mediates sensitivity to mechanical pain in mice. Sci. Transl. Med. 10, (2018).

173. Cruccu, G. & Truini, A. Tools for assessing neuropathic pain. PLoS Med. 6, e1000045 (2009).

174. Hargreaves, K., Dubner, R., Brown, F., Flores, C. & Joris, J. A new and sensitive method for measuring thermal nociception in cutaneous hyperalgesia. Pain 32, 77–88 (1988).

175. Prus, A. J., James, J. R. & Rosecrans, J. A. Conditioned Place Preference. (CRC Press/Taylor & Francis, 2009).

