## Supplemental Figure 1 for "The Dorsal Column Nuclei Scale Mechanical Sensitivity in Naive and Neuropathic Pain States"

Supplemental Figure 1 - related to Figure 1. The Gr receives ascending input primarily from Aβ-LTMR's and spinal projections

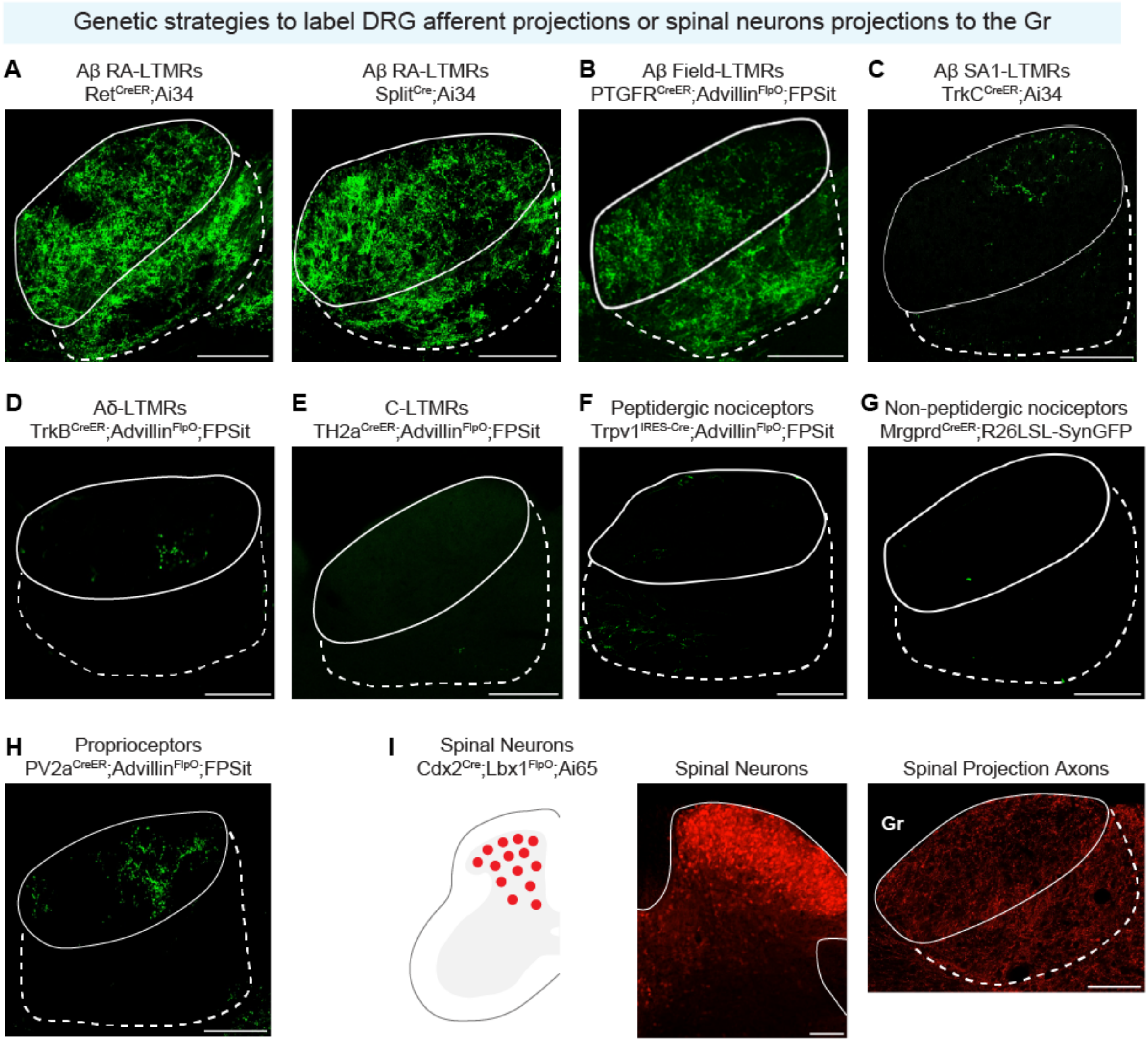
