## Supplemental Figure 2 for "The Dorsal Column Nuclei Scale Mechanical Sensitivity in Naive and Neuropathic Pain States"

Supplemental Figure 2 - related to Figure 1. The Gr receives descending cortical input from S1 hindlimb

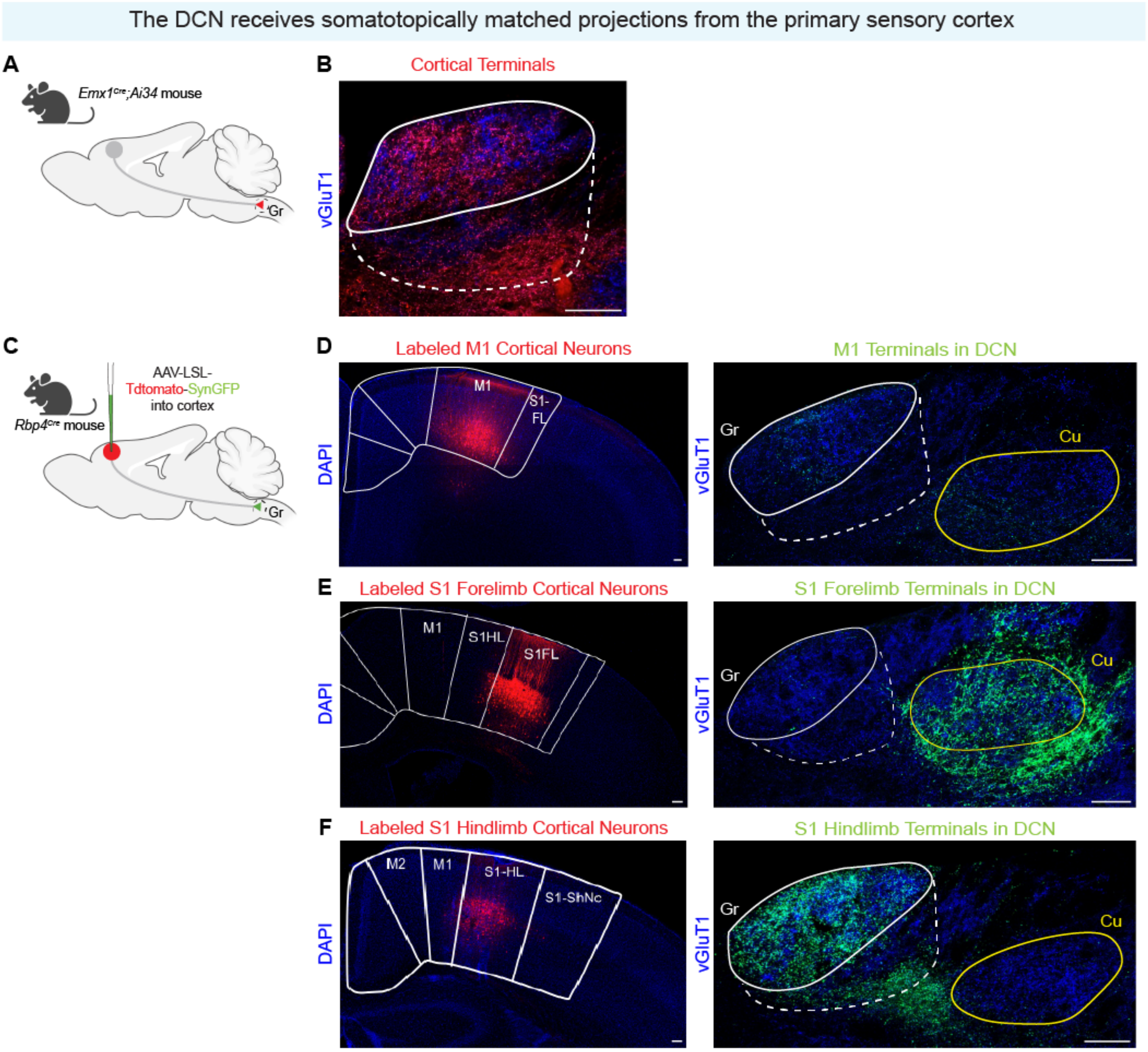
