## Supplemental Figure 3 for "The Dorsal Column Nuclei Scale Mechanical Sensitivity in Naive and Neuropathic Pain States"

Supplemental Figure 3 - related to Figure 3. Primary afferent and cortical terminals receive axoaxonic contacts from inhibitory neurons

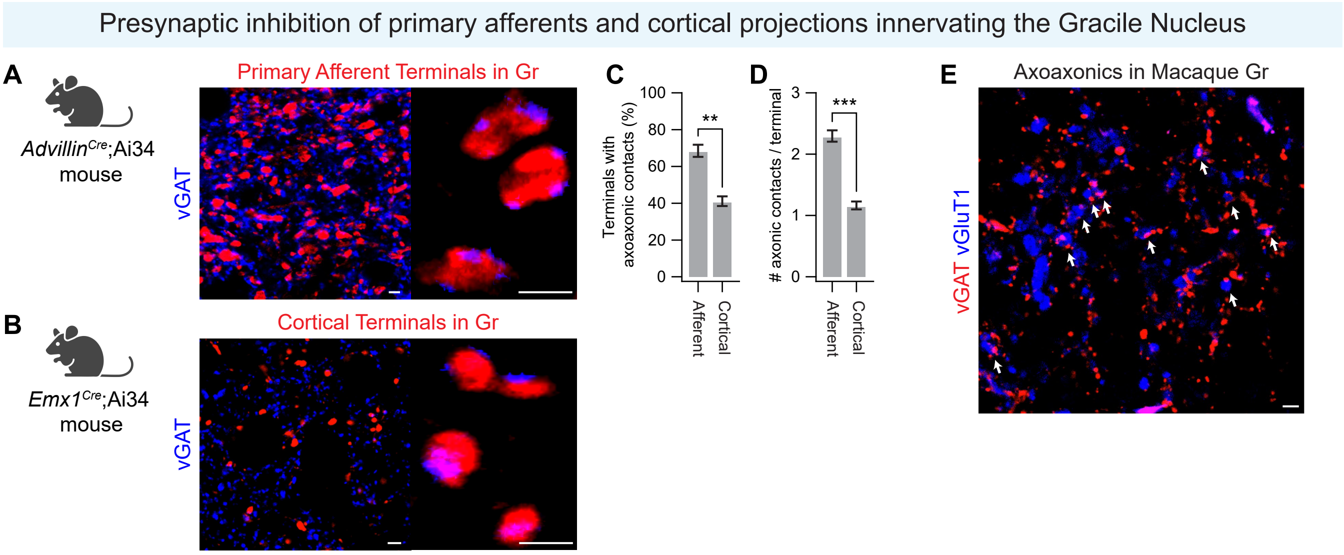
