## Supplemental Figure 4 for "The Dorsal Column Nuclei Scale Mechanical Sensitivity in Naive and Neuropathic Pain States"

Supplemental Figure 4 - related to Figure 4. Silencing VPL-PNs or inhibitory neurons does not affect noxious mechanical or thermal sensitivity, and CNO alone does not affect mechanical or thermal sensitivity or place preference

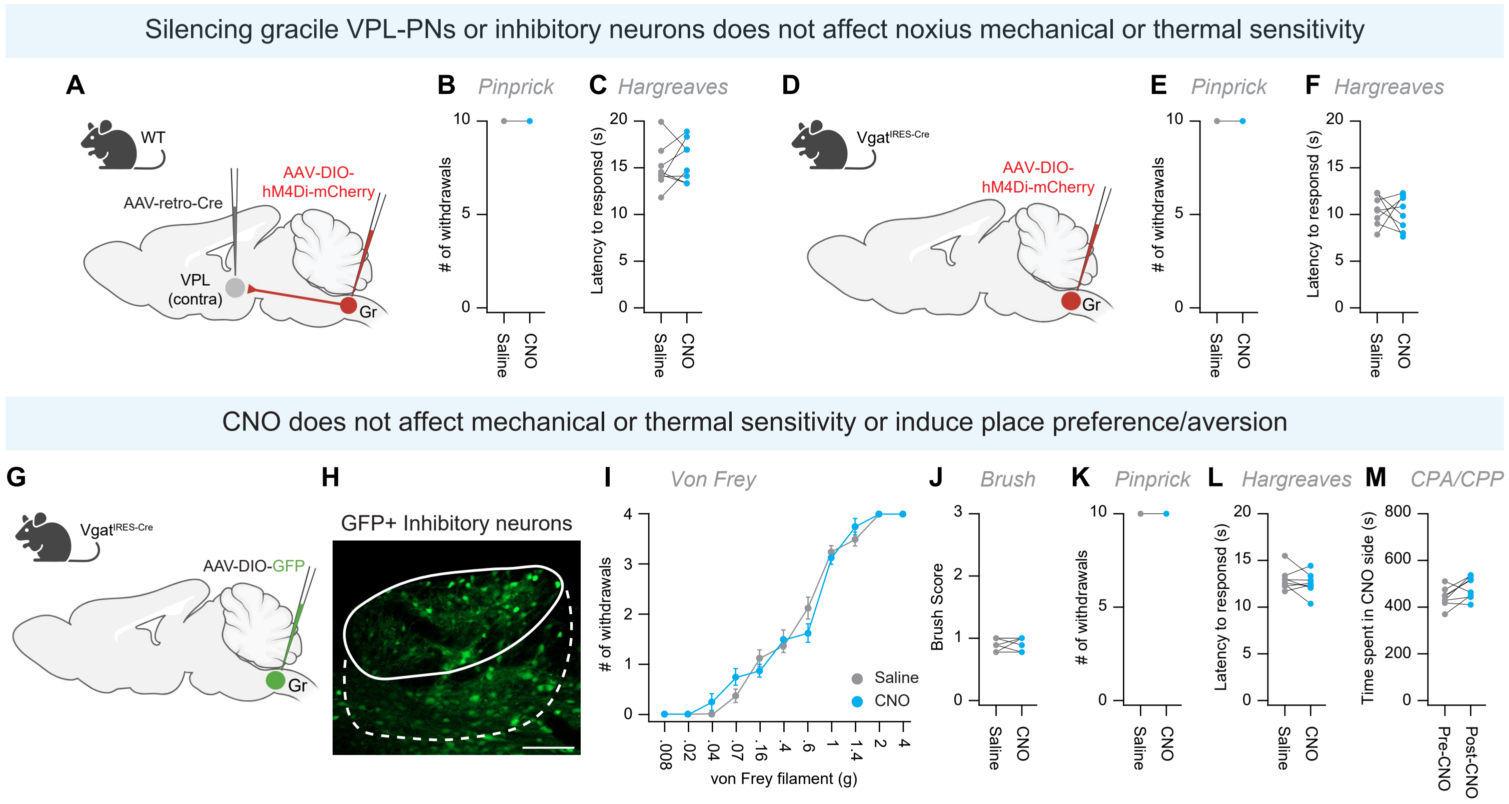
