## Supplemental Figure 5 for "The Dorsal Column Nuclei Scale Mechanical Sensitivity in Naive and Neuropathic Pain States"

Supplemental Figure 5 - related to Figure 5. Neuropathic injury evokes robust microglia activation in the Gr

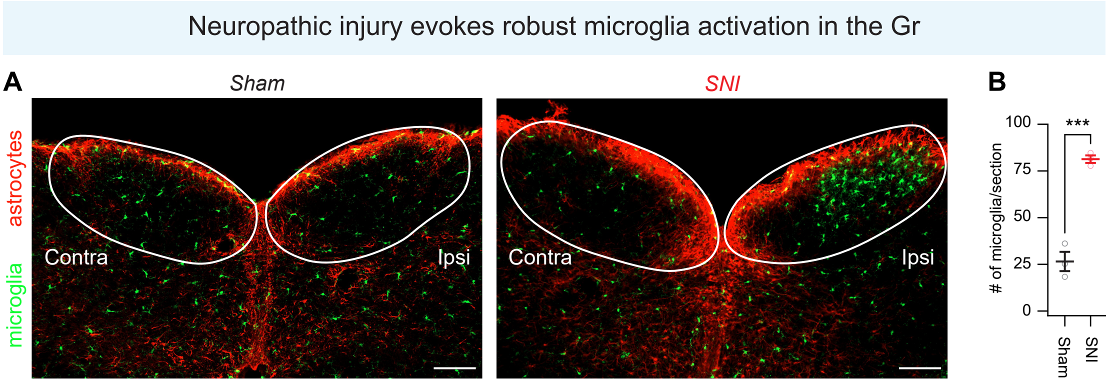
