## Supplemental Figure 6 for "The Dorsal Column Nuclei Scale Mechanical Sensitivity in Naive and Neuropathic Pain States"

Supplemental Figure 6 - related to Figure 5. VPL-PN and inhibitory neuron electrophysiological properties remain largely unchanged by neuropathic injury

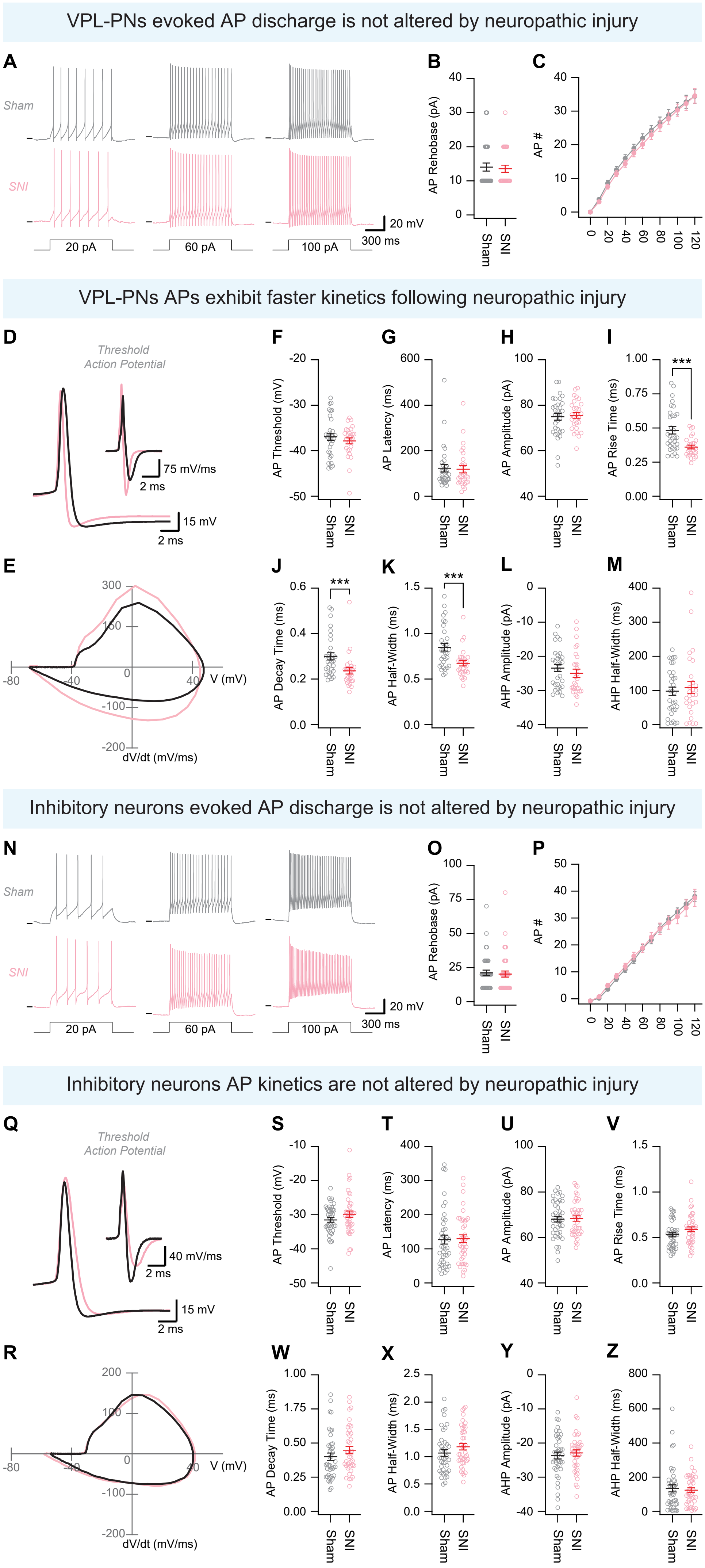
