## Supplemental Figure 7 for "The Dorsal Column Nuclei Scale Mechanical Sensitivity in Naive and Neuropathic Pain States"

Supplemental Figure 7 - related to Figure 5. VPL-PN and inhibitory neuron spontaneous activity and excitatory drive is altered by neuropathic injury

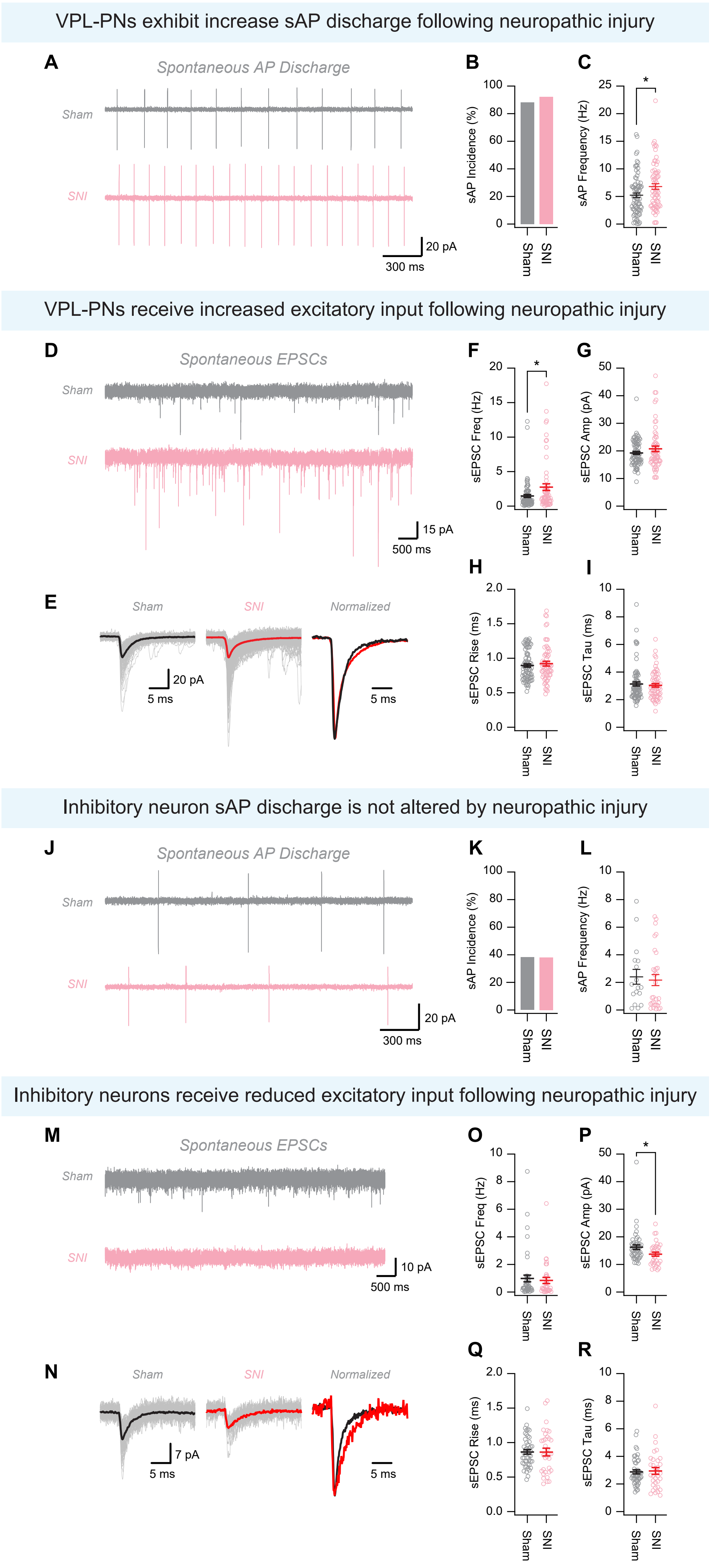
