## Supplemental Figure 8 for "The Dorsal Column Nuclei Scale Mechanical Sensitivity in Naive and Neuropathic Pain States"

Supplemental Figure 8 - related to Figure 5. Primary afferent input to inhibitory neurons, and spinal input to the Gr is unchanged by neuropathic injury

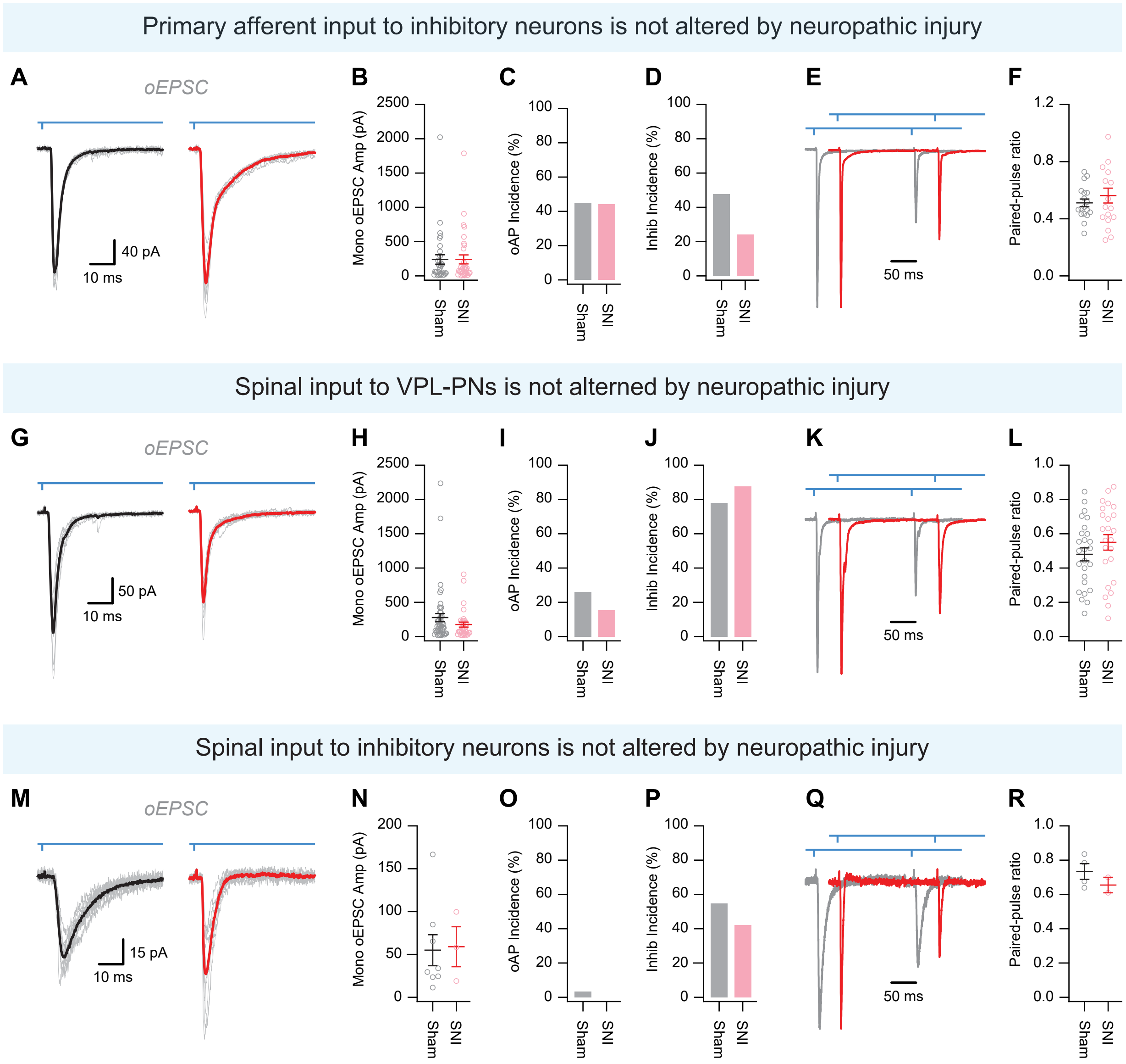
