## Supplemental Figure 9 for "The Dorsal Column Nuclei Scale Mechanical Sensitivity in Naive and Neuropathic Pain States"

Supplemental Figure 9 - related to Figure 5. Reduced inhibition onto VPL-PNs is not responsible for altered afferent signaling following neuropathic injury

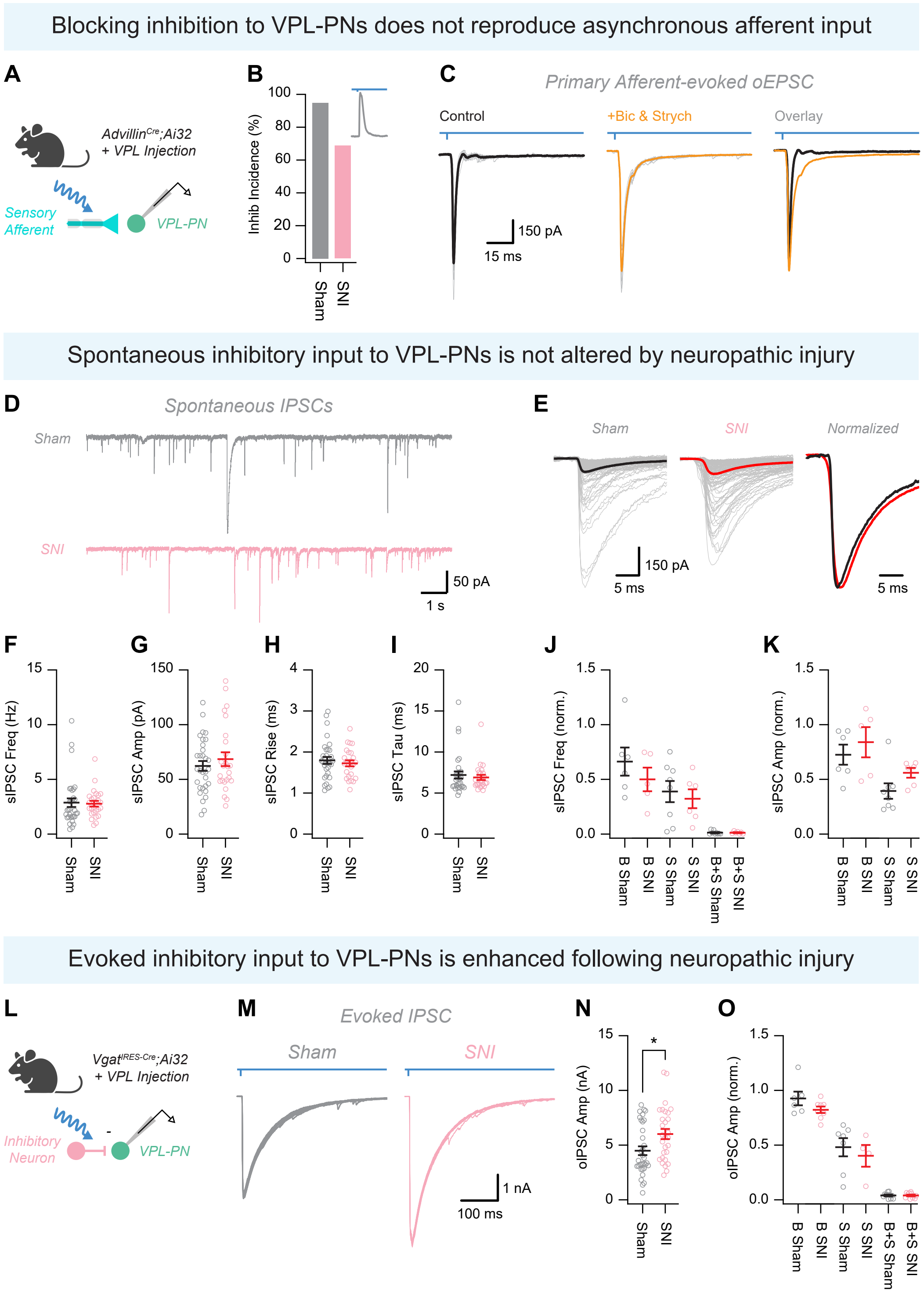
