## Supplemental Figure 10 for "The Dorsal Column Nuclei Scale Mechanical Sensitivity in Naive and Neuropathic Pain States"

Supplemental Figure 10 - related to Figure 6. SIlencing VPL-PNs or activating Gr inhibitory neurons does not affect noxious mechanical or thermal hyperalgesia during neuropathic pain

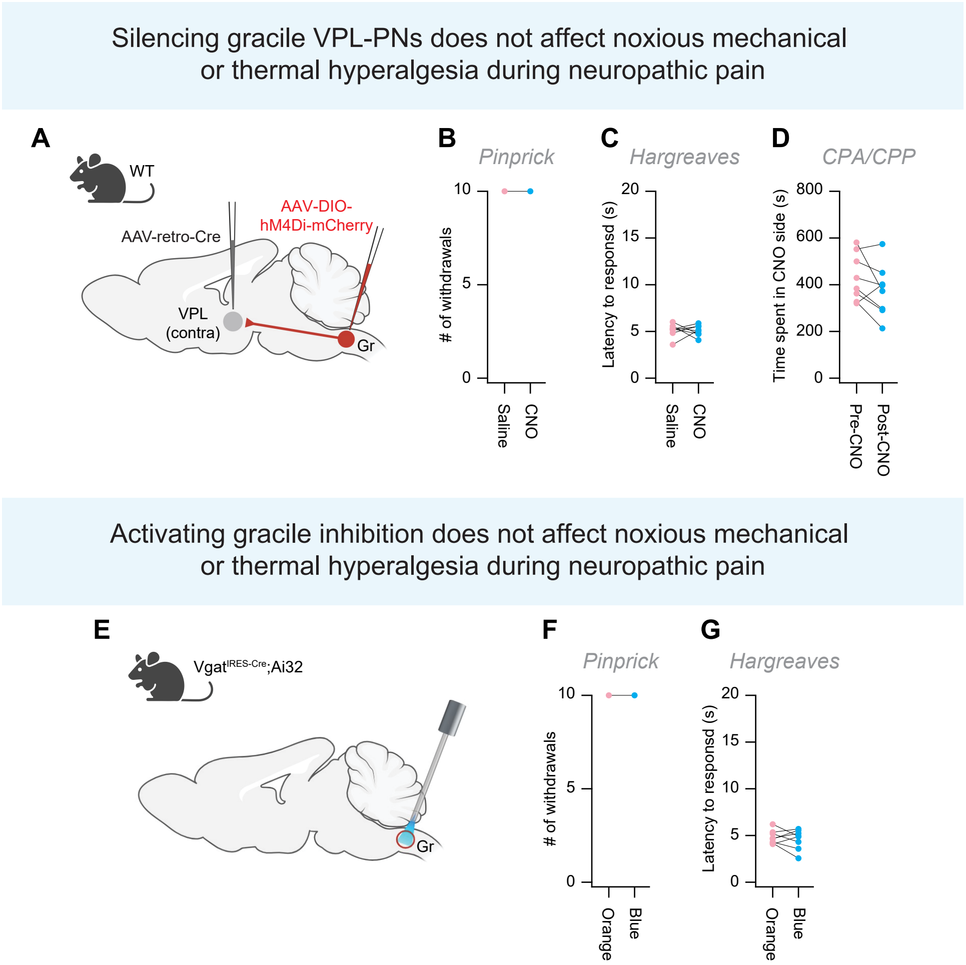
