## Supplemental Table 1 for "The Dorsal Column Nuclei Scale Mechanical Sensitivity in Naive and Neuropathic Pain States"

Supplemental Table 1- related to Figure 4. PAWS parameters for withdrawal responses of SNI mice compared to uninjured mice

| Withdrawal responses following SNI | | | |
|---|---|---|---|
| | Uninjured | SNI | p-value |
| 0.6 VF - Max height | 9.034 ± 0.6094 | 7.791 ± 0.5749 | n.s. 0.1346 |
| 0.6 VF - Distance traveled | 41.05 ± 7.87 | 87.7 ± 16.46 | ** 0.002 |
| 0.6 VF - Max y-velocity | 1318 ± 109.8 | 1367 ± 141.3 | n.s. 0.5457 |
| 0.6 VF - # of shakes | 2 ± 0.4024 | 3.75 ± 0.8241 | * 0.0317 |
| 0.6 VF - Guarding duration | 0.2729 ± 0.08339 | 0.5228 ± 0.07557 | * 0.0302 |
| Brush - Max height | 9.824 ± 0.5117 | 6.579 ± 0.5462 | *** 0.0002 |
| Brush - Distance traveled | 62.38 ± 13.76 | 193.9 ± 47.06 | * 0.0256 |
| Brush - Max y-velocity | 1415 ± 153.6 | 1191 ± 261.8 | n.s. 0.1514 |
| Brush - # of shakes | 2.143 ± 0.4671 | 4.5 ± 0.6124 | * 0.0269 |
| Brush - Guarding duration | 0.3206 ± 0.08725 | 0.5924 ± 0.07234 | * 0.0215 |
| Pinprick - Max height | 11.66 ± 1.591 | 6.125 ± 0.2533 | ** 0.0093 |
| Pinprick - Distance traveled | 109.1 ± 33.56 | 62.99 ± 12.71 | n.s. 0.4697 |
| Pinprick - Max y-velocity | 1526 ± 173.3 | 881.4 ± 138.2 | ** 0.0056 |
| Pinprick - # of shakes | 4 ± 0.6381 | 2.154 ± 0.4783 | n.s. 0.0593 |
| Pinprick - Guarding duration | 0.251 ± 0.06577 | 0.4721 ± 0.068 | ** 0.0078 |
