## Supplemental Table 2 for "The Dorsal Column Nuclei Scale Mechanical Sensitivity in Naive and Neuropathic Pain States"

Supplemental Table 2- related to Figure 4. PAWS parameters for withdrawal responses of mice during silencing of Gr inhibitory neurons

| Withdrawal responses while silencing Gr inhibitory neurons | | | |
|---|---|---|---|
| | Saline | CNO | p-value |
| 0.6 VF - Max height | 7.436 ± 0.6854 | 9.831 ± 0.8464 | * 0.0365 |
| 0.6 VF - Distance traveled | 39.9 ± 7.436 | 106.2 ± 17.65 | ** 0.0027 |
| 0.6 VF - Max y-velocity | 937.9 ± 106.1 | 1223 ± 120.1 | n.s. 0.0634 |
| 0.6 VF - # of shakes | 2.75 ± 0.5809 | 6.438 ± 0.8707 | ** 0.0052 |
| 0.6 VF - Guarding duration | 0.324 ± 0.07467 | 0.5586 ± 0.08703 | * 0.0386 |
| Brush - Max height | 8.873 ± 0.4813 | 9.73 ± 0.6737 | n.s. 0.281 |
| Brush - Distance traveled | 53.44 ± 6.784 | 85.06 ± 13.83 | * 0.025 |
| Brush - Max y-velocity | 1078 ± 102.7 | 1274 ± 84.14 | n.s. 0.1167 |
| Brush - # of shakes | 2.5 ± 0.4183 | 4.563 ± 0.689 | * 0.0154 |
| Brush - Guarding duration | 0.5063 ± 0.09655 | 0.3913 ± 0.07984 | n.s. 0.4332 |
| Pinprick - Max height | 10.43 ± 0.8766 | 9.853 ± 0.7378 | n.s. 0.5619 |
| Pinprick - Distance traveled | 81.41 ± 12.36 | 84.43 ± 14.29 | n.s. 0.886 |
| Pinprick - Max y-velocity | 1340 ± 101.9 | 1186 ± 107.1 | n.s. 0.234 |
| Pinprick - # of shakes | 4 ± 0.5845 | 4.125 ± 0.8056 | n.s. 0.7356 |
| Pinprick - Guarding duration | 0.319 ± 0.08456 | 0.3673 ± 0.09533 | n.s. 0.782 |
