## Supplemental Table 3 for "The Dorsal Column Nuclei Scale Mechanical Sensitivity in Naive and Neuropathic Pain States"

Supplemental Table 3- related to Figure 6. PAWS parameters for withdrawal responses of mice during activation of Gr inhibition

| Withdrawal responses while activating Gr inhibition | | | |
|---|---|---|---|
| | Orange | Blue | p-value |
| 0.16 VF - Max height | 6.118 + 0.6524 | 5.542 + 0.5771 | n.s. 0.6656 |
| 0.16 VF - Distance traveled | 91.65 + 23.95 | 31.62 + 4.574 | * 0.0342 |
| 0.16 VF - Max y-velocity | 501.3 + 54.81 | 321.3 + 54.63 | * 0.0469 |
| 0.16 VF - # of shakes | 8.125 + 2.039 | 2.333 + 0.7265 | * 0.0156 |
| 0.16 VF - Guarding duration | 0.4397 + 0.07671 | 0.4584 + 0.101 | n.s. 0.4653 |
| 0.6 VF - Max height | 6.456 + 0.6817 | 4.778 + 0.7544 | n.s. 0.0801 |
| 0.6 VF - Distance traveled | 60.89 + 11.88 | 31.03 + 7.247 | * 0.0195 |
| 0.6 VF - Max y-velocity | 454 + 57.08 | 317.4 + 47.3 | * 0.015 |
| 0.6 VF - # of shakes | 4.889 + 0.9044 | 3 + 0.7149 | n.s. 0.0754 |
| 0.6 VF - Guarding duration | 0.4938 + 0.08375 | 0.3986 + 0.09683 | n.s. 0.3633 |
| Brush - Max height | 6.682 + 0.424 | 5.132 + 0.5283 | * 0.0252 |
| Brush - Distance traveled | 57.85 + 8.991 | 28.92 + 4.963 | * 0.0489 |
| Brush - Max y-velocity | 489 + 65.27 | 279.4 + 53.91 | n.s. 0.0781 |
| Brush - # of shakes | 5.625 + 1.034 | 1.667 + 0.6667 | * 0.0391 |
| Brush - Guarding duration | 0.401 + 0.1159 | 0.5053 + 0.08079 | n.s. 0.5592 |
| Pinprick - Max height | 5.16 + 0.647 | 6.162 + 0.5958 | n.s. 0.2791 |
| Pinprick - Distance traveled | 59.5 + 14.73 | 47.75 + 6.845 | n.s. 0.7422 |
| Pinprick - Max y-velocity | 417.5 + 73.24 | 382.9 + 71.74 | n.s. 0.6406 |
| Pinprick - # of shakes | 5 + 0.9718 | 4.111 + 0.4231 | n.s. 0.5415 |
| Pinprick - Guarding duration | 0.4805 + 0.1262 | 0.4716 + 0.1097 | n.s. 0.8438 |
